# Characterization and classification of fine-resolution soil profile for precision agriculture using random forest and self-organizing map

**DOI:** 10.1101/2024.04.02.587707

**Authors:** Ani A. Elias, Megha Sharma, Shailendra Goel

**Affiliations:** Institute of Forest Genetics and Tree Breeding, Tamil Nadu, India; Department of Botany, University of Delhi, Delhi, India

**Keywords:** Fine-resolution soil map, soil prediction model, random forest, self-organizing map, marginal crops, precision agriculture

## Abstract

The availability of high throughput soil profile information is an important component in precision agriculture to perform efficient soil management for sustainable production. We collected 14 soil physiochemical features from Nagpur, Pune, and Haveri, representing target environments of safflower cultivation and also from our experiment station at Delhi, at fine resolution and created graphical maps to depict the variability. Additionally, we evaluated the predictive ability of two statistical learning models, random forest (RF) and self-organizing maps (SOM) against multinomial regression models for correctly classifying the soil profile. Clustering was performed around the medoids produced from the dissimilarity matrices of these models using partitioning around medoids (PAM) model. The robustness, versatility, and predictive ability of models in correctly classifying the soil profile to clusters were then tested using cross-validation which was repeated 100 times. This study was performed using training data with proportionate size varying from 60 to 95%, and increasing the unit area of observation up to nine times (or decreasing the total number of observations up to a ninth). RF model was found to be the best performing with average prediction accuracy above 85% in all settings which reached close to 100% in some settings. The predictive ability of all the models was maintained even when only the most influencing six variables were used for classification. The optimal training population size for prediction was found to be 70 – 80%. Based on our study, it is recommended to i) collect fine resolution edaphic features from a marginal farm before crop season, ii) use RF or SOM model to identify the most influencing features distinguishing the soil samples iii) expand the area of sample collection, find values for the most influencing features, and use RF model to correctly predict the class to which the new set of the soil belongs to.

**Graphical abstract:** 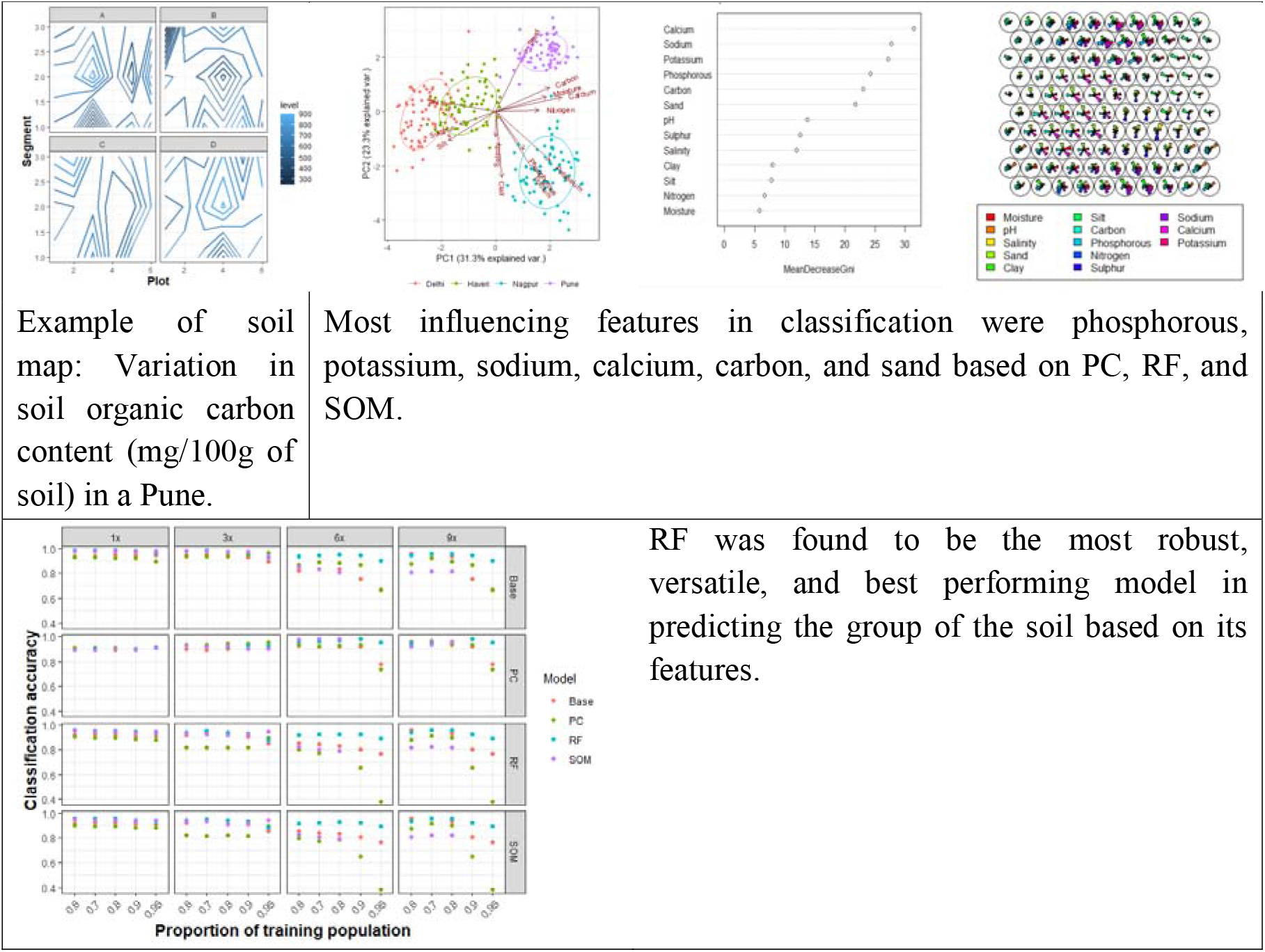

## 1. Introduction

Soil is one of the essential natural resources in a plant’s life. Soil delivers a wide variety of ecosystem services (Hannam & Boer, 2004) through soil functions which are linked to the physicochemical and biological properties of soil (Adhikari, K., & Hartemink, A. E., 2016). Overuse of chemical fertilizers in agricultural land is a main source of carbon emission and also directly threatens human, soil and aquatic health (Naghavi et al., 2022). Characterizing the soil through fine-resolution surveying and use of classification models for correctly grouping the land are important steps for identifying the appropriate soil management practices in precision agriculture (Reddy et al., 2021).

Due to the direct role on plant’s productivity and sustainability of the ecosystem, it is important to assess the soil properties and determine the soil quality. Soil texture and moisture are a couple of the most important physical properties affecting plant growth. Soil texture is determined by the proportion of mineral components of soil which includes sand, silt, and clay. Nutrients and water retention capacity of soil are controlled by texture. Aeration of the soil, root penetration, soil organic matter, nutrient dynamics, movement of water, heat conduction, and susceptibility to erosion are affected by the aggregation of textural components. Soil moisture influences this aggregation and leads to precipitation of the nutrients such as nitrogen for the plants which play a major role in its growth. A combination of different features including clay, soluble salts such as sodium, calcium, magnesium, and potassium lead to the salinity and electrical conductivity of the soil (McNeill, 1980). As the salinity increases, water availability decreases, and ultimately alter the plant growth. Soil organic carbon (SOC), a proxy of soil organic matter, also influences the nutrient retention capacity (Kumar et al., 2022) and is a key element in defining soil quality, fertility, and agricultural profitability. The presence of salts, SOC, and pH influence the chemical property of the soil. The most used indicator for nutrients are soil chemical indicators and for this reason, they are usually referred to as ‘indices of nutrient supply’ (Powers et al., 1998). Characterizing the physical and chemical properties of the soil helps in identifying the requirements for precision agricultural management.

Monitoring spatial changes in the soil quality and understanding its dynamic nature, and role in precision agriculture and land management are crucial. The construction of accurate maps based on the spatial variability in physio-chemical soil properties is the important first step in this process (Behera et al., 2016). Detailed maps based on high-resolution sampling can act as stepping stones for many conservations policies at the national and international levels. Various attempts have been made to construct soil maps using digital and conventional methods at the global level (Arrouays et al., 2020; Grunwald, 2009; Hengl et al., 2014, 2015; Hengl, Leenaars, et al., 2017; Poggio et al., 2021) national-level (Purushothaman et al., 2022; Reddy et al., 2021) and at the local level (Arora et al., 2021; Dharumarajan et al., 2020, 2021, 2022; Kalambukattu et al., 2018; Kaushal et al., 2021; Santra et al., 2017; Srinivasan et al., 2022). However, most of the maps are at low resolution levels. With Global Soil Map project (Arrouays et al., 2014) aimed for fine-resolution mapping using digital soil mapping (DSM) approaches, there is an exponential growth in high-resolution (>10,000 km^2^) soil maps in the last two decades (Chen et al., 2022).

Creation of the soil maps requires sampling and measurement of features of interest, followed by statistical analysis involving interpolations, simple regressions, or the use of the artificial intelligence models to process the patterns from the datasets. With fine resolution sampling, more information can be extracted with greater confidence for more useful analysis and interpretation. Such complex data provide better inference. Linear and non-linear statistical models can be used to evaluate spatial variability in the soil from complex datasets (Cavigelli et al., 2005). Principal component analysis (PCA) has been widely used in soil analysis (Collins & Ovalles, 1988; Fox & Metla, 2005; Ladoni et al., 2010; Levi & Rasmussen, 2014; Sena et al., 2002), as it can correlate several variables simultaneously (Sena et al., 2002). Also, PCA can handle high-dimensional data arrays, for e.g., three-way PCA can be useful to analyze many soil variables obtained from the multiple locations as a function of other experimental conditions such as sampling time (Henrion, 1993). Multivariate linear regression (MLR) and correlation are other most commonly used methods (Besalatpour et al., 2013; Elaoud et al., 2017; Zornoza et al., 2007), to infer the relationship among soil features. MLR has a simple structure, easy calculation, and interpretation, however, it is inadequate to detect non-linear relationships (Fong et al., 2002). Machine learning (ML) and deep learning (DL) approaches have been used to conclude linear and non-linear relation between the soil features and their distribution (Padarian et al., 2020). The most popular ML and DL tools for the estimation of the soil features, its classification, and prediction are random forest (RF) and artificial neural network (ANN) (Gopal & Bhargavi, 2019; Xie et al., 2021). RF is a classification machine learning approach with the ability to model complex and nonlinear relationships (Brokamp et al., 2017). Besides this, its capacity for determination of the relevant variables, resistance to over-fitting, robustness, and establishment of an impartial measure of error rate (Breiman, 2001; da Silva Chagas et al., 2016; R. Zhang et al., 2016), are the advantages which made RF most widely applied approach for the analysis of soil variables (Gambill et al., 2016; Heung et al., 2014; Hounkpatin et al., 2018; Ließ et al., 2012; Wiesmeier et al., 2011; Y. Zhang et al., 2019). ANN, analog to the biological neural network, is more complex and uses synaptic weights to establish a connection among predictor variables through multiple layers of networks and predicts the classification (Grunwald, 2022). Some of the features of the neural network such as no assumption of a prior relationship, availability of multiple training algorithm detections of complex nonlinear relationships, and detection of all possible interactions between predictor variables (Tu, 1996), made it another popular soil classification model (Alvarez et al., 2011; Ayoubi & Karchegani, 2012; Baligh et al., 2020; Hossein Alavi et al., 2010; Koekkoek & Booltink, 1999; Minasny et al., 2004, 2016; Ozturk et al., 2011). Application of multiple models for the classification of soil can be useful in arriving at more robust prediction as well providing a comparison among different models.

India exhibits high soil diversity. Although alluvial is the most prominent, red, laterite, black, and brown forest, mountain meadow are the other soil types found in India (https://www.soilmanagementindia.com/). In these different soil types, nine major types of oilseed crops are grown which include sunflower and safflower. Safflower (*Carthamus tinctorius* L.) is an oilseed crop belonging to the family Asteraceae. It is a minor seed crop and produces a healthy oil rich in unsaturated fatty acids. India produces 2640 tons of safflower in its major soil type, the black soil (<u>Directorate of Economics And Statistics,Ministry Of Agriculture,Government Of India (dacnet.nic.in))</u>. However, safflower can grow in other soil types producing varying phenotypes. In this study, we aim to (i) create soil maps, (ii) classify soil samples collected at fine-resolution from target environments (TE) of safflower based on their physicochemical features, and (iii) evaluate variability in soil features and develop soil prediction models for safflower cultivation. To our knowledge, this is the first type of prediction model study in the arena of precision agriculture of safflower focusing on a large set of physicochemical features for fine-resolution mapping in India.

## 2. Material and methods

### 2.1. Experiment design

As a part of the safflower crop improvement program, we conducted a multi-environmental trial for a safflower breeding population (recombinant inbred lines) at different locations in India. The locations are representative of the TE or suitable environment for safflower cultivation. Soil samples were collected from these locations for soil profiling. Four soil profiles representing alluvial (Delhi University (DU) Research Field, Delhi, 28°74′ N, and 77°12′ E), black (Nimbkar Agriculture Research Institute (NARI), Pune, 18°57′ N, and 74°05′ E), red and yellow (Ankur seeds (AS) Pvt. Ltd., Nagpur, 21°13′ N, and 79°09′ E), and laterite (a farmer’s field in Haveri, 14°78′ N and 75°39′ E) (Fig S1) were collected. In this fine-resolution soil profile study, a total of 288 samples (72 samples × 4 locations) were collected for evaluating the micro and macro variations. Each location had four blocks labelled A, B, C, and D placed as far as possible from each other in the given area. The distance between blocks was the furthest in the DU research field where they were kept in a ∼40,000 m^2^ land. In other locations, the blocks were placed logistically in ∼2,000 m^2^ land.

In general, a block was of dimension 20 m × 17 m which was partitioned into six rectangular plots each of dimension 20 m × 2 m separated by a gap of 1 m along the width. This design was originally created for safflower cultivation. For soil collection, the plots were again partitioned into segments of ∼7 m × 2 m dimension. To understand the micro-variation in the physiochemical characteristics of the soil within the location, the samples were collected from these segments in a plot. From each location, 72 samples (18 samples × 4 blocks) were collected from pits of depth 30 cm by mixing soils from different layers along the depth. Soil collection was done from five random pits in a segment and later homogenized to make one composite. Samples were carried to a soil laboratory at Delhi University from all sites in airtight zip lock bags for the physiochemical analysis. The samples were dried, sieved through a 2mm sieve and stored at room temperature for analysis. Soil was collected before the start of crop season to make sure that any management or nutrient application not biasing the soil features.

### 2.2. Physiochemical analysis of soil

Soil features including moisture content, presence of micro and macro nutrients such as nitrogen, phosphorous, potassium, calcium, sodium, sulphur, and organic carbon, texture such as percentage of sand, clay, and silt, salinity, electrical conductivity, and pH were measured in the soil science laboratory of the University of Delhi and Indian Agricultural Research Institute, Delhi. To determine the soil moisture percentage (gravimetric water content (GWC)), 10g of fresh soil was taken in petri dish and oven dried at 105°c until constant weight was achieved. Estimation of GWC percentage was done as per the protocol by Allen et al. (1974). The GWC (%) was measured as follows:

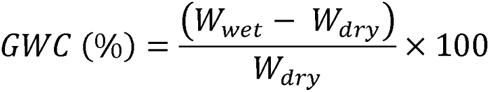

Where *W_wet_* and *W_dry_* are the weights of wet and dry soil in grams respectively.

A soil suspension was prepared using 20 g of soil and 40 ml of deionised water (1:2 soil to water ratio). Homogenisation of soil sample was done by stirring it on a magnetic stirrer for one hour and later filtered through Whatman-1 filter paper. Electrical conductivity (*µs/cm*), pH, and salinity (ppm) of the supernatant were measured using Multiparameter Tester (PCS Tester 35). Soil texture (%) was analysed using the Bouyoucos Hydrometer method (Motsara, 2015). Nutrients were measured in the unit mg/100g of soil. Available K, sodium (Na), and Ca were extracted using neutral ammonium acetate and estimated with flame photometric method (Allen et al., 1974). Total available N was estimated with automatic micro-kjeldahl distillation unit (Gallaher, R. N., Weldon, C. O., & Boswell, F. C., 1976). Total organic C was estimated using the wet oxidation method (Chan et al, 2001). Turbidimetric method was used for the estimation of available S(Williams & Steinbergs, 1959). The available P was estimated using Olsen’s method (Sims, 2000)

### 2.3 Statistical analysis

#### 2.3.1 Background

Average value from three replicates was used for analysis except for soil texture, where only one observation was taken. Missing values were imputed as average values from nearest neighbour segments. A correlation analysis was performed and removed EC having high collinearity with salinity (Fig 1). Thirteen soil features were chosen for further analysis. Graphical soil maps were created showing the variability of these features within blocks in a location (Fig S2).

**Figure 1.**
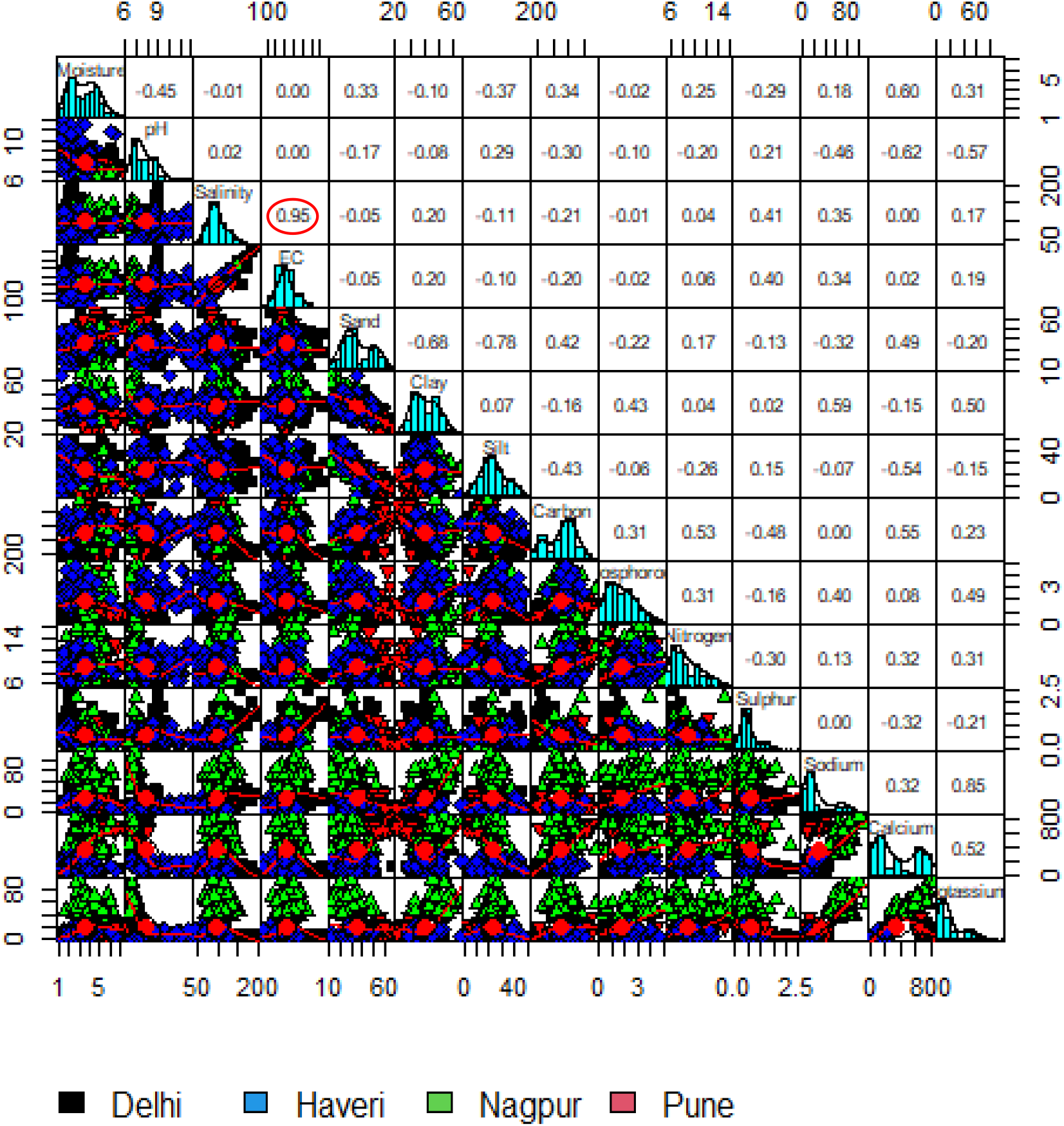
: : Correlation and variability of all soil features collected from the field. The value with red circle is considered to have high correlation. The electrical conductivity (EC) and salinity follow the same pattern of correlation with other variables also. To avoid multicollinearity, variable EC is removed from further analysis.

The general model is

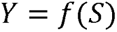

Where Y is the classification labels based on soil features and S is the set of soil features. This generalized model was evaluated using four main model frameworks for narrative and predictive abilities. The features were standardized before modeling.

Four models were used for classification and prediction: Multinomial model using i) all the variables hereafter called as base model ii) principal component analysis (PCA), a dimension reduction method; the first two principal components (PCs) which explained >50% of the variation were then used in the model hereafter called as PC model iii) RF, a classification ML model, and iv) self-organizing maps (SOM), an ANN model. The models were tested for classification prediction for locations using 70:30 training: test datasets cross-validation method which was repeated 100 times. The accuracy in classification and confusion matrix were used as criteria for predictability evaluation. Later, unsupervised models were used for classification of the data based on the similarities of the soil features to overcome the limitation of grouping based on locations. The similarity matrices from the models were used to cluster the soil profile using a k-medoid algorithm. The predictability of all models in correctly classifying the soil profile into clusters were evaluated using proportionately partitioned training: test datasets cross-validation method which was repeated 100 times. Average classification accuracy was used for predictability evaluation.

#### 2.3.2 Models

##### 2.3.2.1 Multinomial regression models

The model used is as follows:

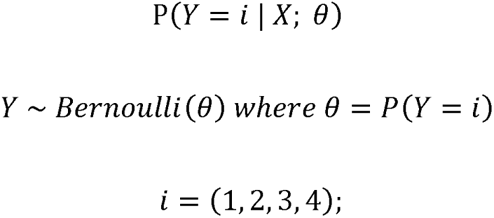

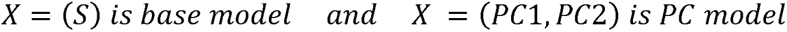

The first two PCs which explained >50% of the variation was used in the PC multinomial model.

##### 2.3.2.2 Random Forest model

Random Forest (RF) (Breiman, 2001) is an ensemble classifier machine learning approach growing many classification trees (Louppe, 2014). Each tree gives a classification giving ‘votes’. The forest choses the classification having the most votes over all the trees. The random forest uses bootstrap aggregating (bagging) algorithms for an internal cross-validation for getting unbiased estimate of the classification error. The bagging process is repeated ‘B’ times.

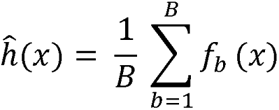

The out of bag (oob), *x*, would be fitted with the classification function *f_b_* and then all the results from the individual trees are averaged.

Gini impurity measure, *G,* is used as the classification function in RF. The best classification at a split (node) in a tree is made based on the probability of using a set of variables *p*, the value of *p* is held constant during the tree growth. *G* for the descendent nodes is less than the parent node. Adding up *G* for each variable set *p* over all the trees in the RF provides criterion for ranking variables based on importance. The Gini impurity measure for a tree, *G*(*p_t_*) is

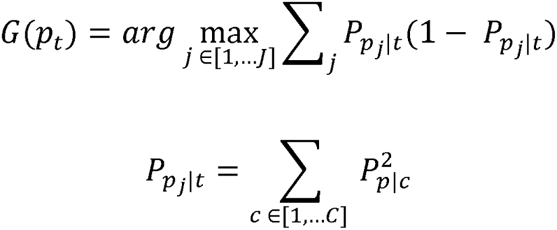

Where *P* is the probability of picking *p* set of variables; *c* is the class; *j* is the node; *t* is the tree.

A similarity matrix is computed between the observations with classification of oob observation. If two observations are classified in the same tree terminal node, the proximity between them is incremented. One thousand trees are created in the forest using pairwise comparison based on G.

##### 2.3.2.3 Self-organizing map

Self-organizing map (SOM) (Kohonen, 1998) is an ANN that does dimension reduction as well as clustering simultaneously by keeping similar objects together (Wehrens & Kruisselbrink, 2018). The original ‘online’ algorithm of SOM is used in this study. In the algorithm, the data is described by numerical vectors belonging to *K* units on a two-dimensional grid, here in a Euclidean space (ℝ^*p*^). A codebook vector (neuron) *m_k_* ɛ ℝ^*p*^ is attached to each unit *k*, the initial value of the vector is chosen based on the distance function.

A random data is mapped to a trained SOM by calculating the distance of the datapoints to the codebook vectors and assigning each datapoint, *x*, to the unit with the most similar codebook vector called as the best matching unit. The matching units are assigned to clusters. The best matching unit in a cluster *c* at a time *t* is defined by minimizing the distance, here Euclidean distance, between *x* and matching codebook *m_k_* as

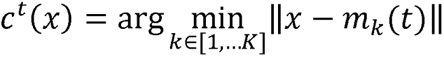

And codebook vectors are updated via

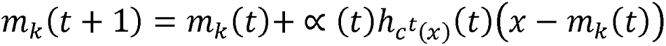

Where ∝ (*t*) is the learning rate; *h*_C^t^(*x*)_(*t*) is the neighbour function, here Gaussian function, at time *t* between winner unit, *j*, and the neighbour unit, *i*.

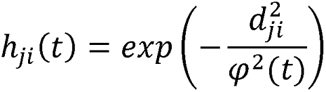

where *d_ji_* is the Euclidean distance; *φ* is the standardization parameter which decrease over time *t* to reduce the intensity and scope of the neighbourhood relations. One thousand iterations were conducted for the model.

#### 2.3.3 Clustering of observations

The observations were grouped into clusters based on dissimilarity using a k-medoid algorithm. The k-medoid clustering is a robust clustering method (P. Arora et al., 2016) and is less sensitive to outliers as it uses representative observations called medoids for clustering. Partitioning around medoids (PAM) (Kaufman & Rousseeuw, 1990) is known as the most powerful k-medoid clustering method (Park & Jun, 2009) and is used in this study. PAM was performed on Euclidean distance matrices obtained from i) normalized data which is used in the base ii) dimension reduced data using PCA where only first two PCs are used iii) proximity matrix from unsupervised RF and iv) unified distance matrix (U-matrix) from unsupervised SOM. The Euclidean distance matrix, *d_ab_*, from two datapoints *a* and *b* is calculated as:

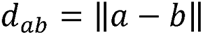

The distance matrix from proximity matrix, *s* of RF is calculated as

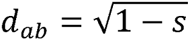

For each randomly selected *k* medoids, PAM algorithm selects datapoints that decrease the distance within a medoid. The total within-cluster sum of square (wss) is calculated for each datapoint *x* for a medoid *k* of a cluster *c* as follows:

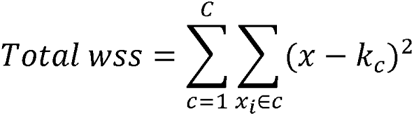

Total wss measures the ‘goodness’ of the cluster, the lower the value the better the cluster. Optimal number of clusters was selected using elbow method by plotting the total wss to the number of clusters and choosing number corresponding to the bend (elbow) of this line graph. Best model for partitioning the data was chosen by comparing the wss among the optimal clusters from all the four methods.

#### 2.3.4 Predictability testing

The predictability of the classification models was tested using cross-validation (CV). The predictability was tested using ‘location’ or ‘cluster’ as the response variable using CV method where the datum was randomly partitioning into training and test datasets. A confusion matrix was used to represent the counts of actual vs predicted classification from each iteration of CV. Later, average values were calculated from all the confusion matrices. The accuracy in classification was calculated for each confusion matrix and averaged as follows:

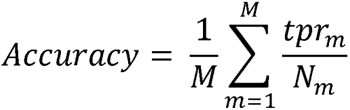

Where *M* is the total number of confusion matrices from a CV; *tpr* and *N* are the counts of truly predicted classes and total counts respectively from a confusion matrix, *m*.

#### 2.3.5 Impact of training population proportion and versatility of model

Various instances of models were performed to check the robustness and versatility using different proportions of training population in predicting the clusters based on soil features. Clusters created using a specific model was tested using all the models to check the predictability of the models irrespective of the origin of clustering. Predictability was also evaluated with varying training population size which was set as low as 60% to as high as 95%. Also, the models were evaluated by increasing the area of observation from three to nine times. In such instances, observations from segments (area of 1x) were averaged together to produce a single value for an area thus decreasing the number of total observations while increasing the area. For example, a soil feature value for a 3x area was calculated by averaging the values of the feature from three logistically adjacent segments in that area thus reducing the total number of observations to a third. The most important variables influencing the classification were filtered out based on PCA, RF, SOM. Similar instances as mentioned above were performed based on these fewer number of variables. Evaluating the models for reduced number of variables and observations were practically and economically important. The CV in these simulation settings was repeated 100 times and the average accuracy in correctly predicting the classes were observed.

#### 2.3.6 Software used

The scripts to perform statistical analysis was written in R language (R v4.2.2). Packages ‘nnet’ (Venables et al., 2002), ‘randomForest’, and ‘kohonen’ were used for running multinomial regression, RF, and SOM models respectively. PAM clustering was performed using the package ‘pmclust’.

## 3. Results

### 3.1 Using original data and settings

Principal component analysis (PCA) was performed for all the variables. The first two PCs explaining 54% of the variation (Fig S3) was chosen for further analysis. The biplot (Fig 2) indicated that the soil from Pune and Nagpur are clearly distinguishable from each other and also from those in Delhi and Haveri. Delhi and Haveri soils have similar features with Haveri being more alkaline in nature. Soil from Pune is sandier with lower clay content. Soil from Delhi and Haveri are siltier in nature. Both Nagpur and Pune soils have high calcium and carbon content while Nagpur soil has more sodium.

**Figure 2:**
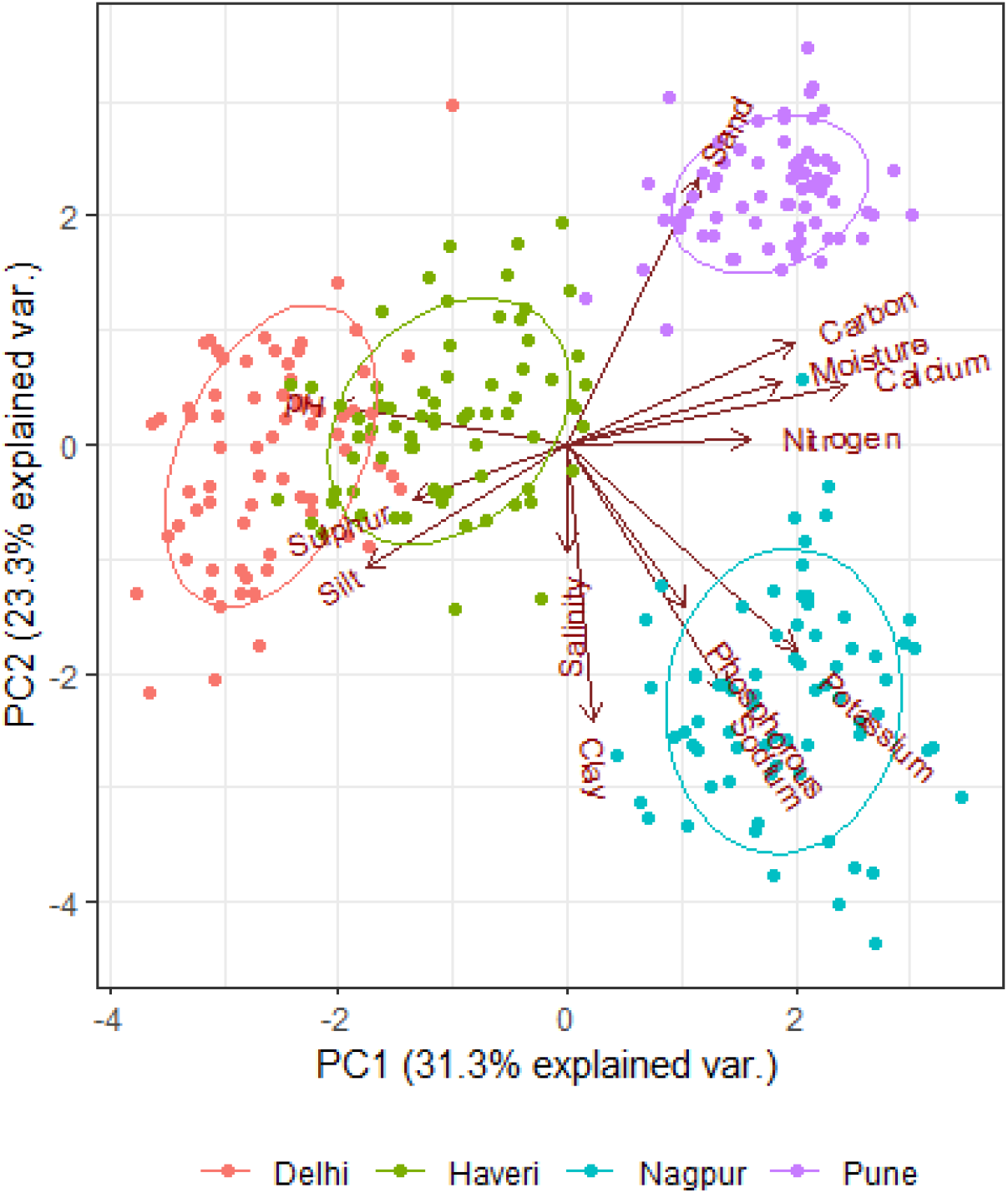
Classification of soil based on PC 1 and 2. The further the loadings (vectors) are away from the origin, the more their influence on the PC. PC1 was predominantly influenced by calcium, carbon, and pH while PC2 by sand, clay, and sodium. Both PC 1 and 2 were similarly influenced by potassium. Pune and Nagpur soil were observed to be clearly distinguishable from each other and those in Delhi and Haveri. There were similar features between Delhi and Haveri soils.

PC values were obtained and used as predictors in a multinomial model. The base model provided an average accuracy of 0.99 (Fig 3 A and B) which could be due to the use of all the variables available. Because of the overlapping features of Delhi and Haveri soil, there are misclassification in prediction of those on an average of 6% (Fig 3C) from PC model which resulted in an average prediction accuracy of 0.92 (Fig 3D). Clustering was performed using PAM and five or four clusters were selected as optimum for base and PC respectively (Fig 4 A-D). The clustering indicated that some segments of a location have features similar to those in other locations.

**Figure 3:**
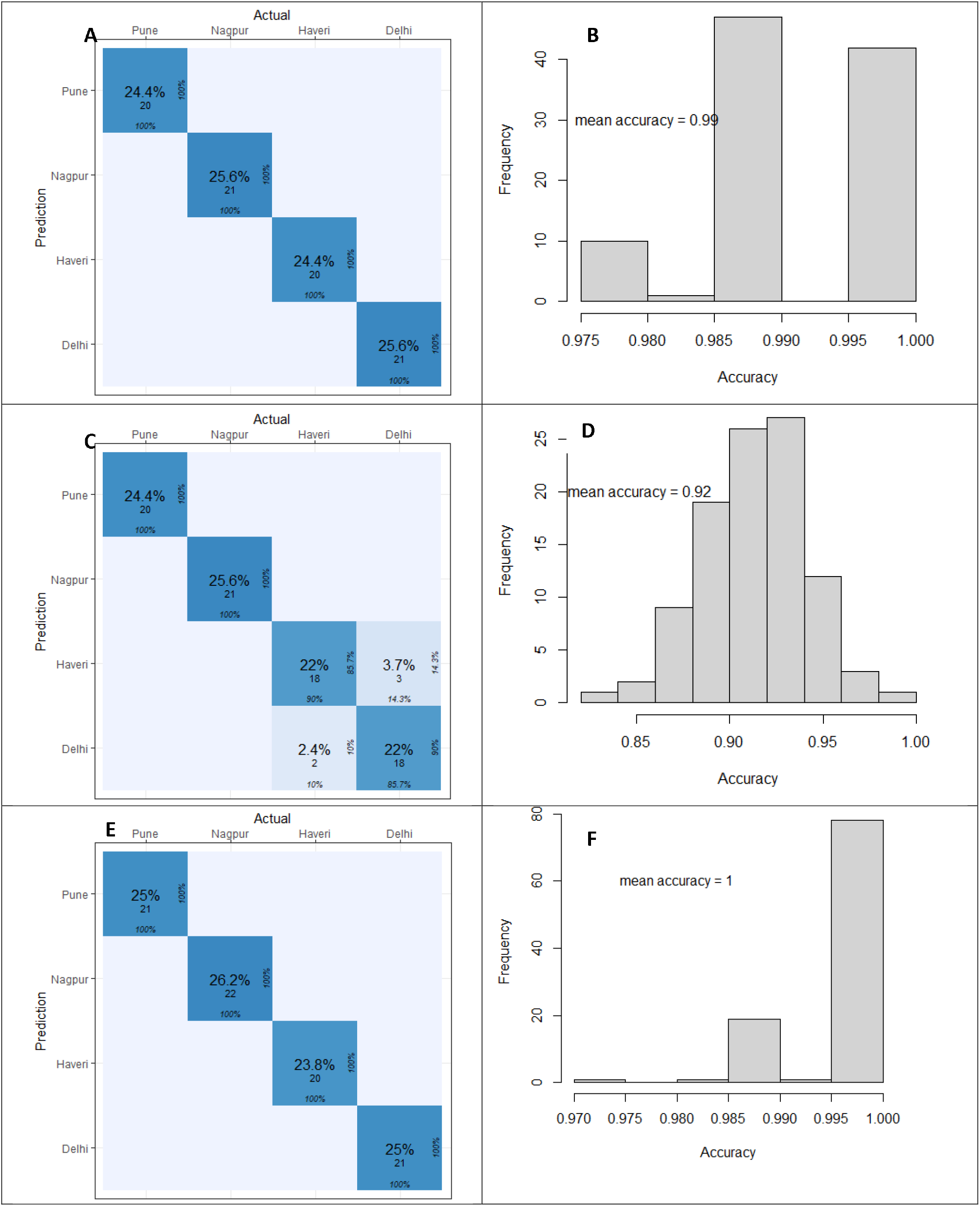
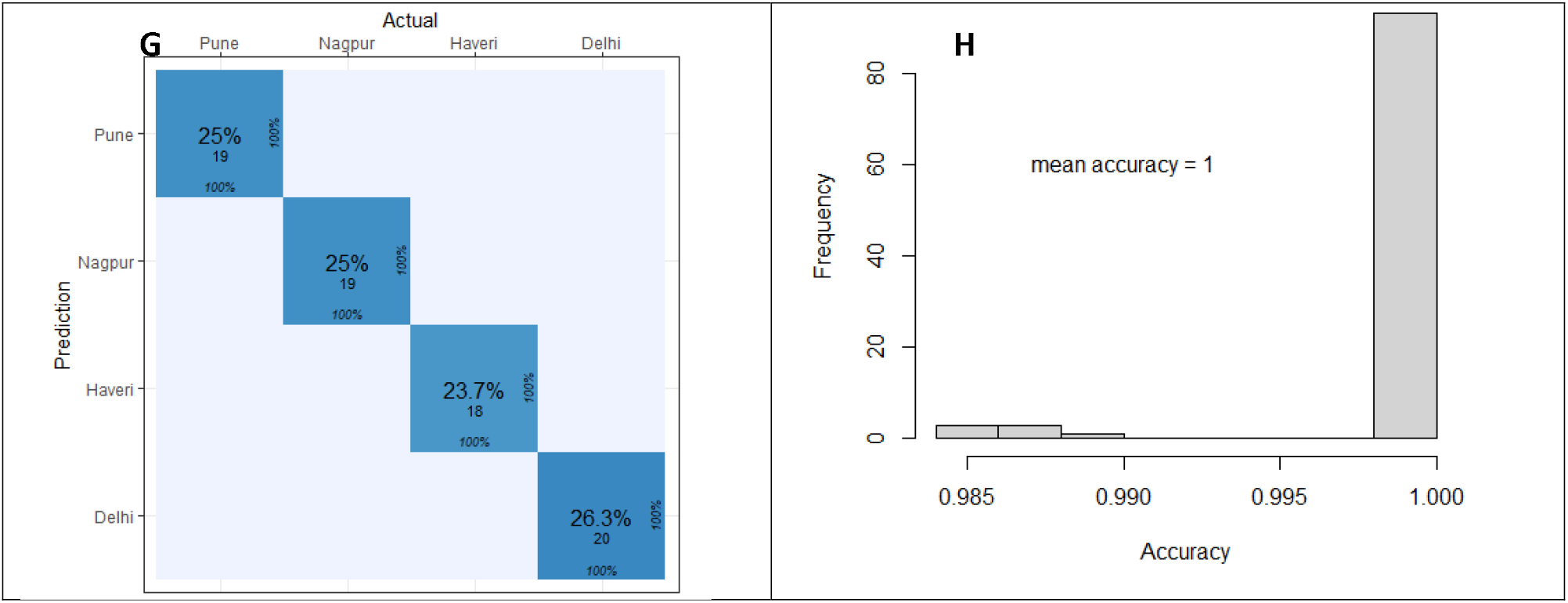
Results obtained from 100 replications of 70:30 training: test cross-validation predicting location. Left hand column represents average confusion matrices and right-hand column represents histogram of accuracy. A and B are results from base multinomial model, C and D from multinomial model using PC1 and 2, E and F from random forest model, and G and H from self-organized maps.

**Figure 4:**
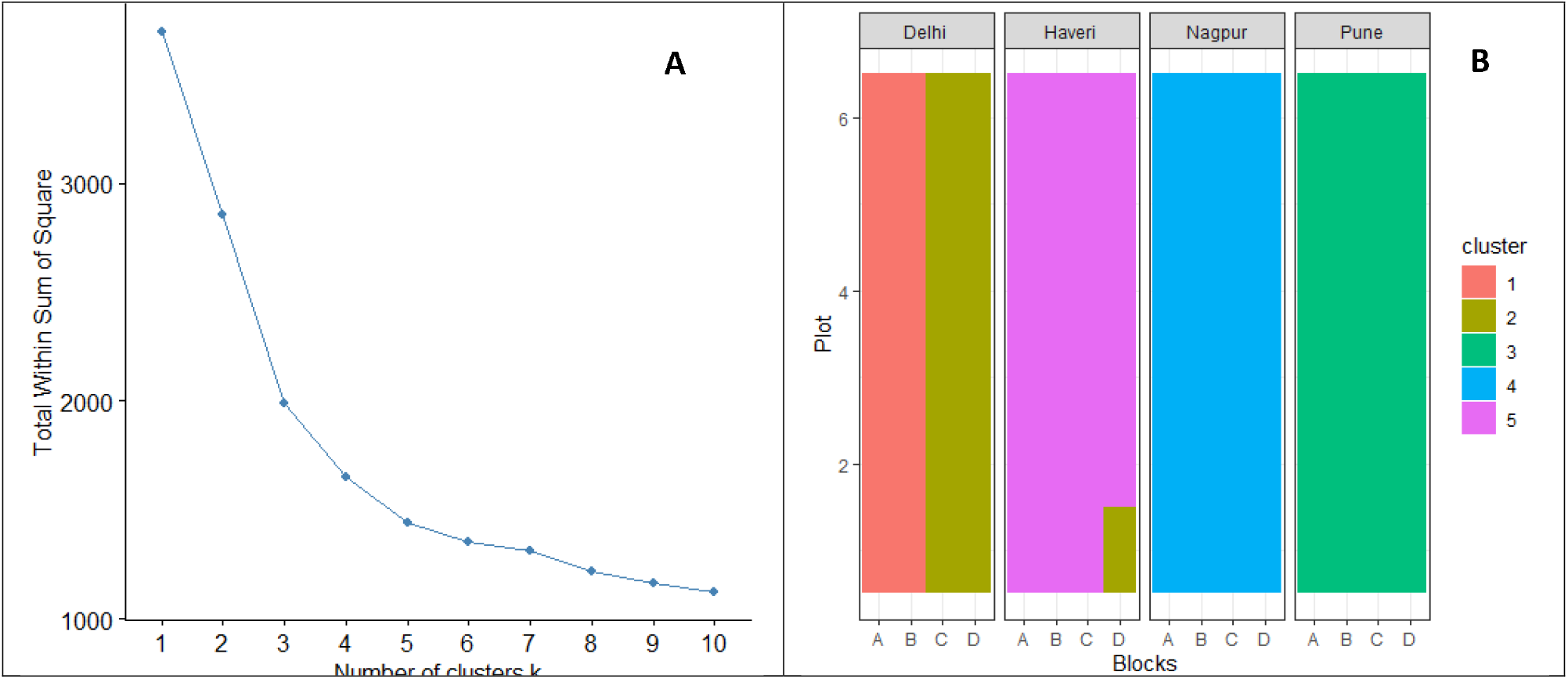
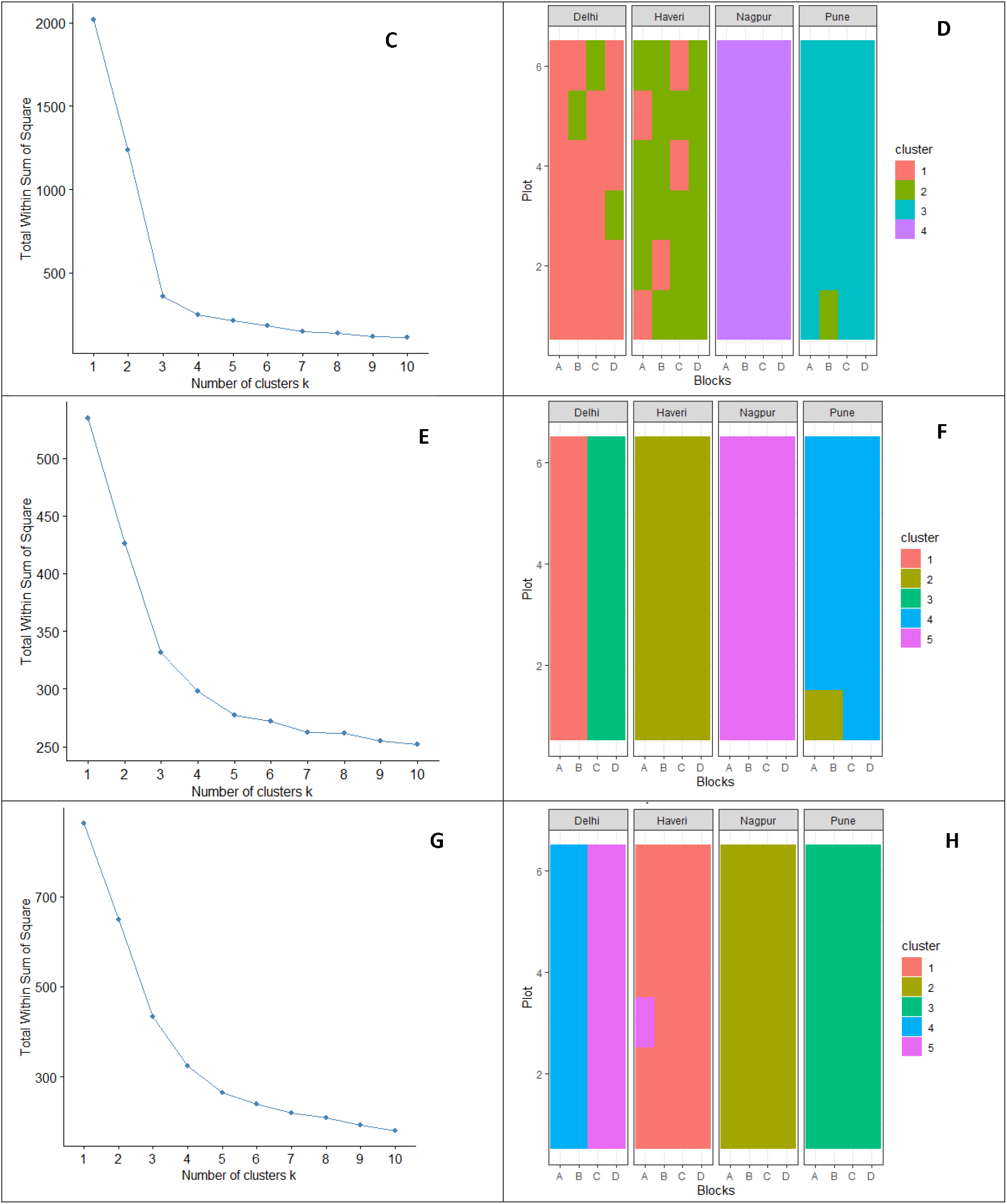
Optimum number of clusters of soil types based on the within sum of squares (wss) using the similarity measurements for each model. Left hand column indicates the wss for all the possible clusters and right-hand column indicates the representation of fields using optimum number of clusters. A) and B) based on all features after nomalizing C) and D) based on PC1 and 2 E) and F) based on the distance matrix from unsupervised random forest, and G) and H) from unsupervised self-organizing maps.

Random forest model was performed with locations as the response variable. Using random forest, the locations were predicted with no misclassification error on an average (Fig 3E and F) with a mean prediction accuracy of 1.00 (Fig 3F). Five clusters optimally distinguished the locations as well as blocks within Delhi (Fig 4 E-F). Calcium content is found to be the most important feature distinguishing the locations (Fig 5A) as well as the clusters (Fig 5B).

**Figure 5:**
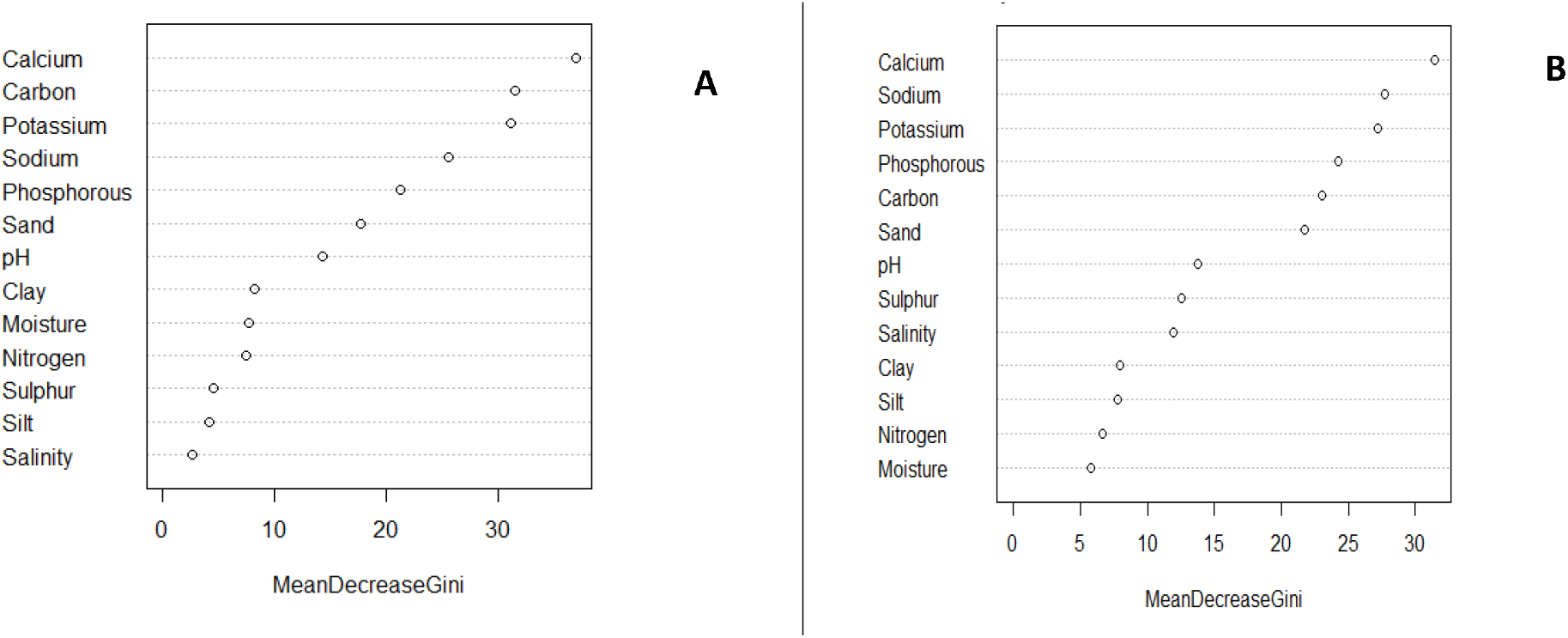
Variables of importance based on mean decrease in Gini value from the supervised random forest model using A) locations as response variable B) cluster as response variable. Calcium content was found to be most important in distinguishing locations and clusters.

Self-organizing maps were used to evaluate the clustering based on soil features and ability to predict location. On an average, there was no misclassification error and the predictability was 1.00 (Fig 3 G-H). An unsupervised SOM was used for identifying the most influential variables from the codebook vector (Fig 6A). A U-matrix (Fig 6B) was used for visualizing the distances between codebook vectors and clustering those (Fig 6C). The model converged around 600 iterations (Fig S4). The re-clustering was done using PAM on U-matrix. Five clusters were chosen as optimum based on the wss (Fig 4 G – H).

**Figure 6:**
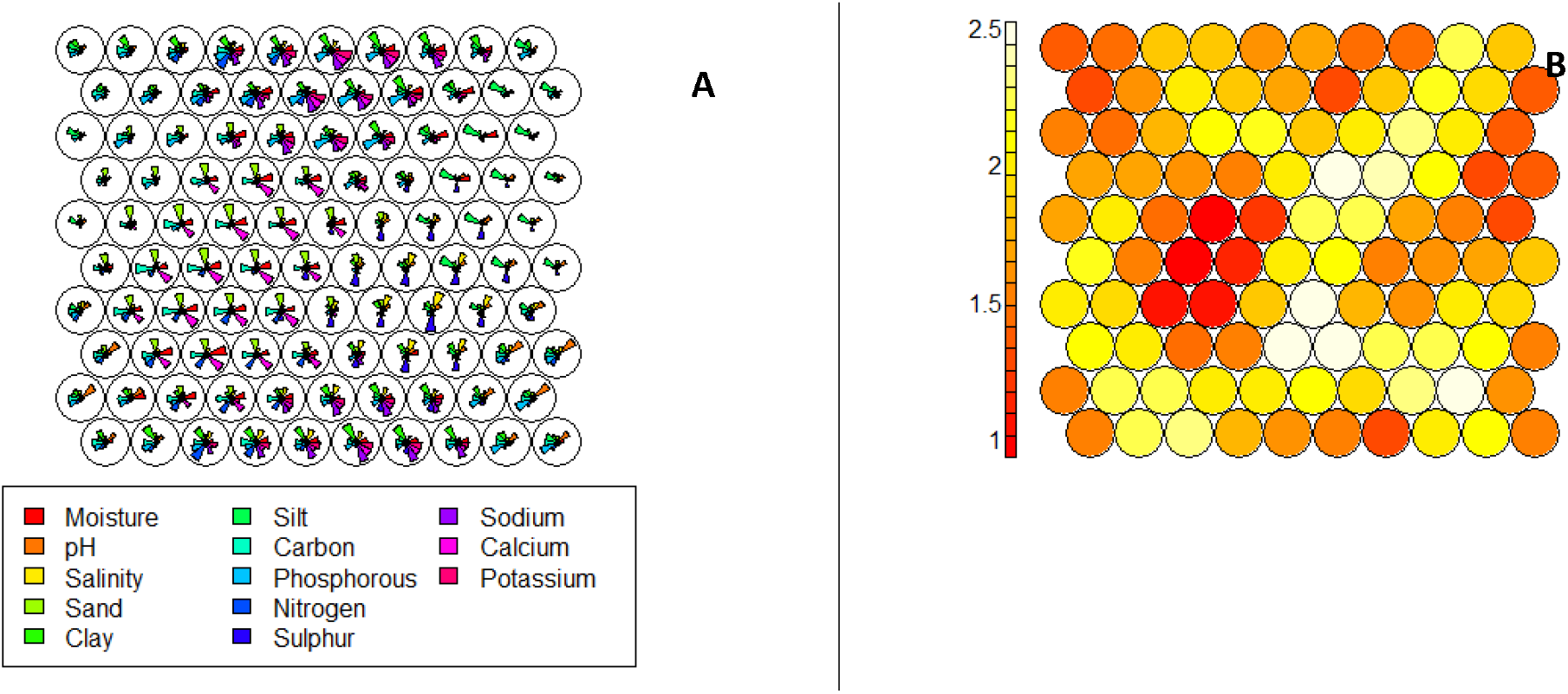
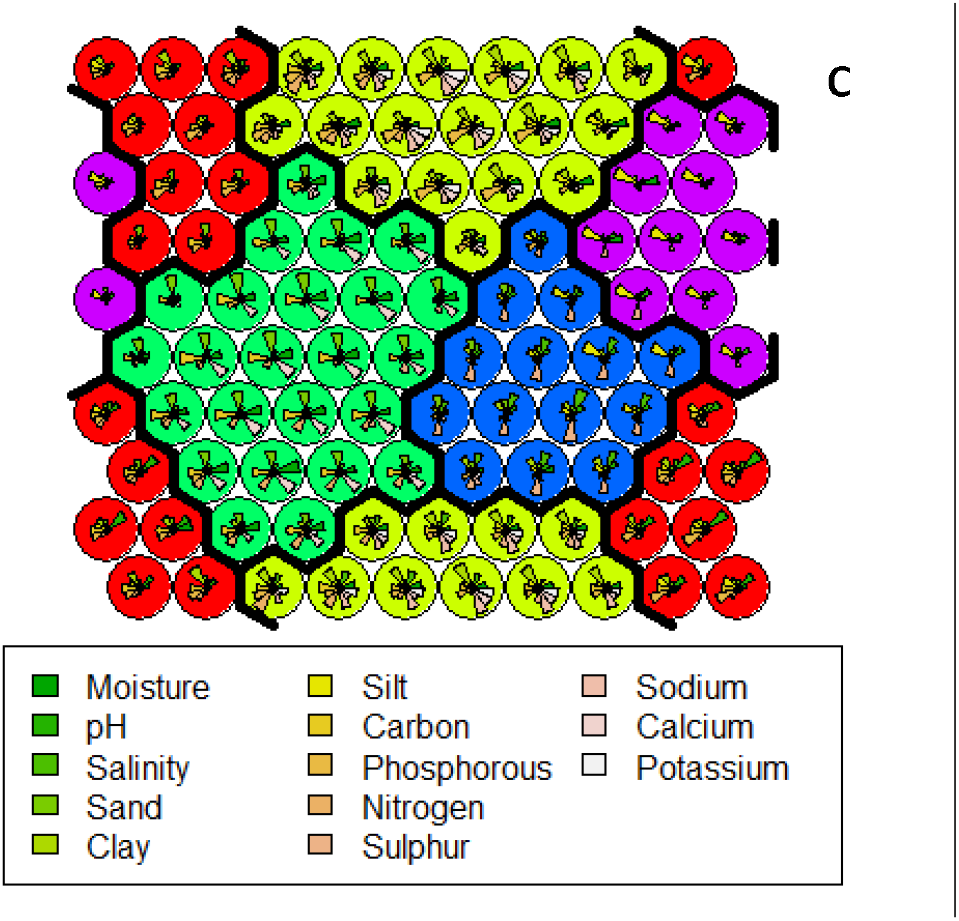
Result from unsupervised SOM. A) neighbourhood codes indicating most influential variables B) U-matrix between neurons where light colors signify the closeness of codebook vectors while the dark colors correspond to the gap between vectors C) cluster map based on optimal number of clusters (five).

Based on the wss values and optimal number of clusters, RF and SOM were considered as best models for clustering. They both produced clusters that can be logistically applicable. The clustering based on base also appeared similar to those from RF and SOM but with a higher wss. The macro nutrients P and K, the micro nutrients Na and Ca, C, and percentage of sand were found to be the most influential variables in classifying the soil profiles used in this study (Fig 2, 5, and 6A).

### 3.2. Versatility, robustness, and predictability of models

Overall, RF was found to be the most robust, versatile, and best performing model with accuracy never below 85% irrespective of the training population size, number of features used, total number of observations in the testing population, and the type of cluster (Fig 7 A and B). The prediction accuracy of RF was never below 90% when only the most influencing features were used in the model. RF and SOM performed similarly except when the area of measurement was increased. Since the total area and number of collected observations are constant in this study, on increasing the unit area per observation (for example, from 1x to 3x), the total number of observations reduces proportionately. Additionally, the prediction accuracy of RF and SOM were close to 100% when clusters from base model were evaluated. SOM failed to perform when the number of observations in the testing population was less than five. Out of the two multinomial regression models, base model performed better, even with increase in the proportion of training population.

**Figure 7:**
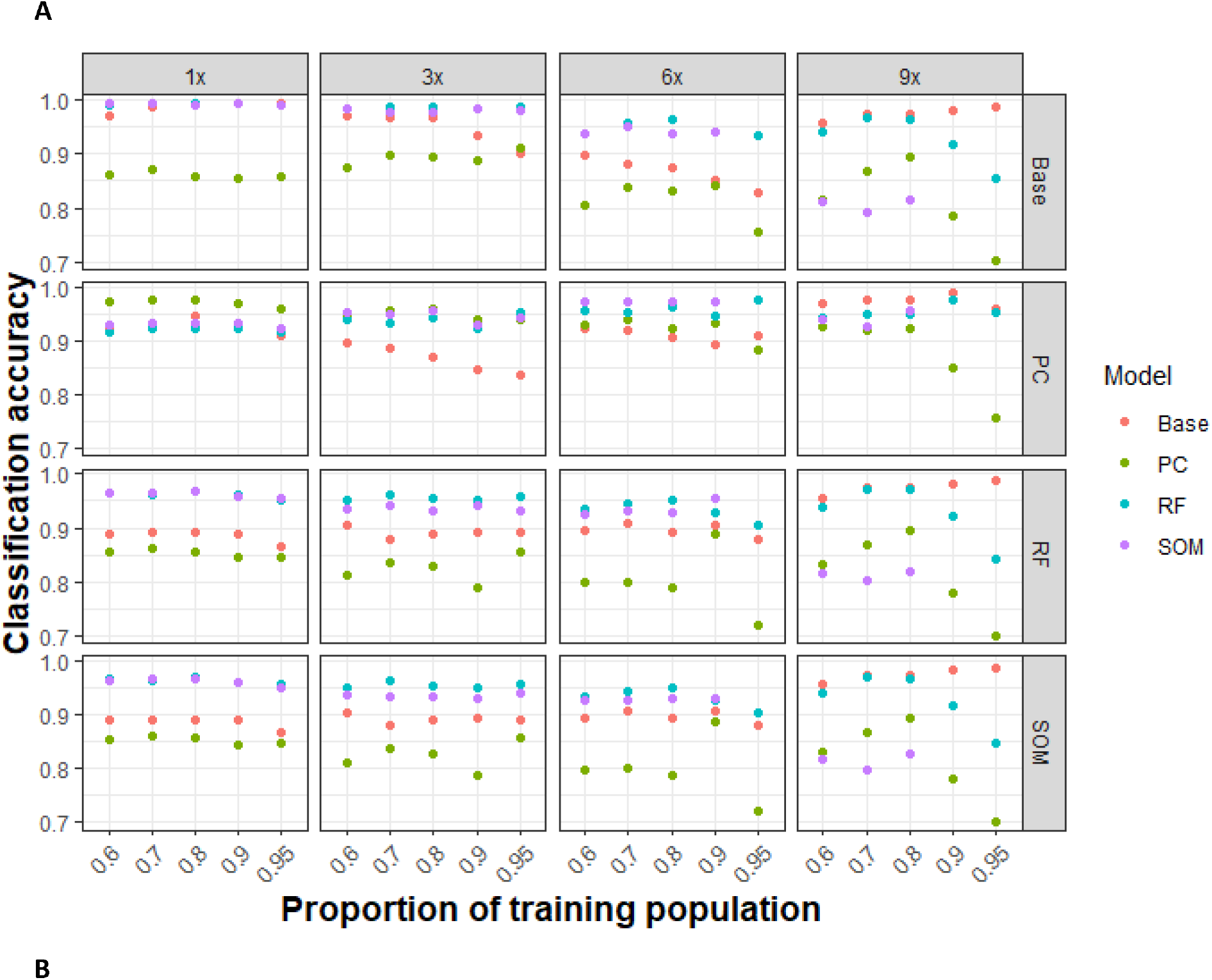
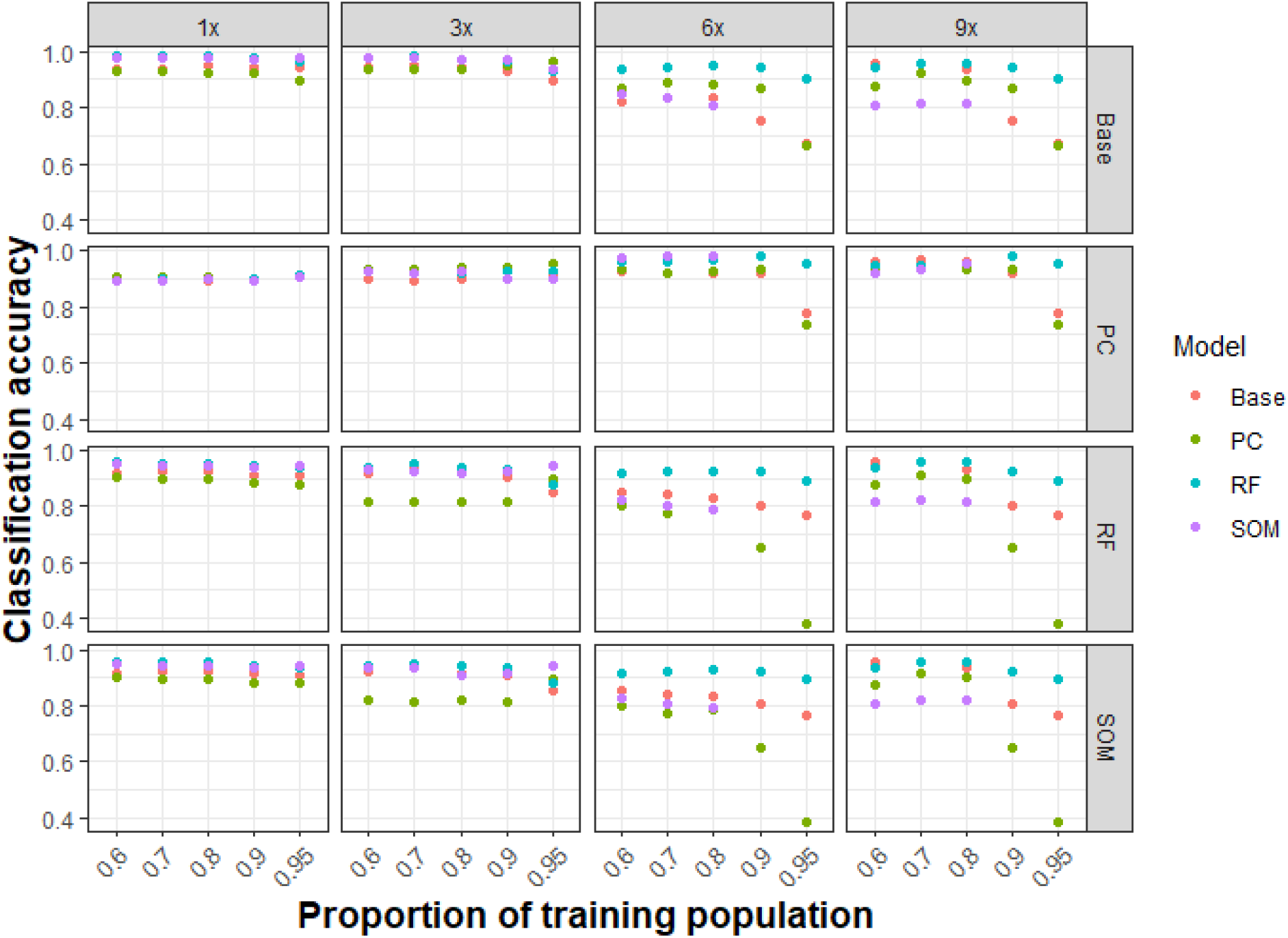
Average classification accuracies from studies using A) all the features (variables) B) most important features which are sodium, calcium, potassium, carbon, phosphorous, and sand. The clusters made from each model (Base, PC – multinomial based on principal components, RF – Random Forest, and SOM – Self Organizing Maps) are represented in horizontal panels while the unit area for observations is represented in the vertical panels. The smallest unit area is a segment in a block (∼7 × 2 m) which is expressed as ‘x’. The other three areas are represented as the multiples of this segment. The predictability of all the models which are represented in four different colors is checked against the clusters produced by a specific model. The predictability was checked using varying training population size. Overall, RF was found to the most robust, versatile, and best performing model. RF and SOM performed similarly except when i) the area was increased a.k.a the number of observations was reduced and ii) clusters were based on PC.

In general, predictability can be maintained with fewer number of features, especially when those are the most influencing features in clustering. Size of training population influences the predictability when the number of observations in testing population is reduced to five or less, i.e., when the training population proportion is 90% or above in 6x and 9x unit areas. In general, 70% - 80% of the population can be used for training irrespective of the total number of observations for better accuracy in prediction.

## 4. Discussion

Knowing the characteristics of the field is a critical element for success in precision agriculture for which soil features play a major role. Collecting fine-resolution soil features helps in understanding not only the dissimilarities between distant fields but also within a field. This kind of information is even more important for minor crops and marginal farmers. Based on the features, the management practices such as dose and frequency of fertilizers can be customized for the field. In our study on representative TEs of safflower, we found that the different locations can be distinguished based on their unique soil characteristics. Moreover, we found dissimilarities in soil characteristics within a location.

We recommend to collect soil before the cropping season from sample sites in the TEs. An area of 170 m^2^ is the largest unit area we found to be useful in collecting soil samples from our exercise. A comprehensive study using edaphic as well as topographic features from blocks in multiple sites within a location will provide a better insight on the optimal area for sample collection. The statistical models such as the RF and SOM from our study can be used to identify the major distinguishable features within and between the locations. Collection of soil features from more sites within the TE can be limited to these major distinguishable features. These features can also be used as the base for customizing the management practices in the cropping season. We recommend RF model for predicting similar sites for a cluster. Similar management practices can be applied to a cluster when finalized.

The RF and SOM models considers the underlying relationship among the features while training the model which eventually helps in identifying the clusters with low wss. Although PCA uses the underlying relationship, the variability of data is limited to the first two PCs which results in practically less useful cluster formation.

Representation of variation in the test data influences the prediction accuracy. Therefore, the SOM models fails when the proportion of training population is 90% or higher in 6x and 9x datasets. Transferring the codebook information and finding meaningful distance between objects is challenging when the number of observations decrease in the test population. In this context, the higher prediction accuracy of RF model is significant even in small datasets. For very small datasets, a training population proportion varying from 70 – 80% is optimal for all the models for proper distribution of the variation and relationship among the soil features. A marginal decrease in the prediction accuracy from training population proportion of 0.6 compared to 0.7 and 0.8, especially in 6x and 9x unit areas could be because of the under representation of the relationship among features. The concept of design of experiments with optimal area and number of observations should be used in edaphic surveying for smart agriculture to represent the variation.

Out of the thirteen features we evaluated, calcium, sand, soil organic carbon, phosphorous, potassium, and sodium are found to be most influencing in distinguishing the representative TE into unique clusters. These features represent the uniqueness of the clusters as the soil samples for analysis are collected before the crop season. Based on this information, we expect to modify the standard management practice for safflower cultivation suitable for specific clusters.

We found that calcium is the most influencing soil feature in distinguishing the clusters. In India, nearly one-third of the soil is calcareous (Bhattacharyya et al., 1994). Maharashtra, Andhra Pradesh, Gujrat, Rajasthan, and Tamil Nadu are the states which confine calcareous soil due to the presence of calcium carbonate (CaCO_3)_ during the paedogenic process. In our study, high calcium content is present in Pune and Nagpur with average values of 696mg/100g and 496mg/100g respectively. Whereas Haveri (145mg/100g) and Delhi (89.5mg/100g) have moderate calcium content. Calcium is a macronutrient, which plays a dual role in plants: a structural component in the cell wall as well as a secondary messenger (Thor, 2019). It is involved in the stabilization of the cell wall structures, regulates the ion transport and selectivity, as well as enzyme activities of the cell wall in safflower (Hossein Alavi et al., 2010). Calcium deficiency causes stunted growth, chlorosis and decreases immunity in the plant whereas abundant calcium interferes with the uptake of the micronutrient, which create a deficiency of another nutrients (Pandey, 2018).

Sandy soil has good drainage property. In addition to having high calcium content, Pune soil has relatively high sand content also. Sandy soil is reported to have high topsoil (5000mg/100g) and subsoil (2000mg/100g) organic carbon content (Armolaitis et al., 2013; Jonczak, 2014). In safflower, productivity is increased through the loosening of the soil and the addition of soil organic matter (Quiroga, 2001). Organic carbon present in the soil can be used as a proxy for organic matter content (Peverill et al., 1999) of it is carbon. Therefore, an increase in soil organic carbon can be interpreted as increase in the fertility of the soil. On an average, Pune soil has the highest organic carbon (672mg/100g) followed by that in Haveri (584mg/100g), Nagpur (544mg/100g), and Delhi (246mg/100g). However, these values are lower compared to the expected value in agricultural field (3800mg/100g) (Yost & Hartemink, 2019) and thus we recommend application of manure before every crop season in the TE.

Although the importance of potassium, a major nutrient, in safflower physiology and morphology is still incipient, it is assumed that potassium will positively involve in growth and development (Silva et al., 2021). From our study, Nagpur soil is found to have relatively high average potassium (53mg/100g) content while other clusters have less than 20mg/100g of this nutrient. A high amount of potassium in the Nagpur soil can be due to chloritized vermiculites and mica in the soil that may play an important role in the dynamics of soil potassium in calcareous soil (Kaswala, R.R, 1976). Potassium and phosphorus are widely used nutrients in fertilisers. According to Safflower package of practices (<u>Package of practices | ICAR-Indian Institute Of Oilseeds Research</u> (icar-iior.org.in)), safflower required 168mg/100g of phosphorus and 56mg/100g of phosphorus in Karnataka region, whereas 90mg/100g of phosphorus is required in Maharashtra soil. However, in our study, we found that phosphorus and potassium are in less amount in the soil and thus, required fertiliser treatment throughout growing season.

Phosphorus, another major nutrient, stimulates seed yield in safflower (Golzarfar et al., 2011). It is also critical for cellular processes (Hasanuzzaman et al., 2018) cell division, carbohydrate metabolism, and root development (Razaq et al., 2017). We found that Nagpur and Haveri soils have relatively high phosphorus content with average values 2.7 and 2.4 mg/100g respectively while other clusters have approximately 1.2 mg/100g of the nutrient. Various factors including pH, clay, and exchangeable Ca in the soil influences the availability of phosphorus. As exchangeable Ca and pH increases, the available phosphorus for the plants decreases due to its solubilization (More et al., 1979). Thus, highly calcareous soil of Pune, with the basic pH of ∼7.5 is found to have low available phosphorus whereas the soil in Nagpur, again with high Ca content but with acidic pH of 6.5 is found to have high amount of the nutrient.

Sodium is a “functional nutrient” as plants can survive without it (Subbarao, G. V., et al., 2003). In safflower, sodium stimulates plant height and dry matter absorption (Bains & Fireman, 1964). It is assumed that sodium could be used as a substitution for phosphorus in fertilizers, however, in safflower sodium and phosphorus play their independent functions (Amorim Silva, 1972). Abundance of sodium tends to increase the salinity of the soil and thus causes salt stress in the plant. In our study, we found the sodium levels are beyond the critical value (13mg/100g) (https://ucanr.edu/) for Nagpur (71mg/100g) and in the cluster formed by blocks A and B in Delhi (26mg/100g). Soil salinity depends on the parent rock, soil type, climate, pH, and irrigation routine (Bauder & Brock, 2001). High salinity in Nagpur can be attributed to smectite-dominant shrink-swell soils (Thakare et al., 2013) whereas salinity in the Delhi cluster formed by blocks A and B can be attributed to high pH and irrigation routine. Also, that particular cluster in Delhi is nearby the housing locality, which may also contribute to the high salinity in the soil. Being adapted in semi-arid regions, safflower can be expected to grow in high saline conditions provided other nutrients are adequately available.

The blocks in Delhi are kept the furthest apart within an area of ∼40,000 m^2^ with an arrangement that a pair of blocks are kept at a distance of 160 m apart and this pair is separated from the other pair by a distance of 350 m. The pairs, although from the same location, showed distinct characteristics to be accommodated into separate clusters. Delhi cluster with blocks A and B has higher calcium, sand, and phosphorus as compared to cluster with blocks C and D with higher organic carbon. This variation even within a field thrusts the importance of fine resolution soil surveying to facilitate smart farming.

Although fine-resolution mapping is prevalent globally, it is still in its incipient stage in India (Dash et al., 2022; Searle et al., 2021; Thompson et al., 2020). SoilGrids 2.0 (https://soilgrids.org/) and OpenLandMap (https://openlandmap.org/) are the two DSM which provide information at the fine-resolution of 250 m^2^ in India (Hengl, Mendes de Jesus, et al., 2017). However, poor sampling density made these maps less accurate (Dharumarajan et al., 2022) and thus, plot (0–1 km^2^) and local (1 km^2^ –10^4^ km^2^) level studies are required. Our study with high sampling density and fine-resolution (18 observations per 340 m^2^) highlights the significance of such studies and can pave the way for precision farming and increased agriculture efficiency even for small farms and marginalised crops.

## 5. Conclusions

On identifying the most distinguishable variables from this fine-resolution study, a site-specific recommendation can be done for optimum use of nutrients. Site-specific recommendations are also a useful and effective tool for government initiatives in the agriculture sector such as soil health cards, and delineating suitable crop-growing areas (Dharumarajan et al., 2020; Santra et al., 2017). In conclusion, we present a fine-resolution, high-density site-specific digital soil map for the safflower growing areas which can be useful in the nutrient recommendation and are unique and significant in terms of its application as well its methodology.

## Abbreviations

ANN: artificial neural network
AS: Ankur seeds
B: Boron
C: Carbon
Ca: Calcium
Cl: Chlorine
Cu: Copper
CV: cross-validation
DL: Deep learning
DSM: Digital soil mapping
Fe: Iron
GWA: gravimetric water content
H: Hydrogen
K: Potassium
Mg: Magnesium
ML: Machine learning
MLR: Multivariate linear regression
Mn: Manganese
Mo: Molybdenum
N: Nitrogen
NARI: Nimbkar Agriculture Research Institute
O: Oxygen
P: Phosphorus
PAM: Partitioning around medoids
PC: principal components
PCA: Principal component analysis
RF: Random forest
S: Sulphur
SOM: Self -organizing maps
TE: Target environment
U-matrix: unified distance matrix
Zn: Zinc

## Acknowledgment

We thank the University of Delhi for providing the analysis facility in the soil analytical lab lead by Dr. Ratul Baishya and conducting the experiment in the research field. We also thank the soil analysis lab at the Indian Agricultural Research Institute for conducting the soil analysis. We thank the managing director and staff of Ankur Seeds Pvt. Ltd. Nagpur, president and staff of Nimbkar Agricultural Research Institute, Pune, and the owner and farmer of the field in Haveri, Karnataka.

## Author contributions

AAE: Conceptualization, data curation, formal analysis, funding acquisition, investigation, methodology, resources, software, project administration, supervision, validation, visualization, Writing – original draft, review, and editing

MS: Formal analysis, investigation, methodology, resources, Writing – original draft and review

SG: Project administration, resources, supervision, writing – review, editing

## Funding

This work is supported by the Ramalingaswami fellowship fund provided to AAE by the Department of Biotechnology, Government of India.

## Data availability

Data and algorithm in R will be available on request.

## Declaration of Competing Interest

The authors declare that they have no known competing financial interests or personal relationships that could have appeared to influence the work reported in this paper.

## Declaration of generative AI and AI assisted technologies in the writing process

The authors declare that they have not used AI and AI assisted technologies in the writing process.

## Notes

### Competing Interest Statement

The authors have declared no competing interest.

