## Supplementary files for "Characterization and classification of fine-resolution soil profile for precision agriculture using random forest and self-organizing map"

**Supplementary tables and figures**

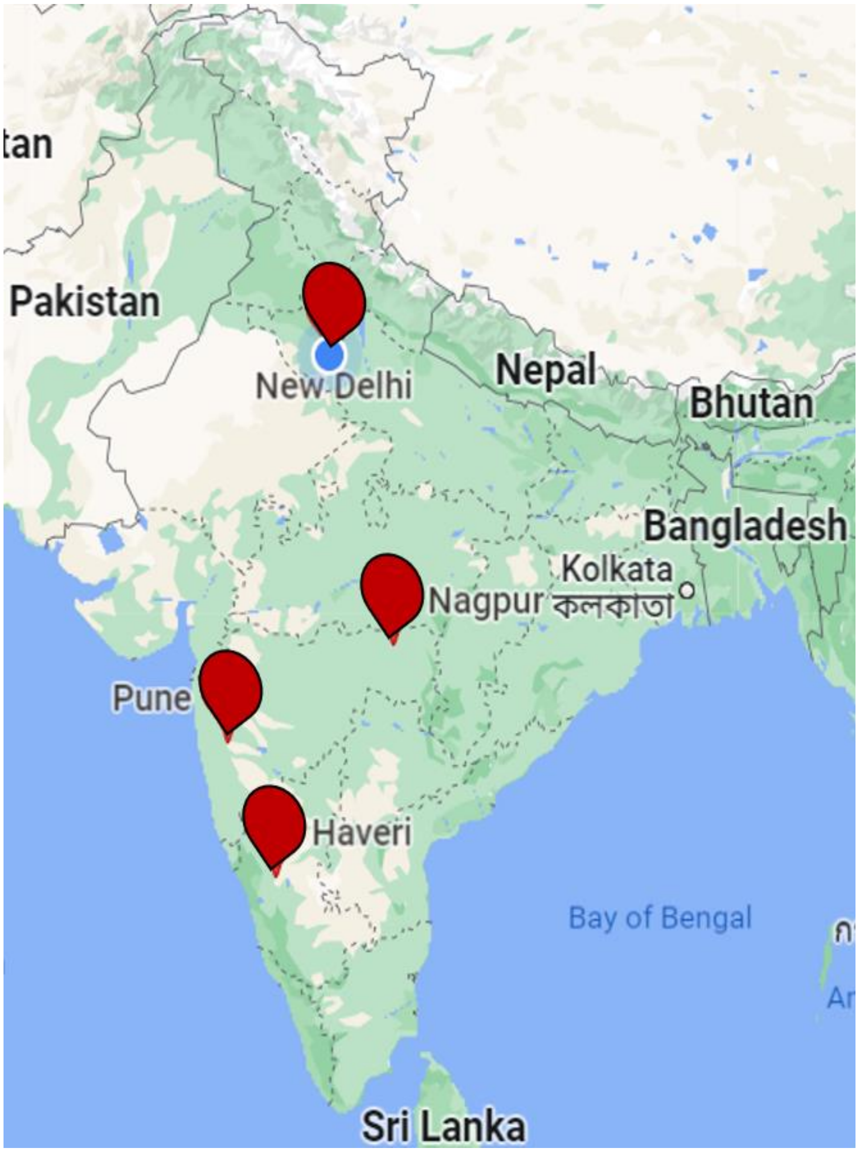


**Fig S1.** Locations representing target environments of safflower cultivation and research field in Delhi. Soil samples were collected from these four locations representing the diversity of soil profile in which safflower grows.

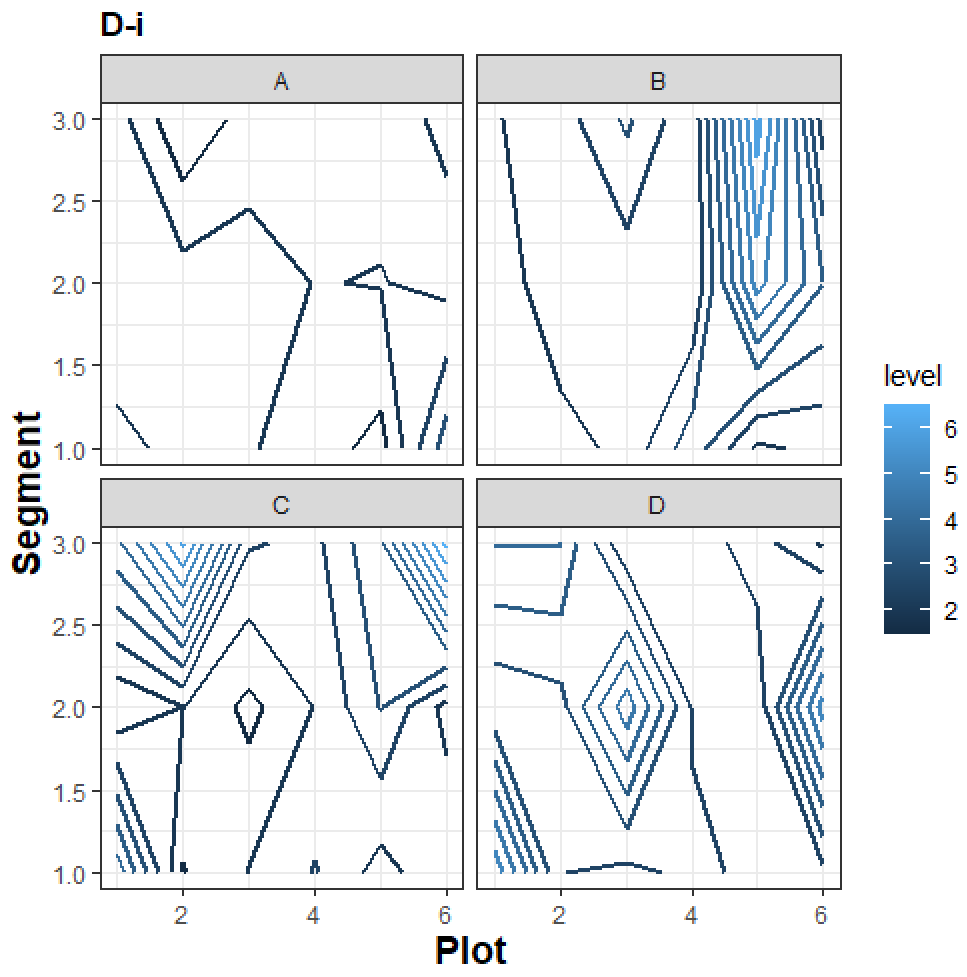
D-i
A
B
C
D
Segment
3.0
2.5
2.0
1.5
1.0
Plot
2
4
6
level
6
5
4
3
2

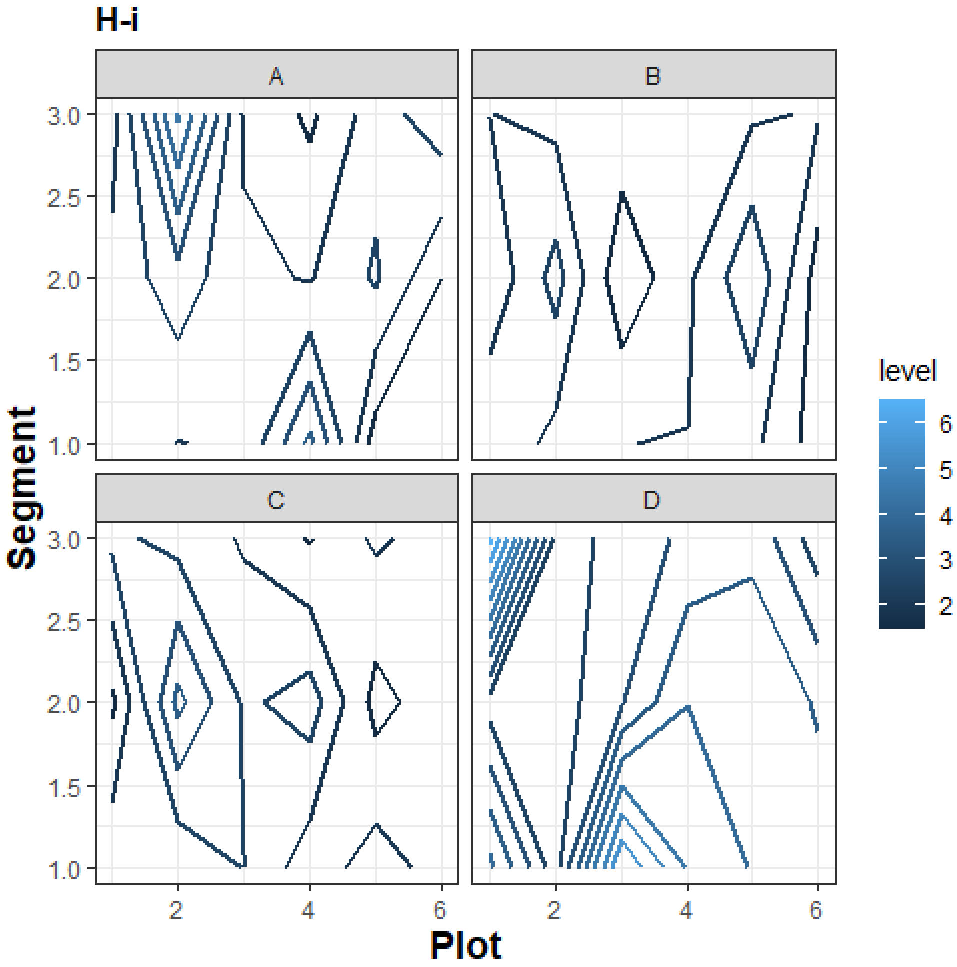
H-i
A
B
C
D
Segment
3.0
2.5
2.0
1.5
1.0
2
4
6
Plot
level
6
5
4
3
2

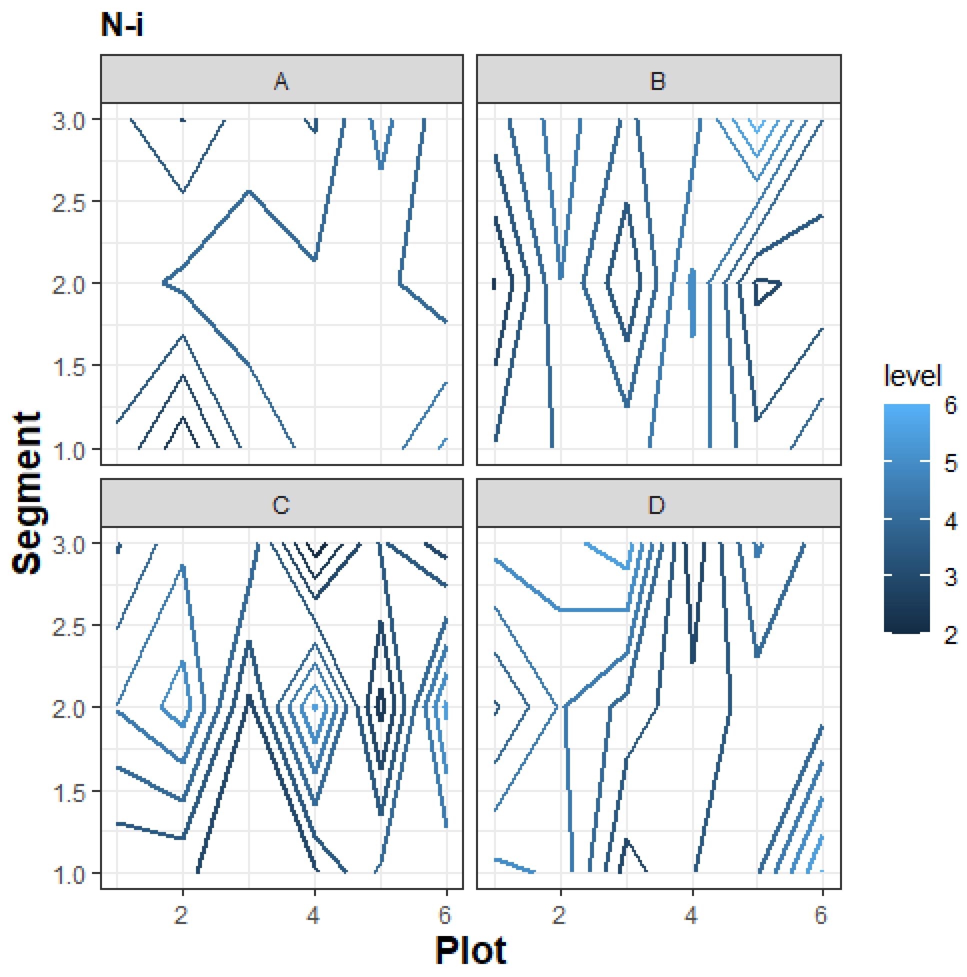
N-i
A
B
C
D
3.0
2.5
2.0
1.5
1.0
Segment
2
4
6
Plot
level
6
5
4
3
2

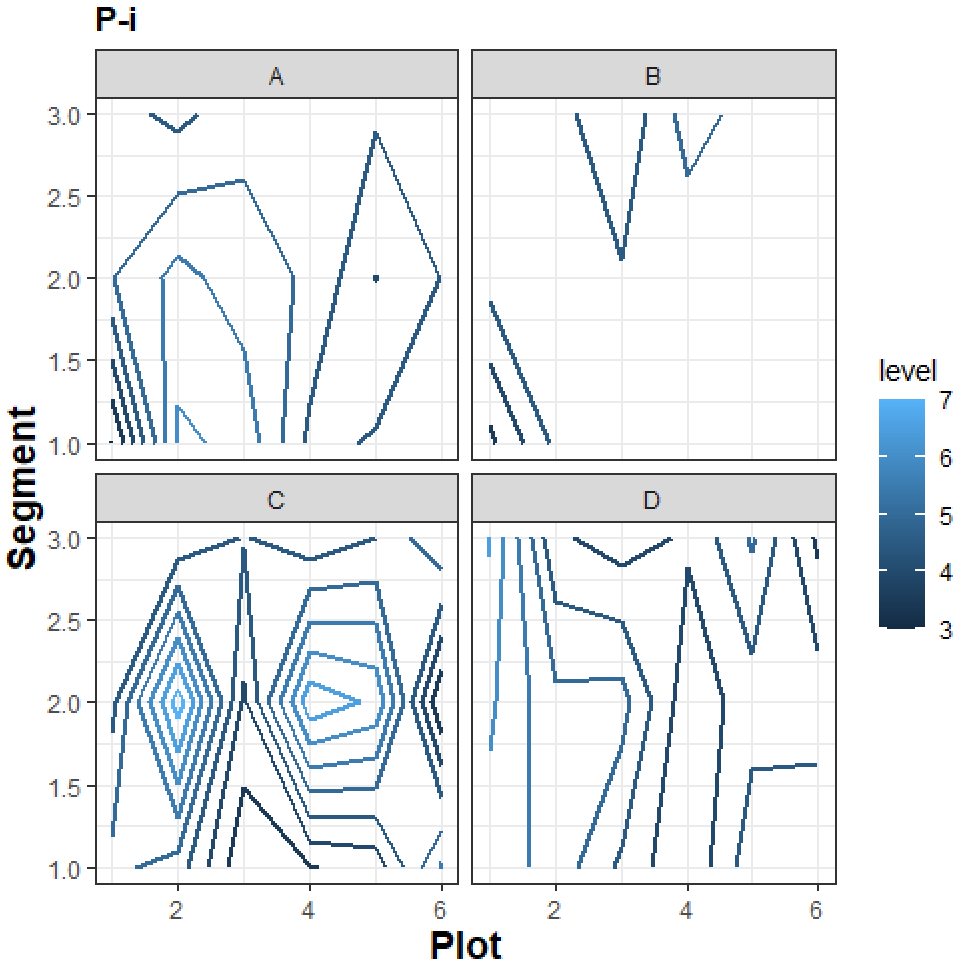

P-i
A
B
C
D
3.0
2.5
2.0
1.5
1.0
Segment
2
4
6
Plot
level
7
6
5
4
3

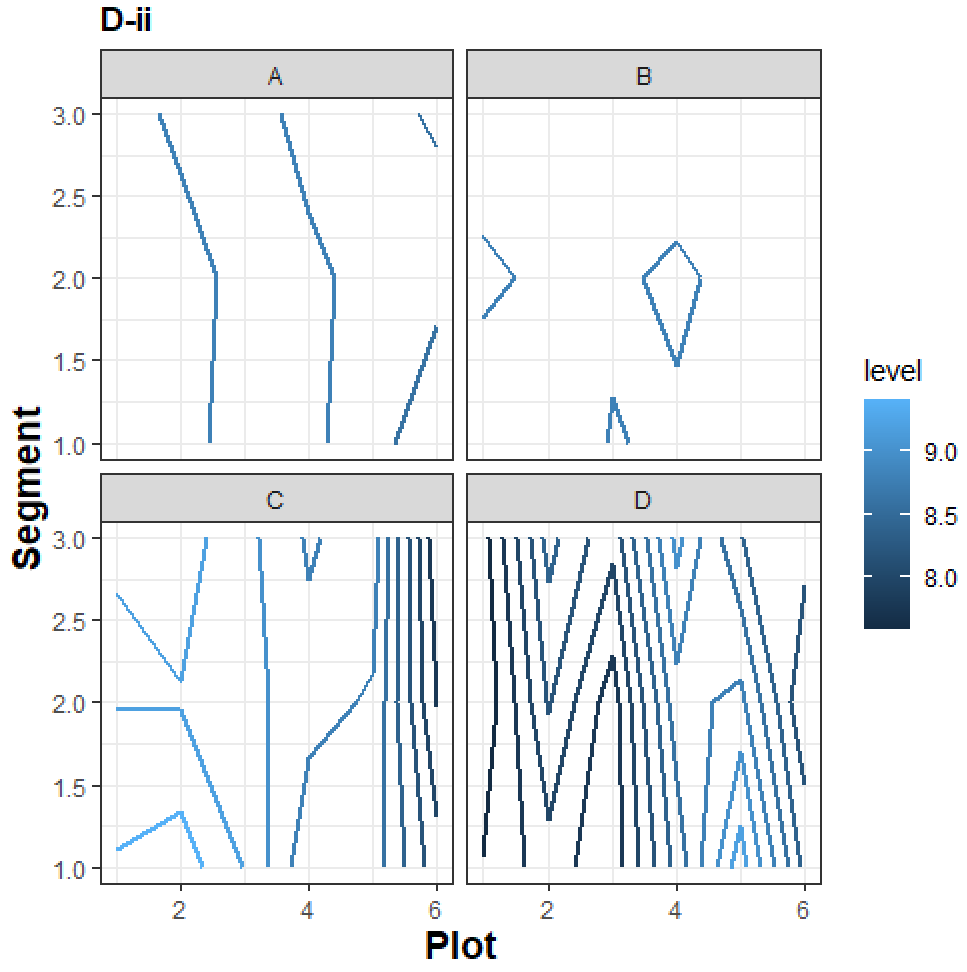
D-ii
A
B
C
D
Segment
3.0
2.5
2.0
1.5
1.0
Plot
2
4
6
level
9.0
8.5
8.0

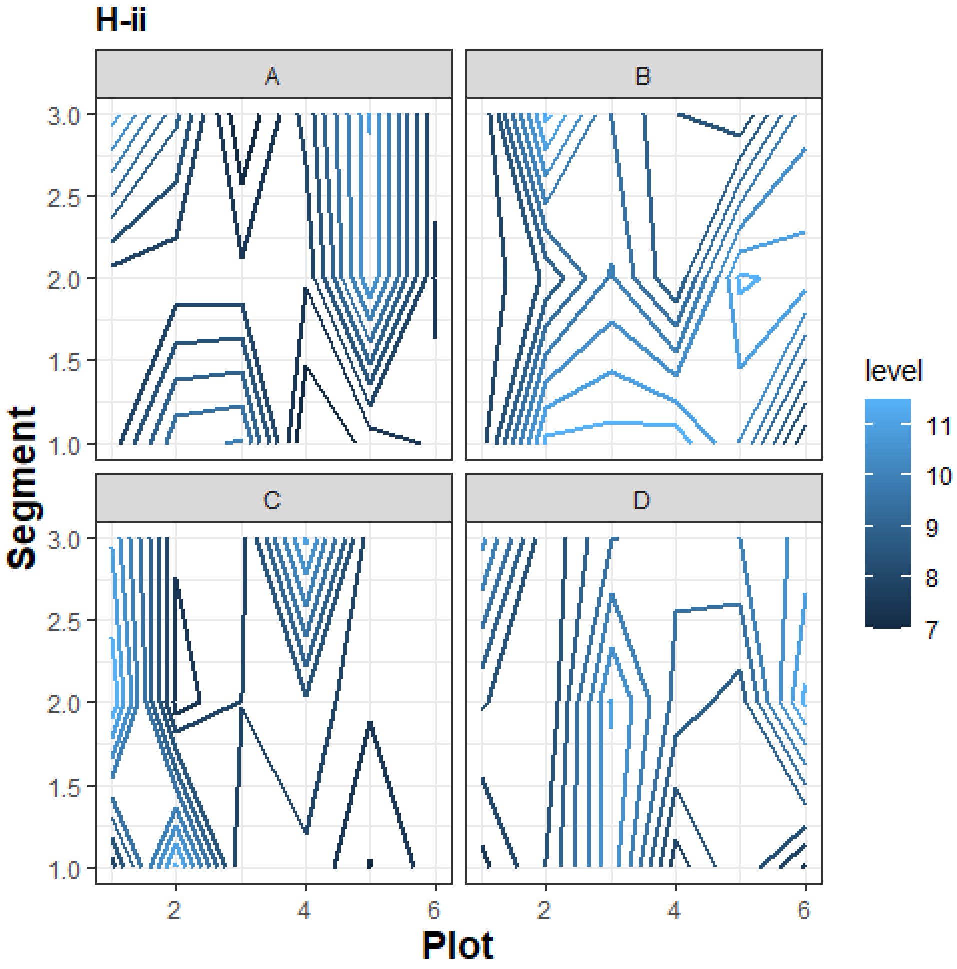
H-ii
A
B
C
D
3.0
2.5
2.0
1.5
1.0
Segment
2
4
6
Plot
level
11
10
9
8
7

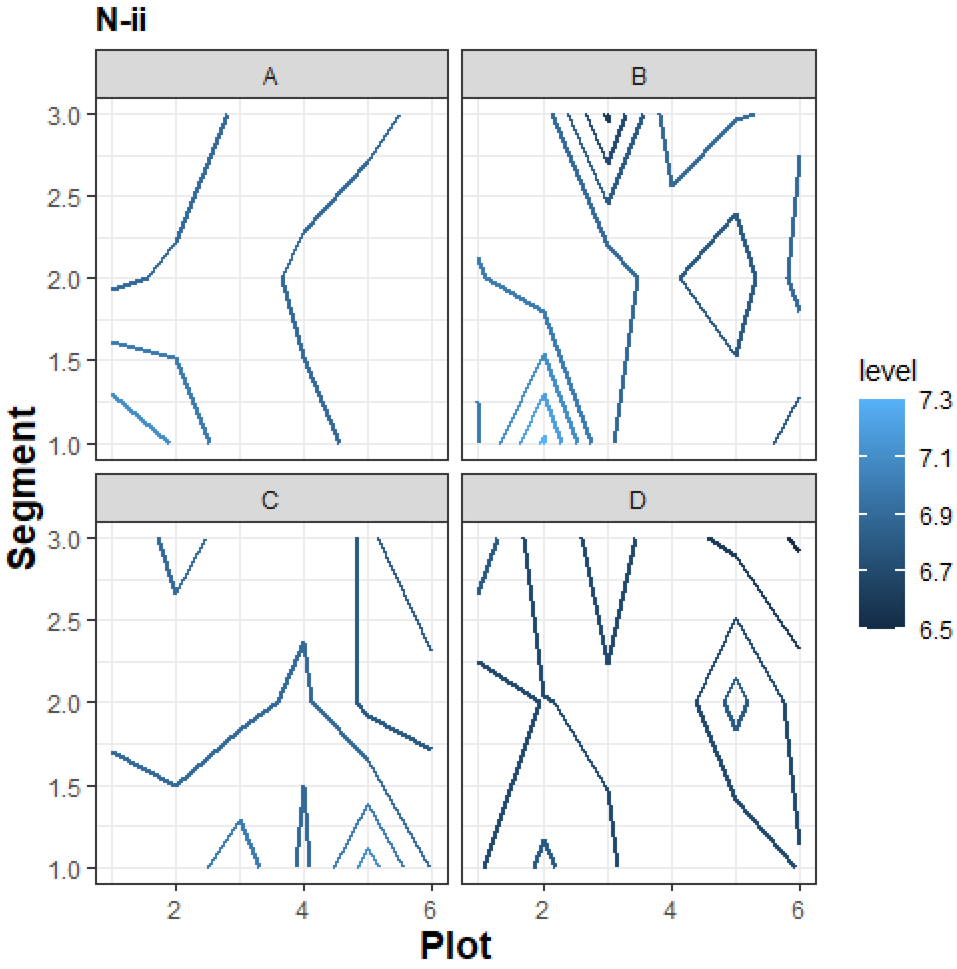
N-ii
A
B
C
D
3.0
2.5
2.0
1.5
1.0
Segment
2
4
6
Plot
level
7.3
7.1
6.9
6.7
6.5

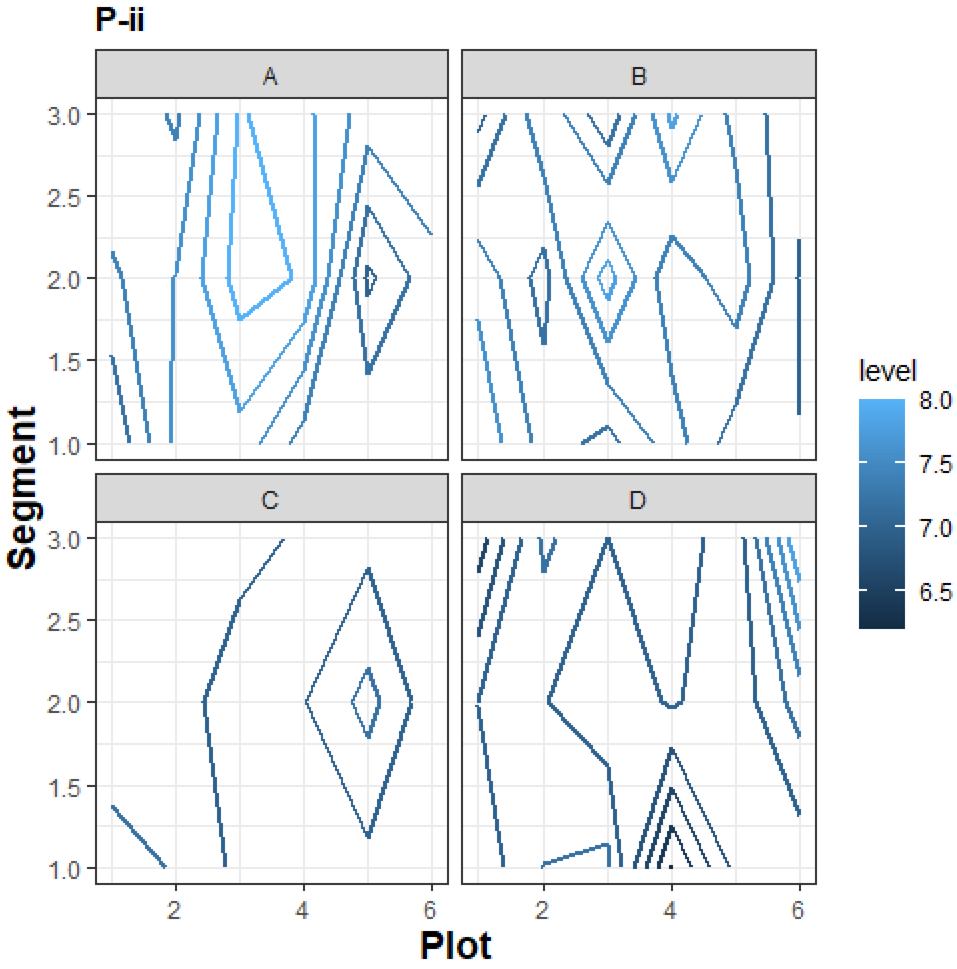

P-ii
A
B
C
D
Segment
Plot
level
8.0
7.5
7.0
6.5
3.0
2.5
2.0
1.5
1.0
2
4
6

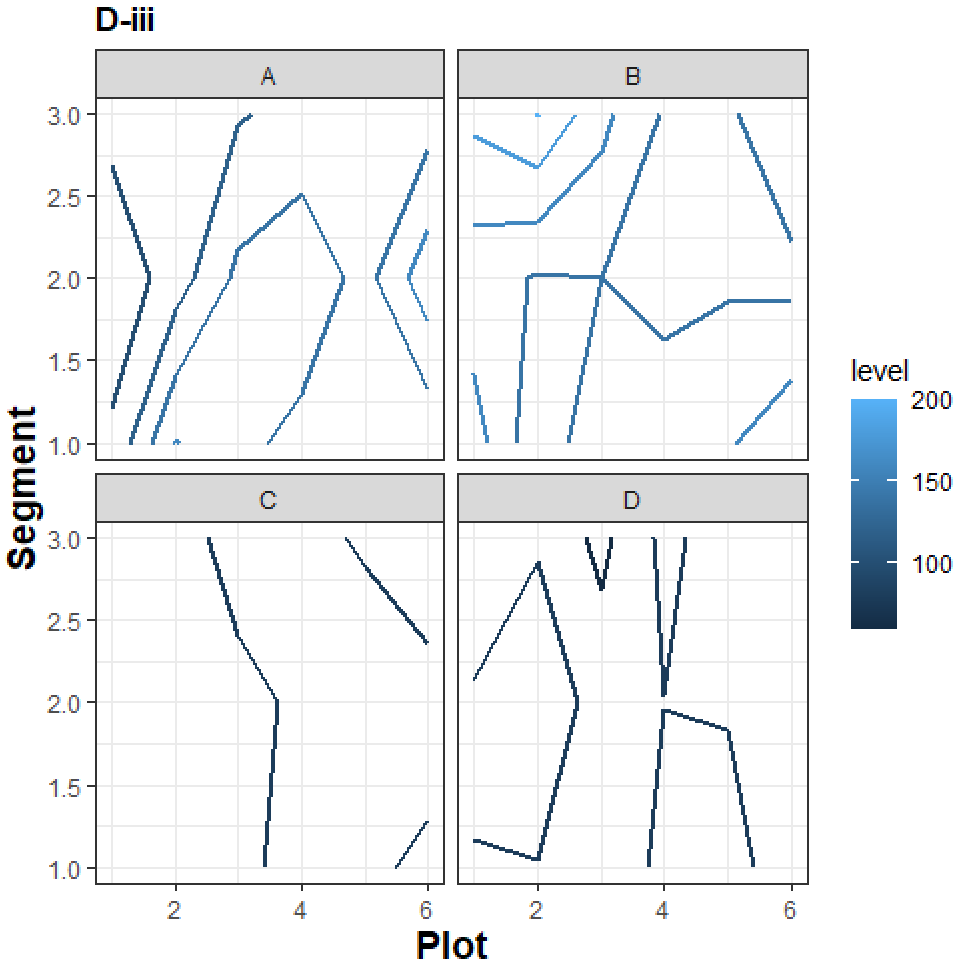

D-iii
A
B
C
D
3.0
2.5
2.0
1.5
1.0
Segment
2
4
6
Plot
level
200
150
100

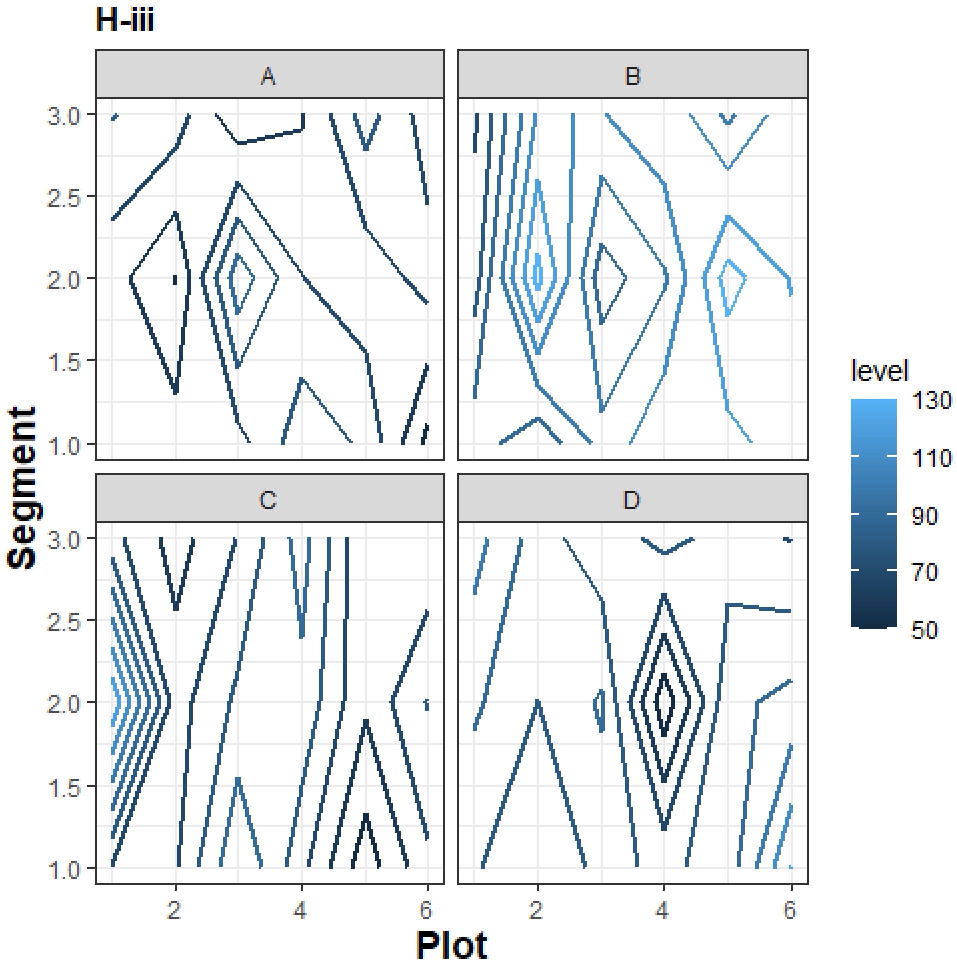
H-iii
A
B
C
D
3.0
2.5
2.0
1.5
1.0
Segment
2
4
6
Plot
level
130
110
90
70
50

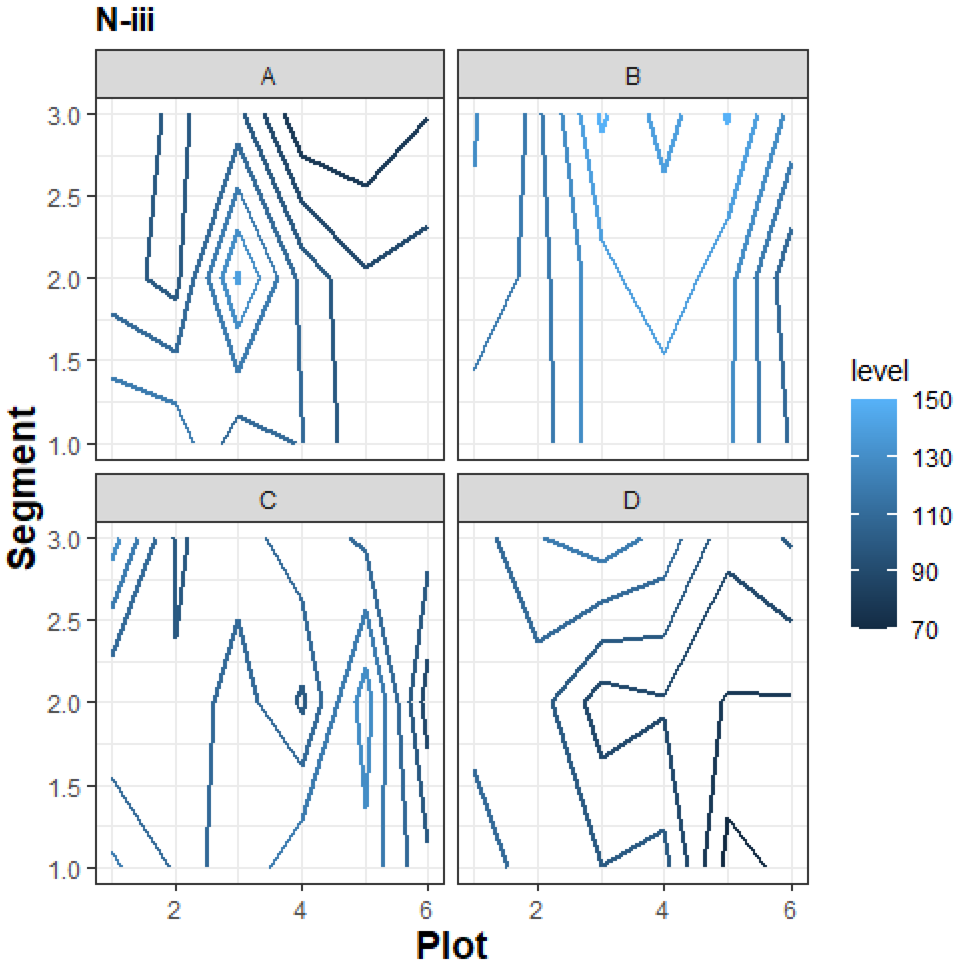

N-iii
A
B
C
D
Segment
3.0
2.5
2.0
1.5
1.0
Plot
2
4
6
level
150
130
110
90
70

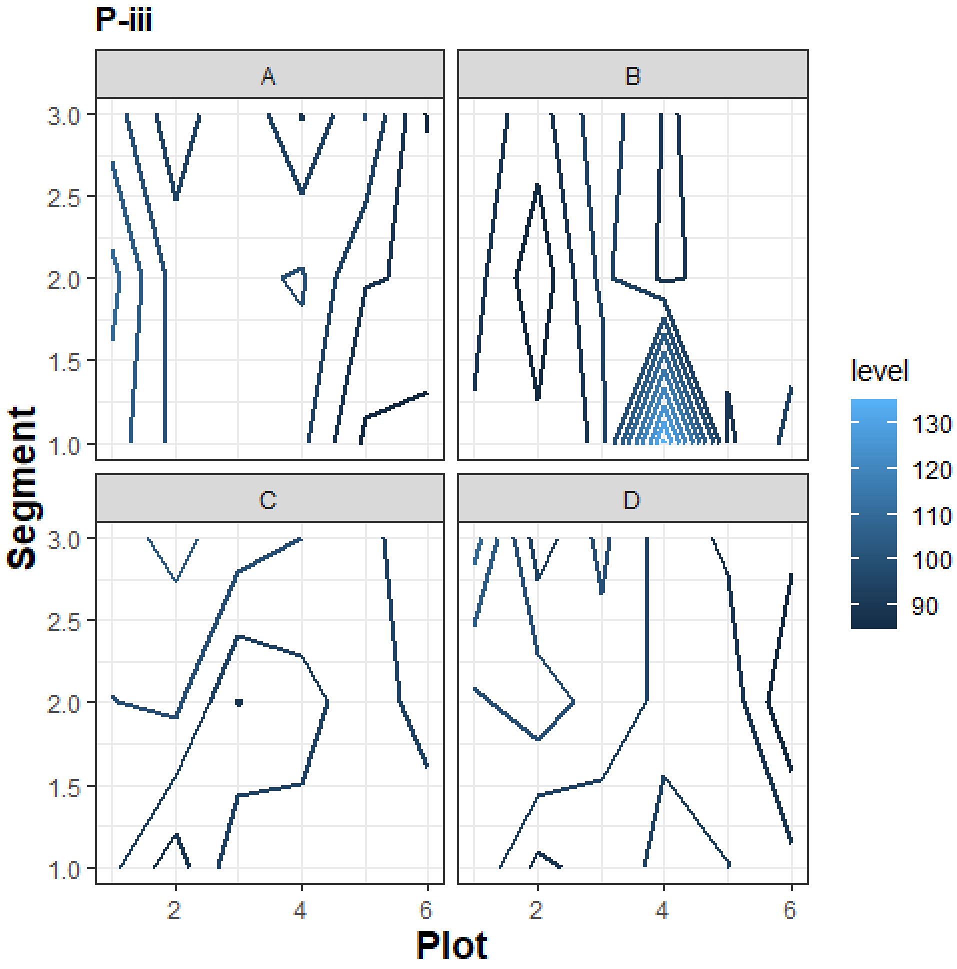
P-iii
A
B
C
D
Segment
3.0
2.5
2.0
1.5
1.0
Plot
2
4
6
level
130
120
110
100
90

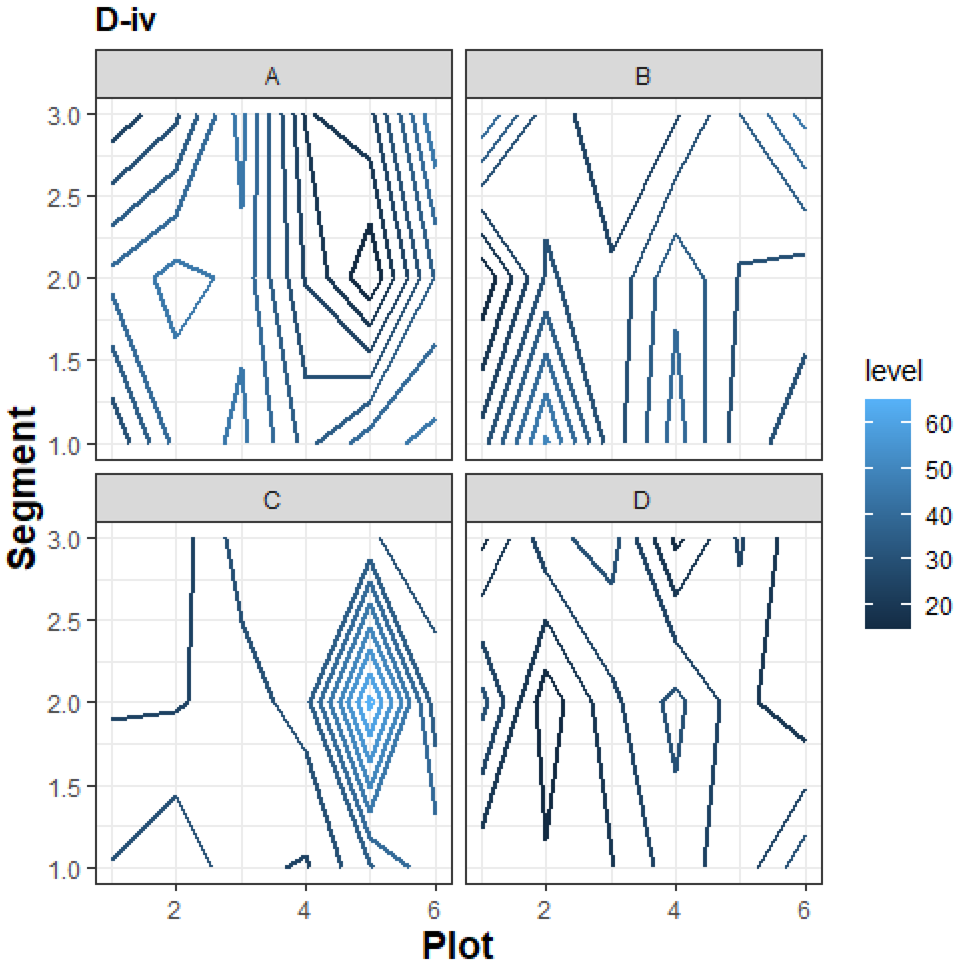

D-iv
A
B
C
D
3.0
2.5
2.0
1.5
1.0
Segment
2
4
6
Plot
level
60
50
40
30
20

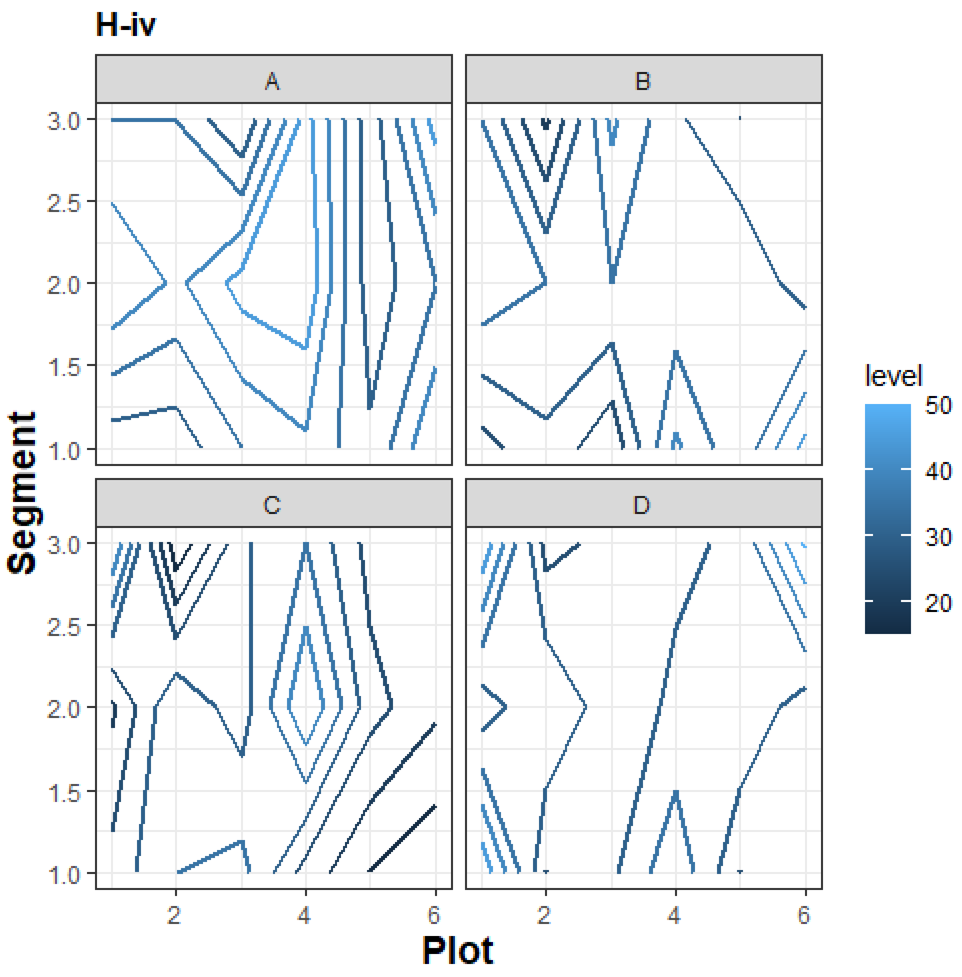

H-iv
A
B
C
D
Segment
Plot
level
50
40
30
20
3.0
2.5
2.0
1.5
1.0
2
4
6

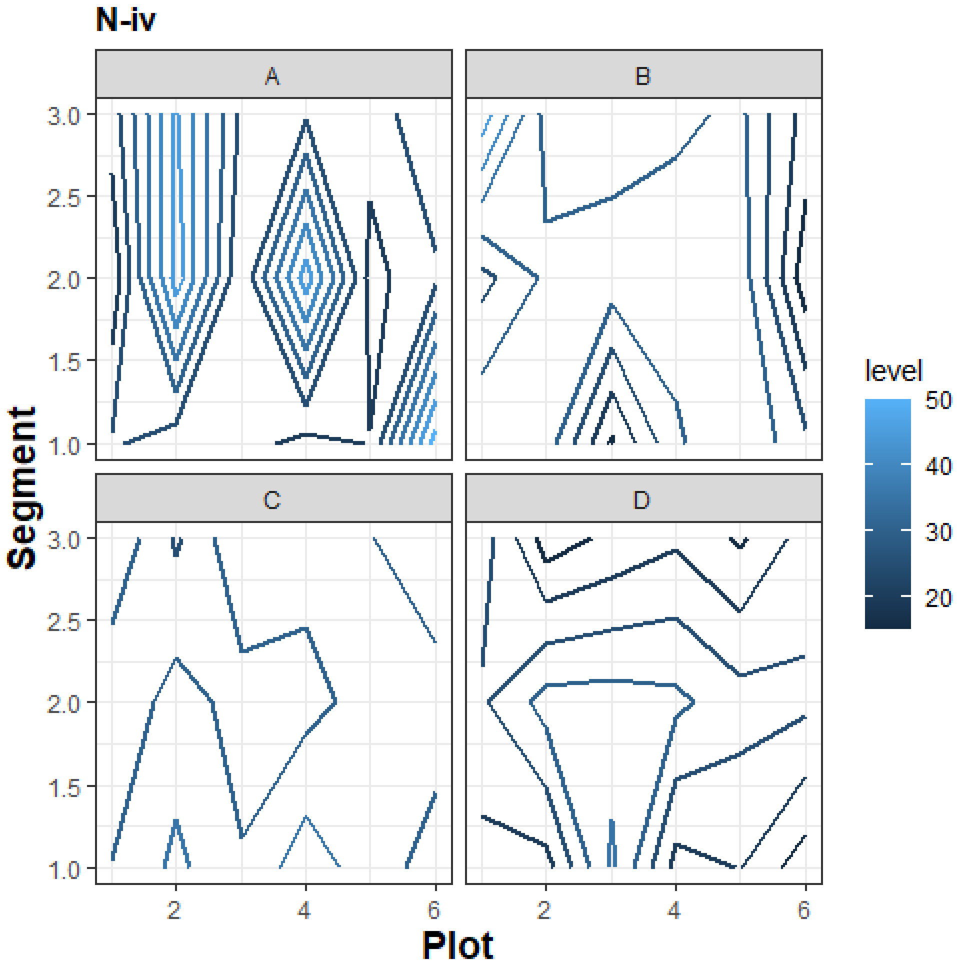
N-iv
A
B
C
D
Segment
Plot
level
50
40
30
20
3.0
2.5
2.0
1.5
1.0
2
4
6

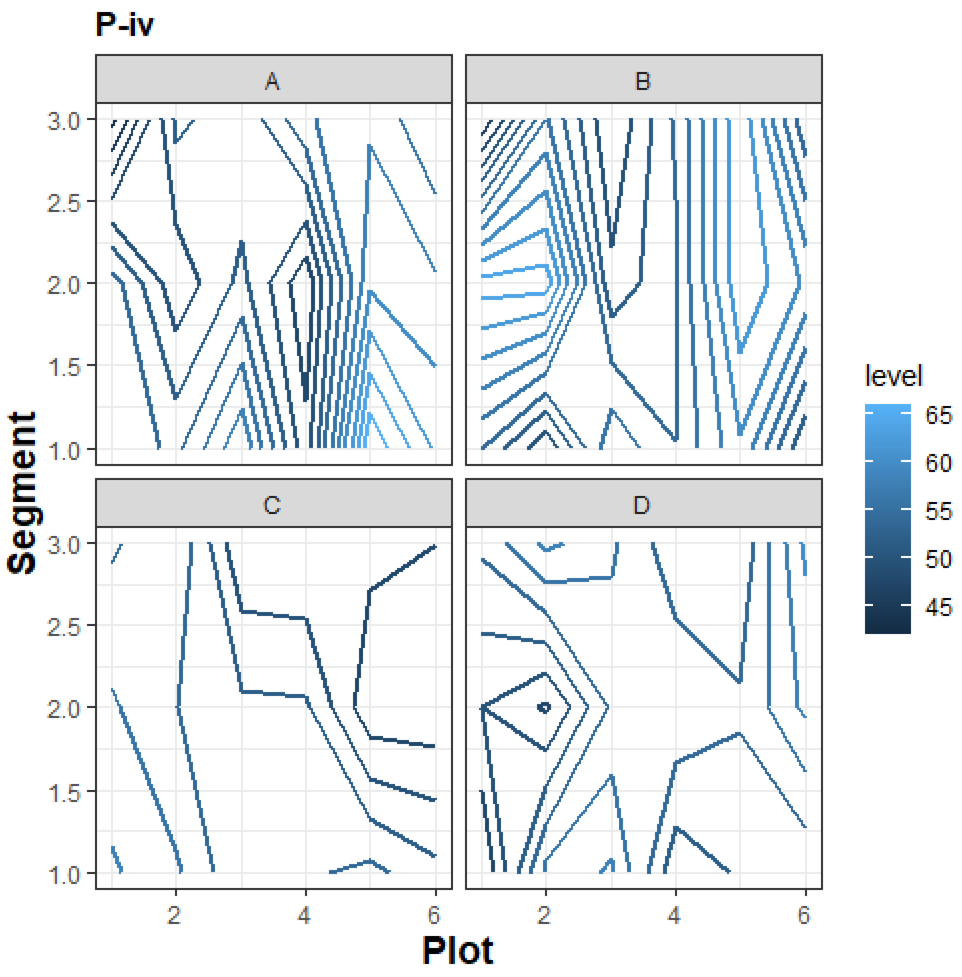
P-iv
A
B
C
D
Segment
Plot
3.0
2.5
2.0
1.5
1.0
2
4
6
level
65
60
55
50
45

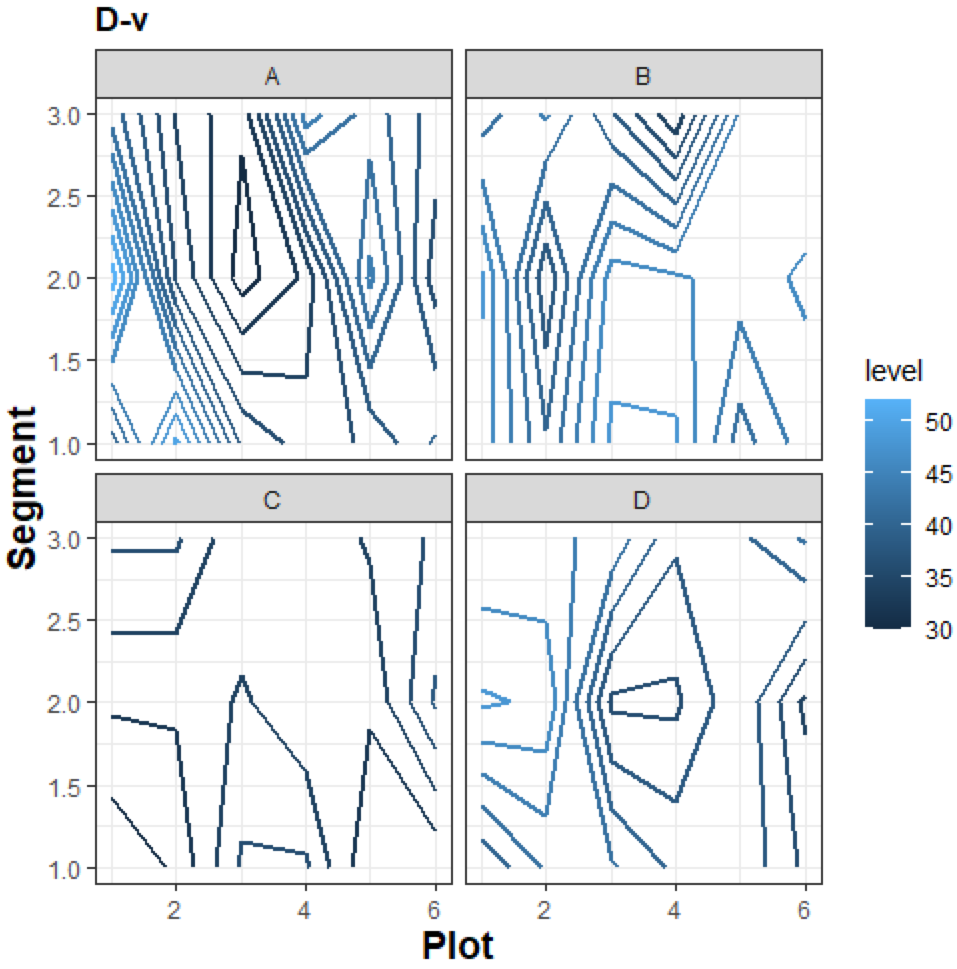
D-v
A
B
C
D
Segment
Plot
level
50
45
40
35
30
3.0
2.5
2.0
1.5
1.0
2
4
6

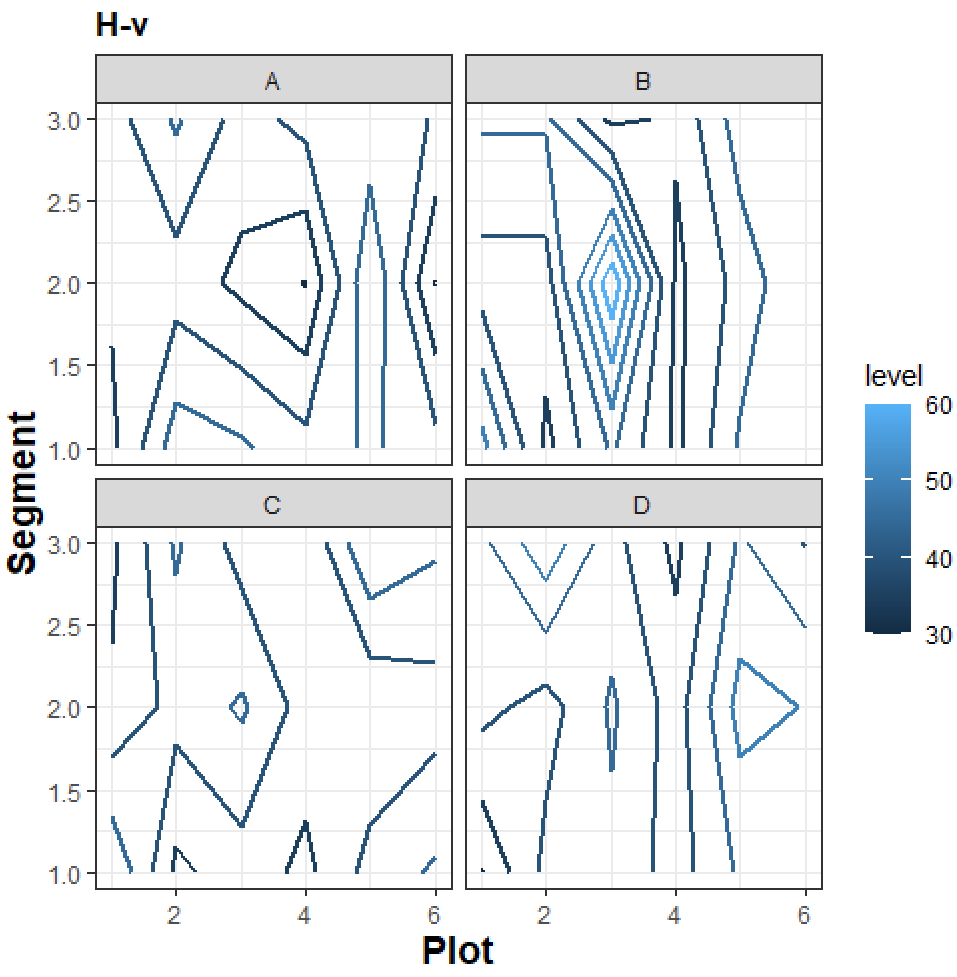
H-v
A
B
C
D
Segment
3.0
2.5
2.0
1.5
1.0
Plot
2
4
6
level
60
50
40
30

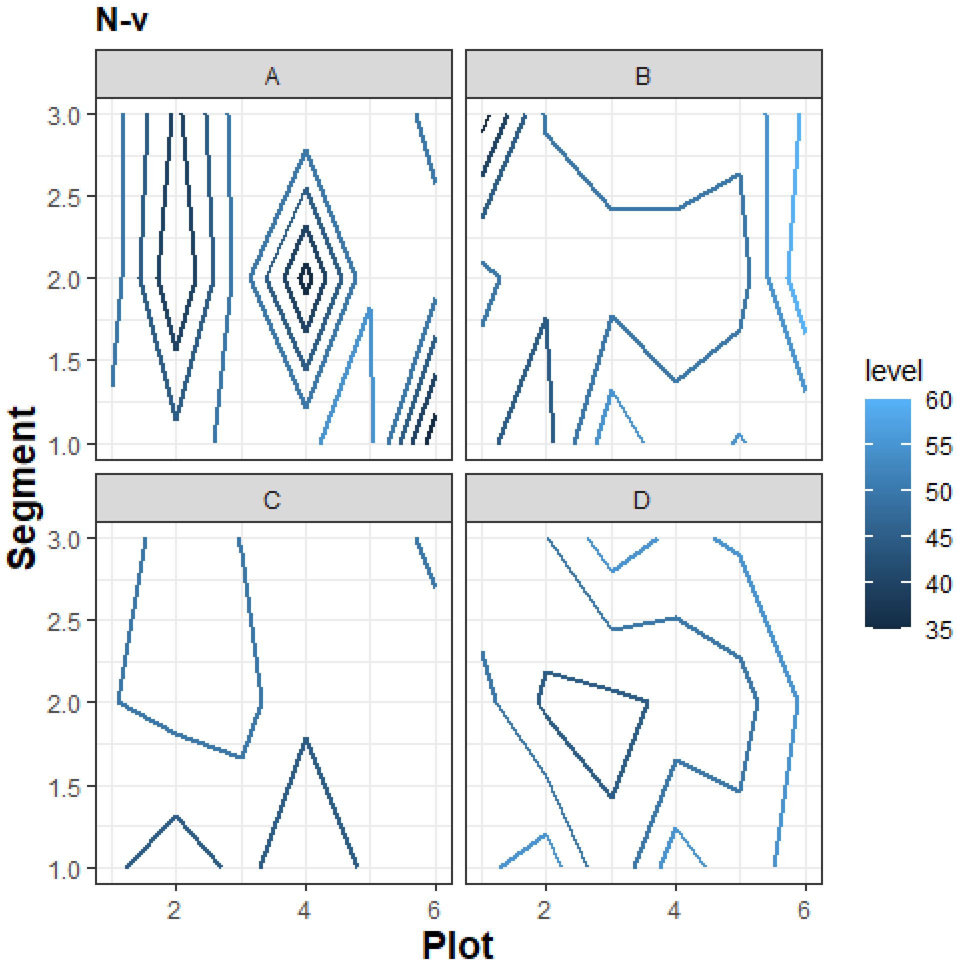
N-v
A
B
C
D
Segment
3.0
2.5
2.0
1.5
1.0
Plot
2
4
6
level
60
55
50
45
40
35

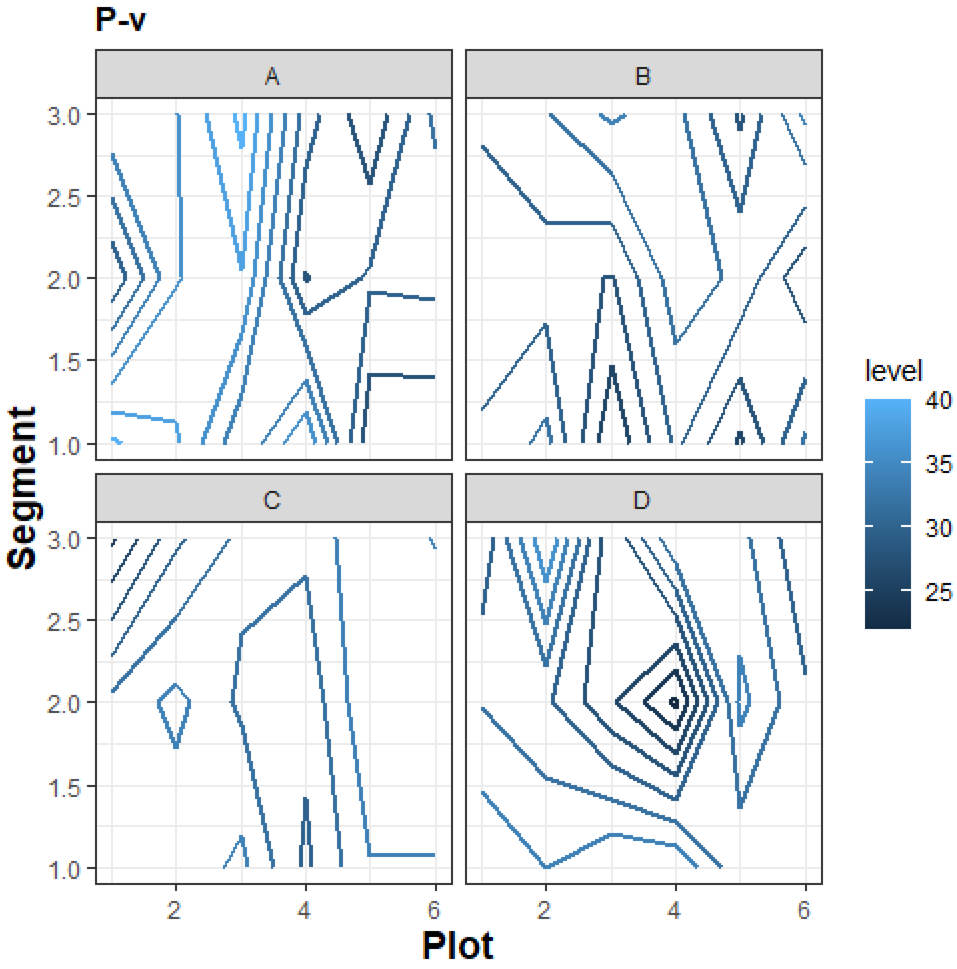
P-v
A
B
C
D
Segment
Plot
level
40
35
30
25
3.0
2.5
2.0
1.5
1.0
2
4
6

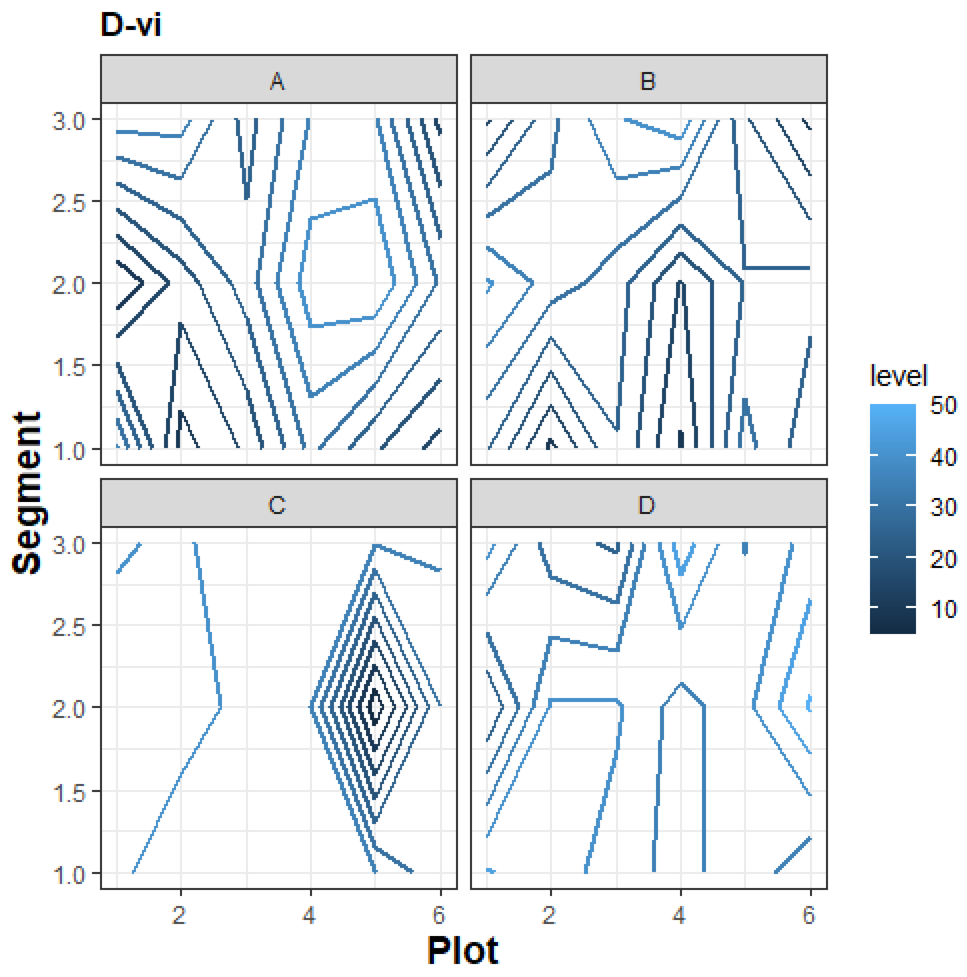
D-vi
A
B
C
D
Segment
Plot
level
50
40
30
20
10
3.0
2.5
2.0
1.5
1.0
2
4
6

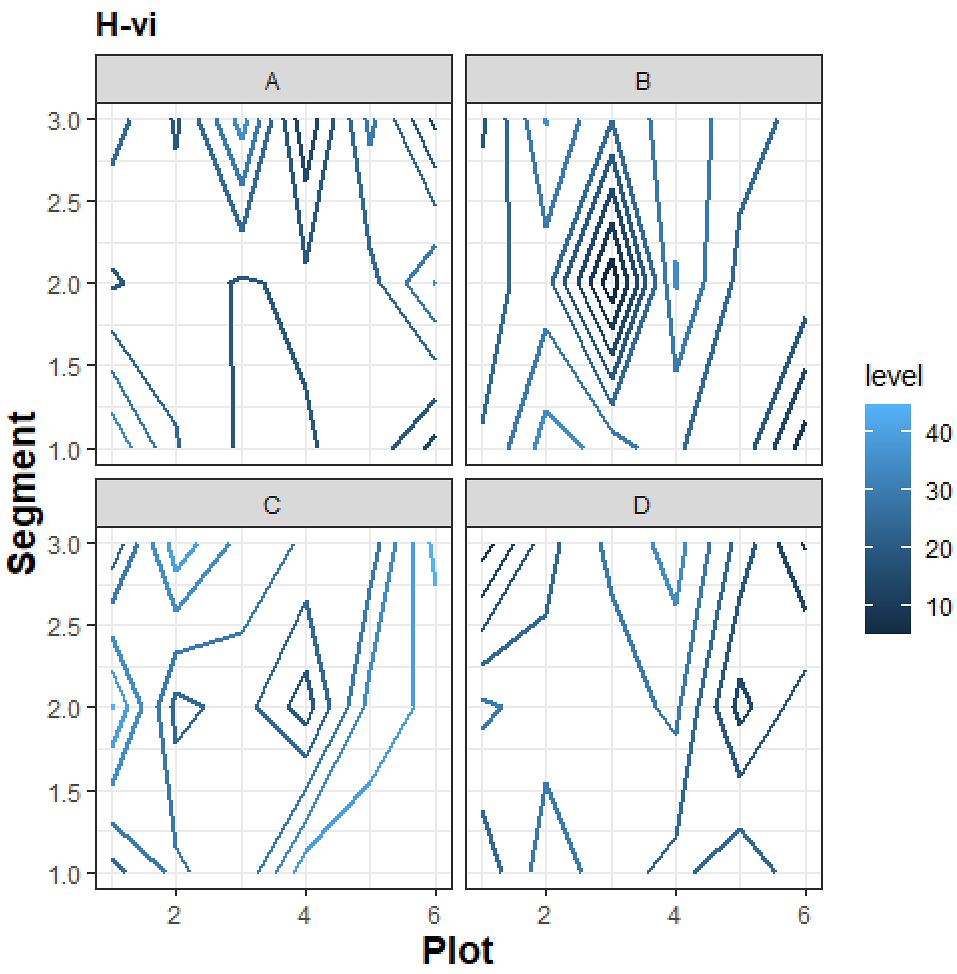

H-vi
A
B
C
D
Segment
Plot
level
40
30
20
10
3.0
2.5
2.0
1.5
1.0
2
4
6

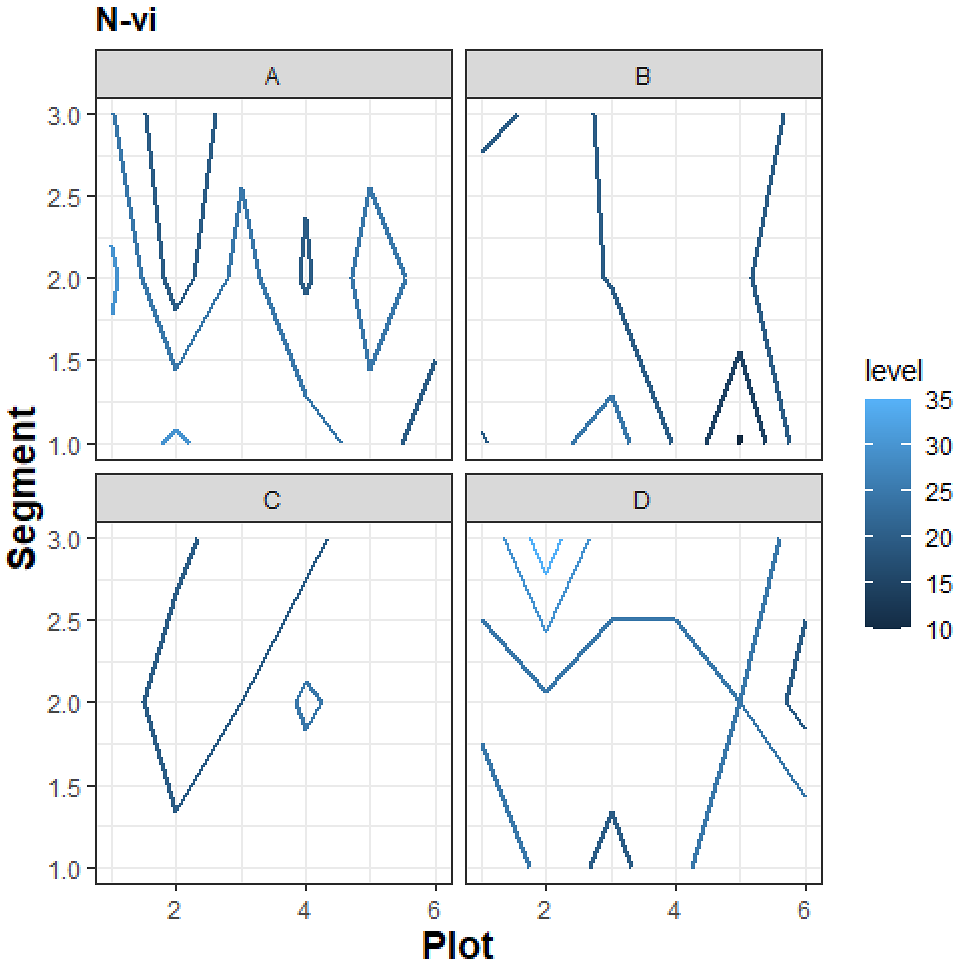

N-vi
A
B
C
D
3.0
2.5
2.0
1.5
1.0
Segment
2
4
6
Plot
level
35
30
25
20
15
10

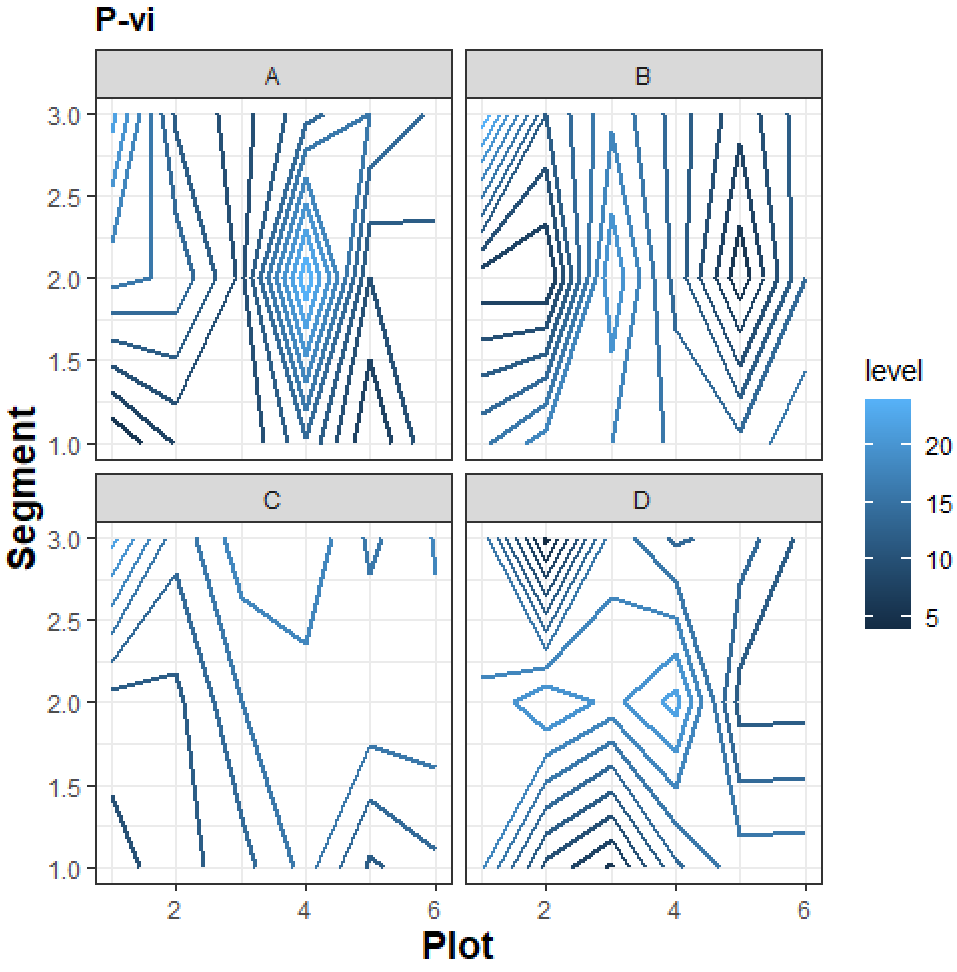

P-vi
A
B
C
D
Segment
Plot
level
20
15
10
5
3.0
2.5
2.0
1.5
1.0
2
4
6

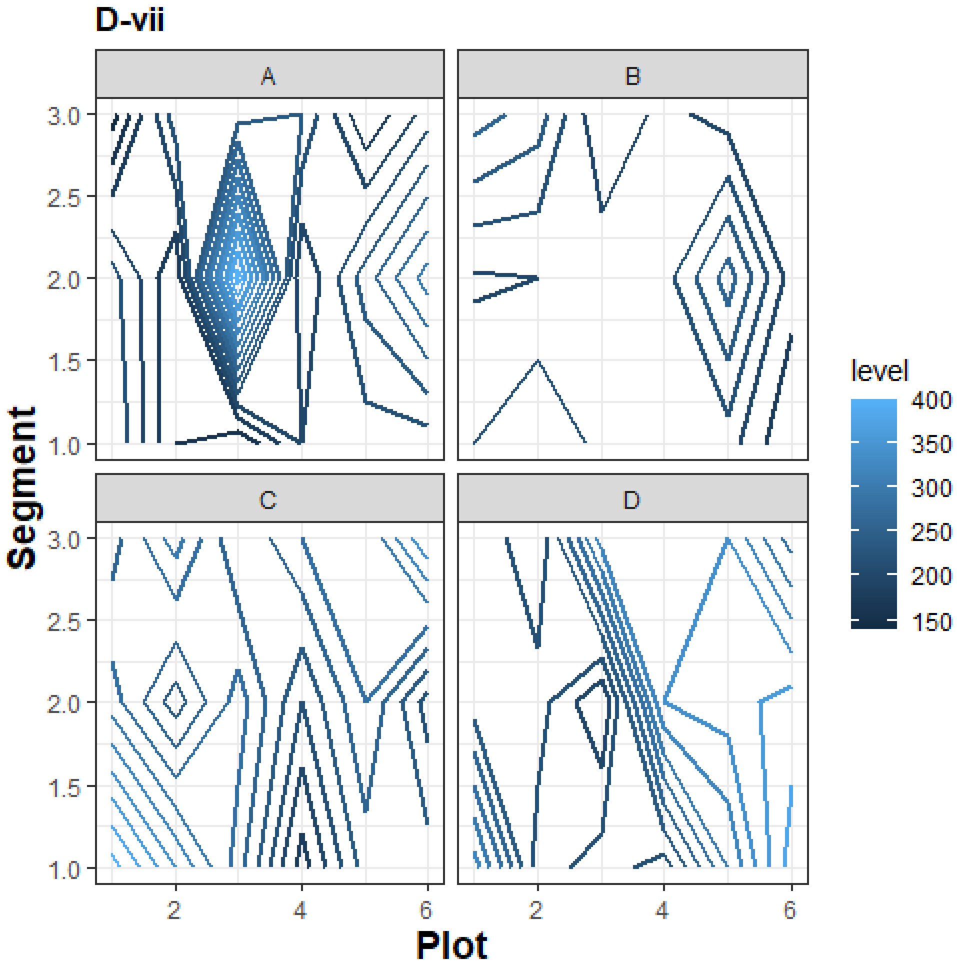

D-vii
A
B
C
D
Segment
Plot
level
400
350
300
250
200
150
3.0
2.5
2.0
1.5
1.0
2
4
6

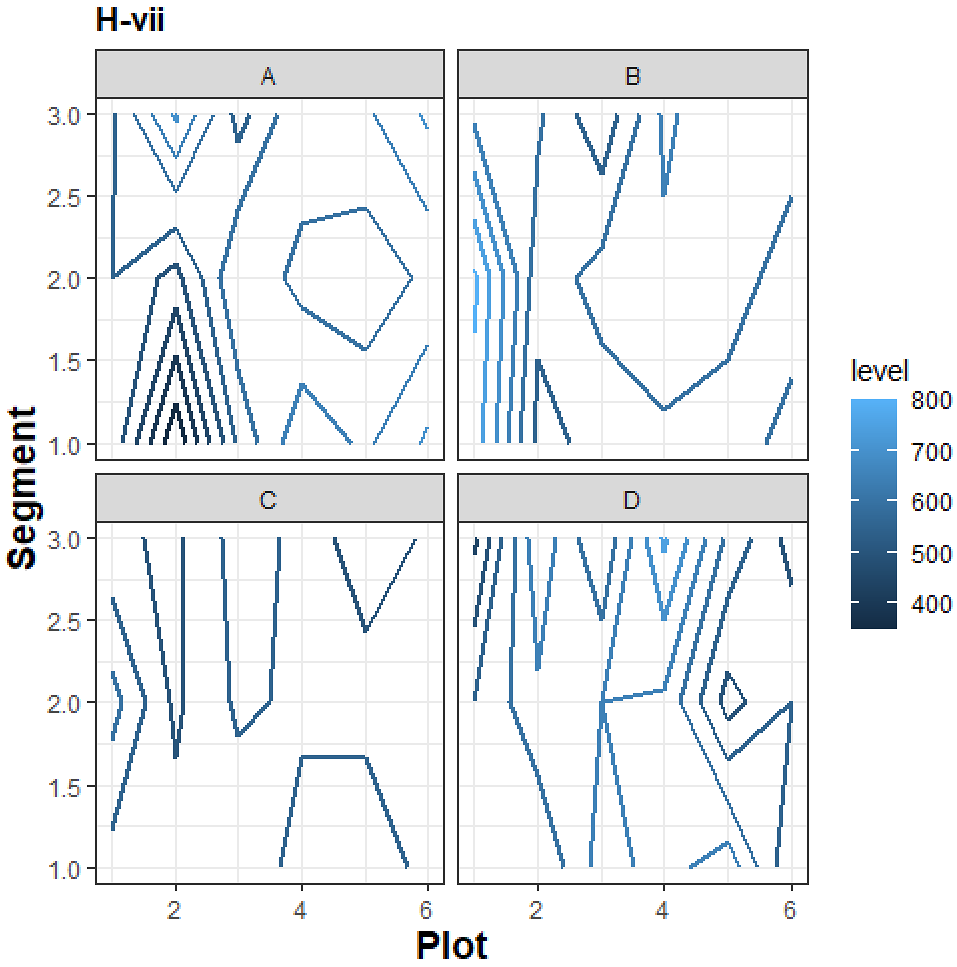

H-vii
A
B
C
D
3.0
2.5
2.0
1.5
1.0
Segment
2
4
6
Plot
level
800
700
600
500
400

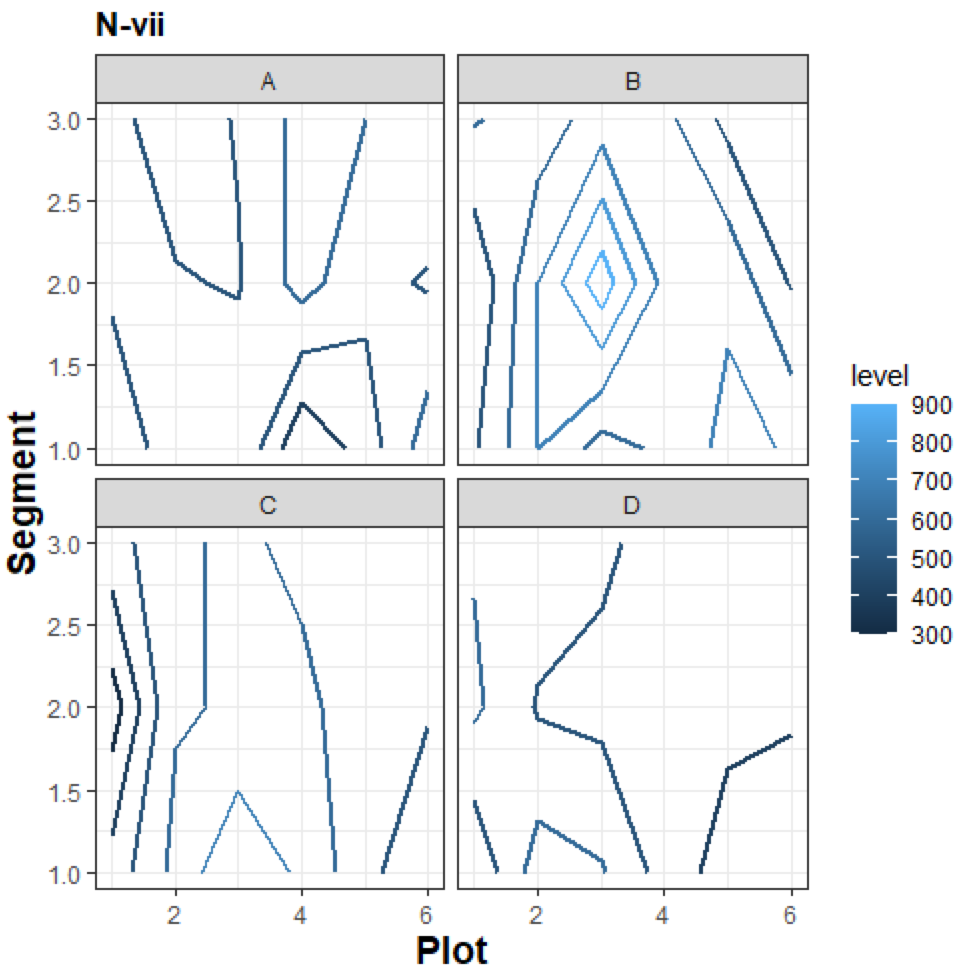
N-vii
A
B
C
D
Segment
Plot
level
900
800
700
600
500
400
300
3.0
2.5
2.0
1.5
1.0
2
4
6

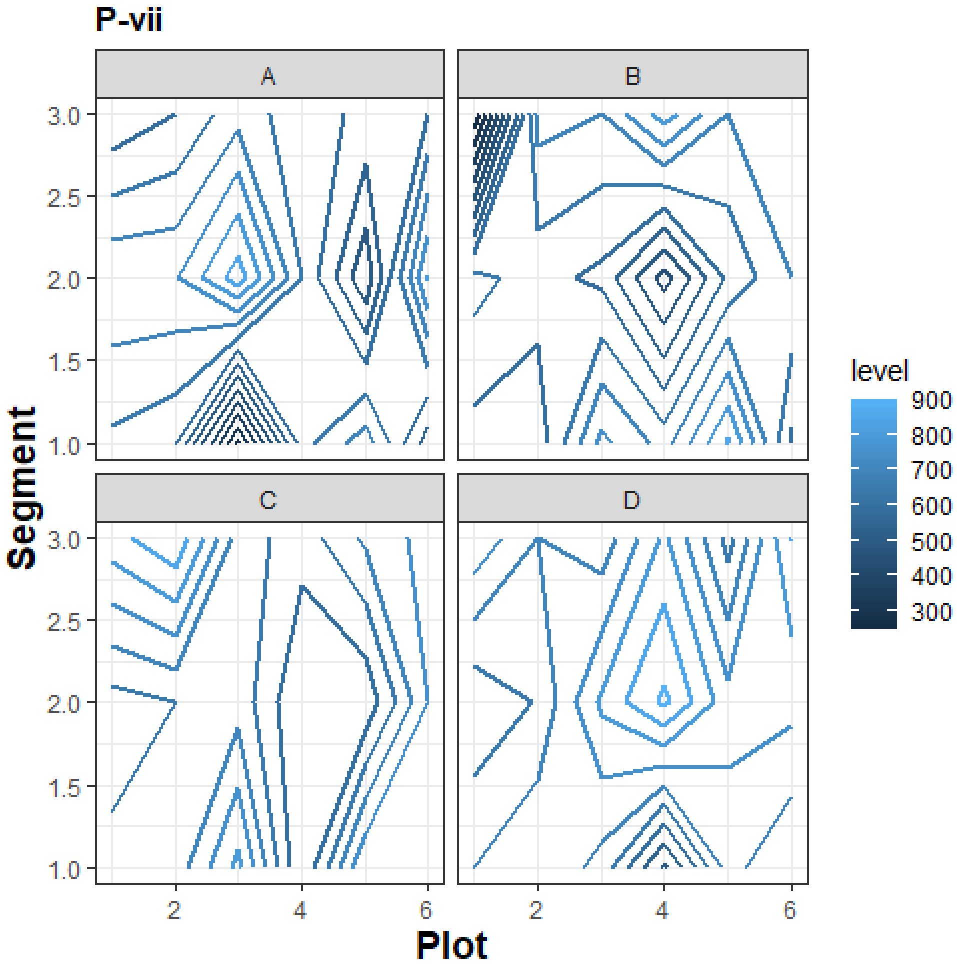
P-vii
A
B
C
D
Segment
Plot
level
900
800
700
600
500
400
300
3.0
2.5
2.0
1.5
1.0
2
4
6

D-viii

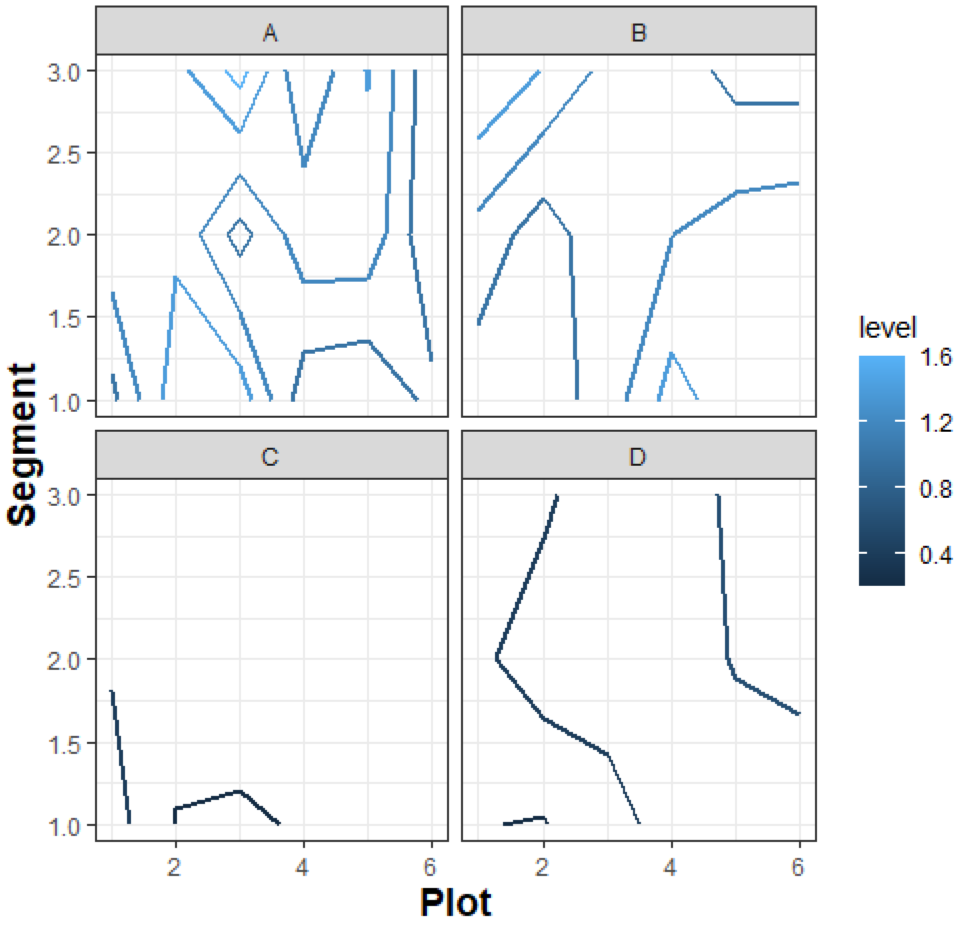

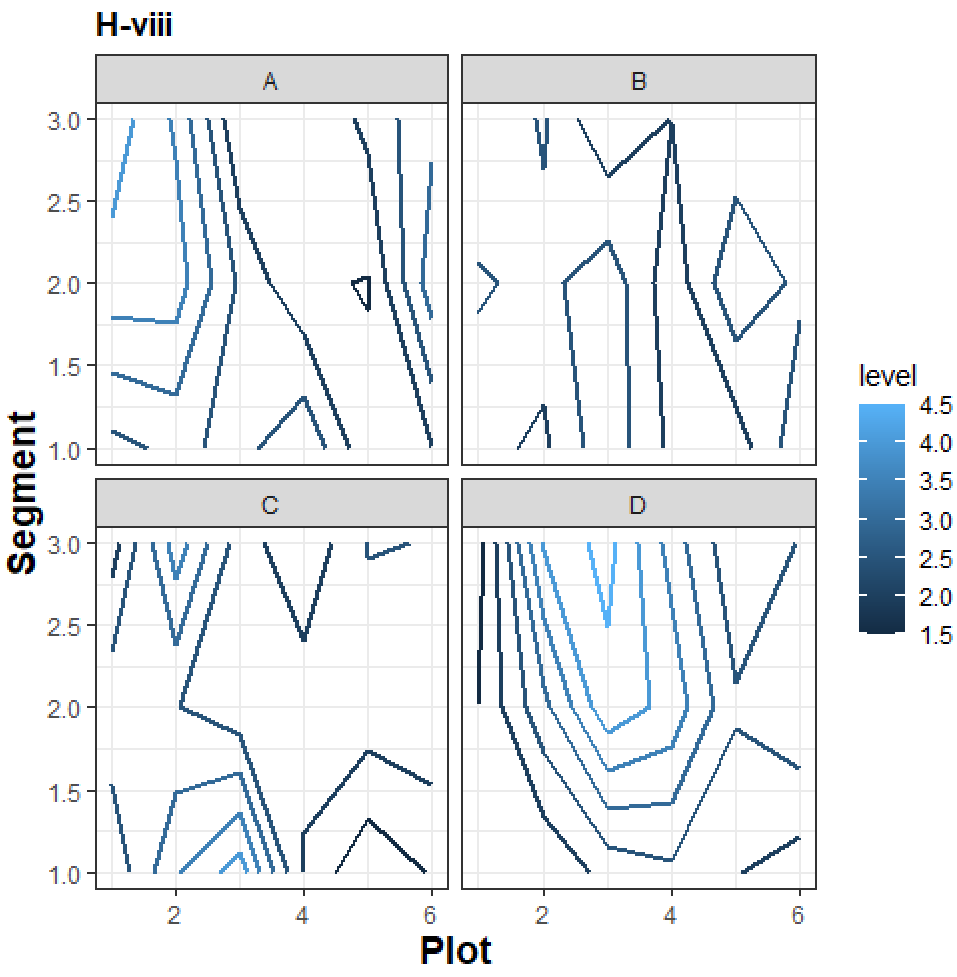
H-viii
A
B
C
D
Segment
Plot
level
4.5
4.0
3.5
3.0
2.5
2.0
1.5
3.0
2.5
2.0
1.5
1.0
2
4
6

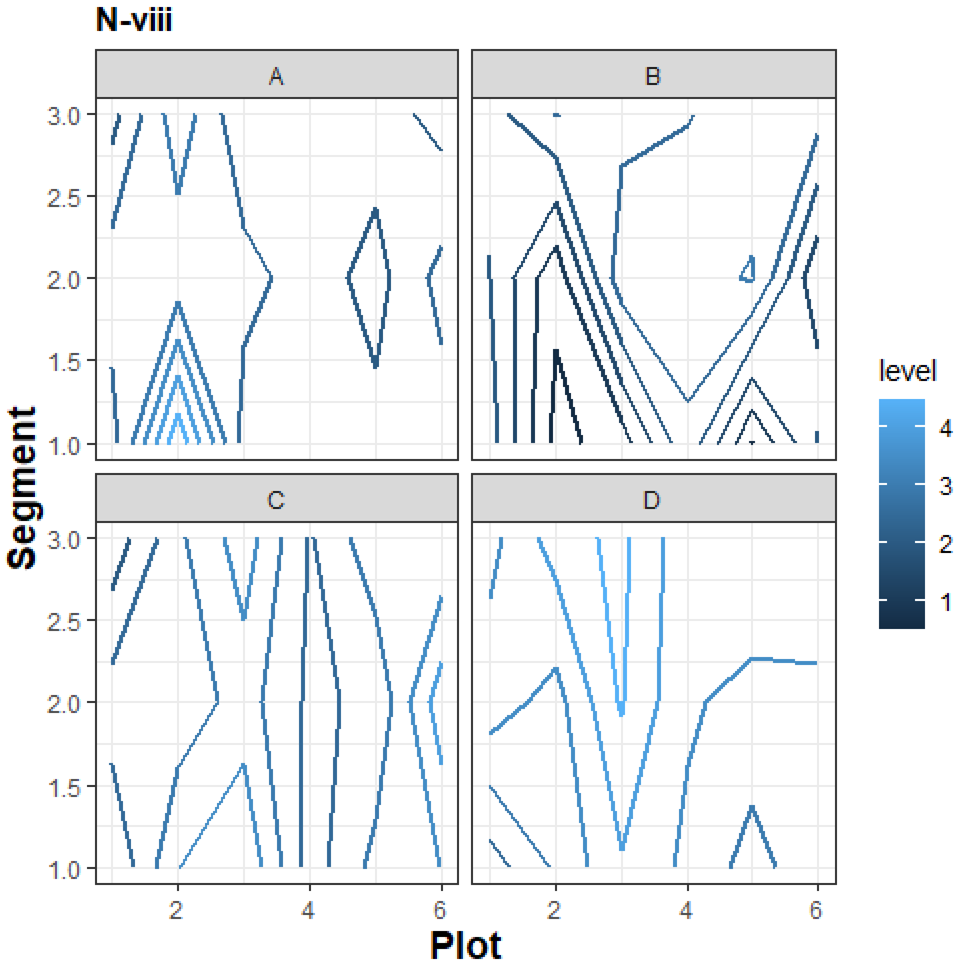

N-viii
A
B
C
D
3.0
2.5
2.0
1.5
1.0
Segment
2
4
6
Plot
level
4
3
2
1

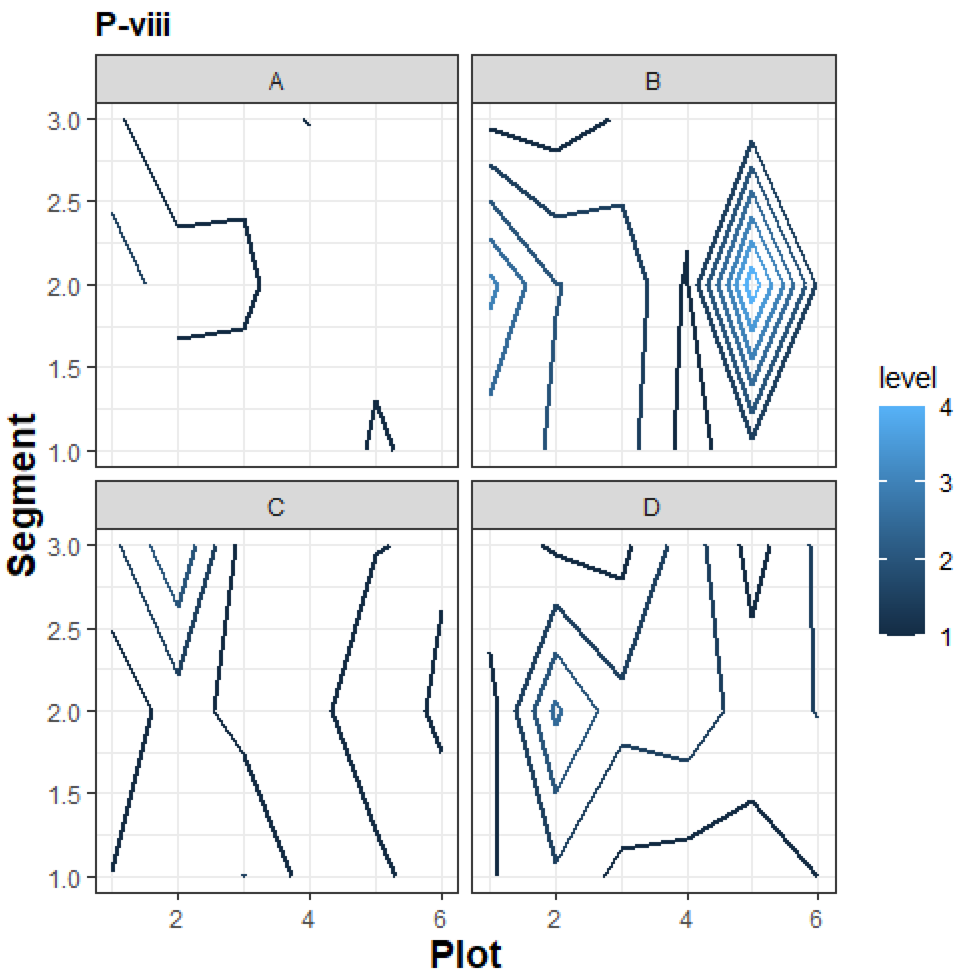
P-viii
A
B
C
D
Segment
Plot
level
4
3
2
1
3.0
2.5
2.0
1.5
1.0
2
4
6

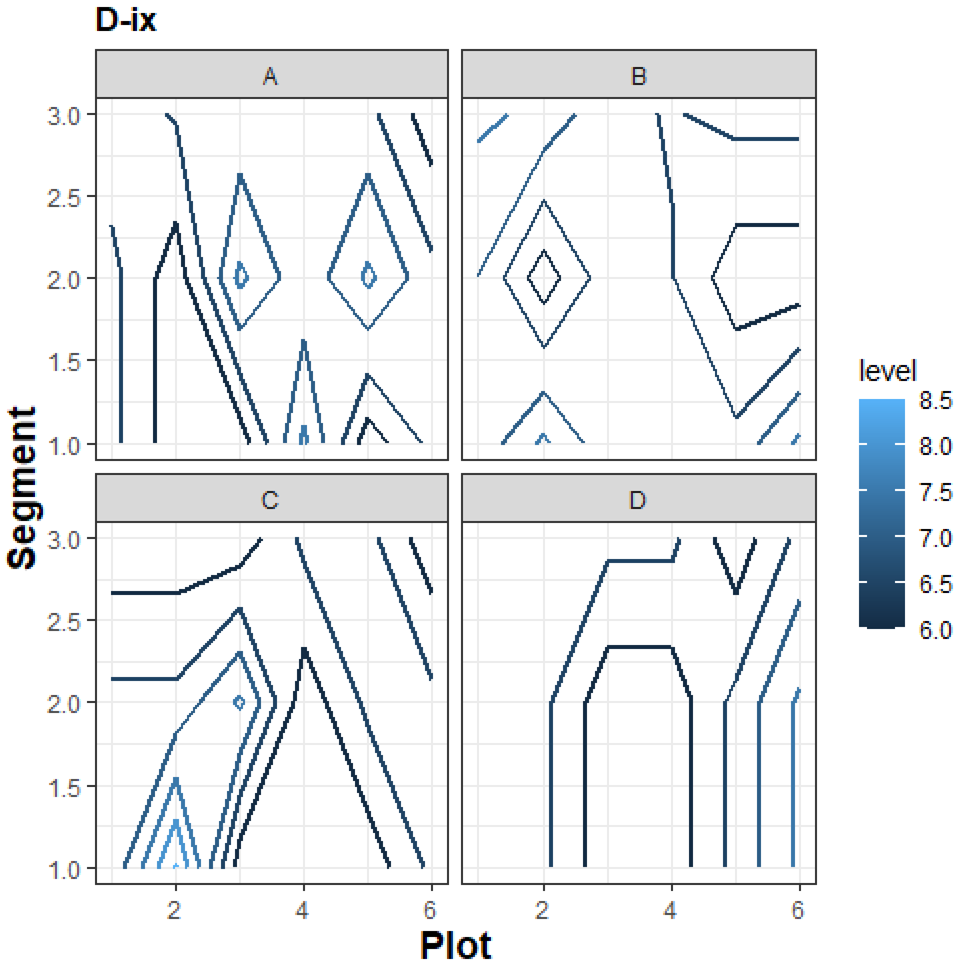

D-ix
A
B
C
D
3.0
2.5
2.0
1.5
1.0
Segment
2
4
6
Plot
level
8.5
8.0
7.5
7.0
6.5
6.0

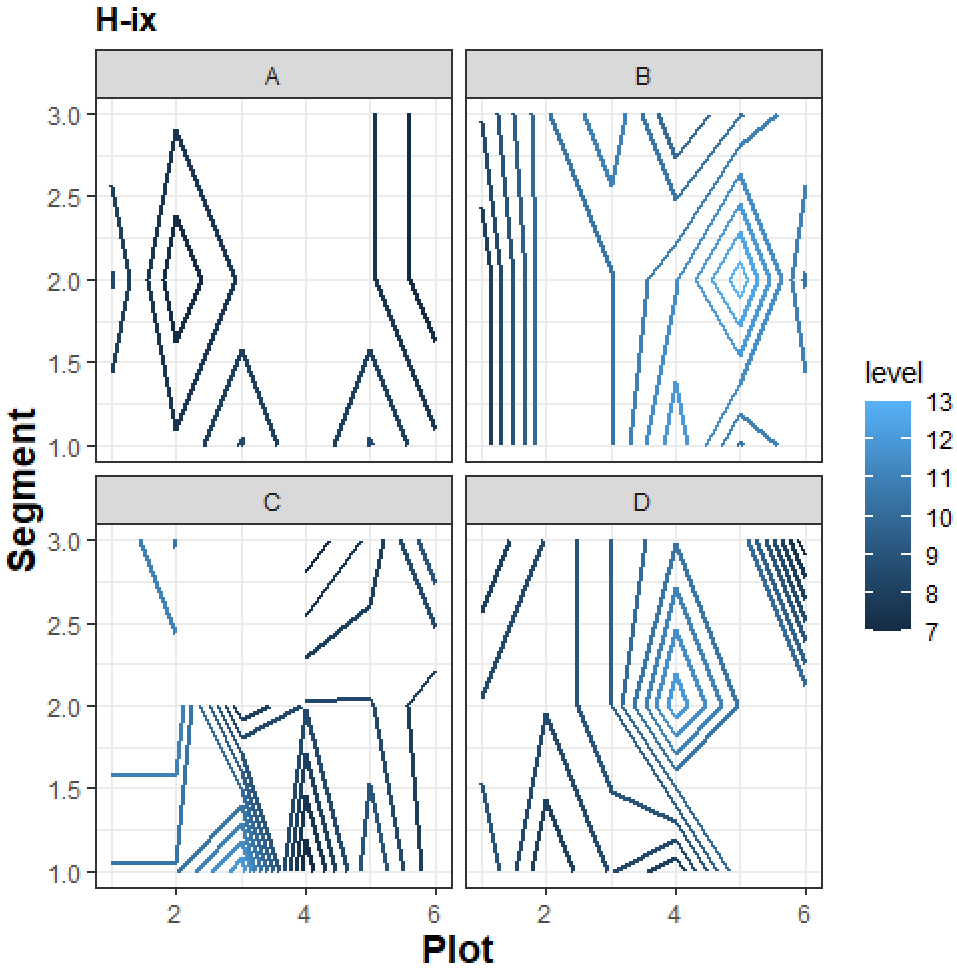
H-ix
A
B
C
D
3.0
2.5
2.0
1.5
1.0
Segment
2
4
6
Plot
level
13
12
11
10
9
8
7

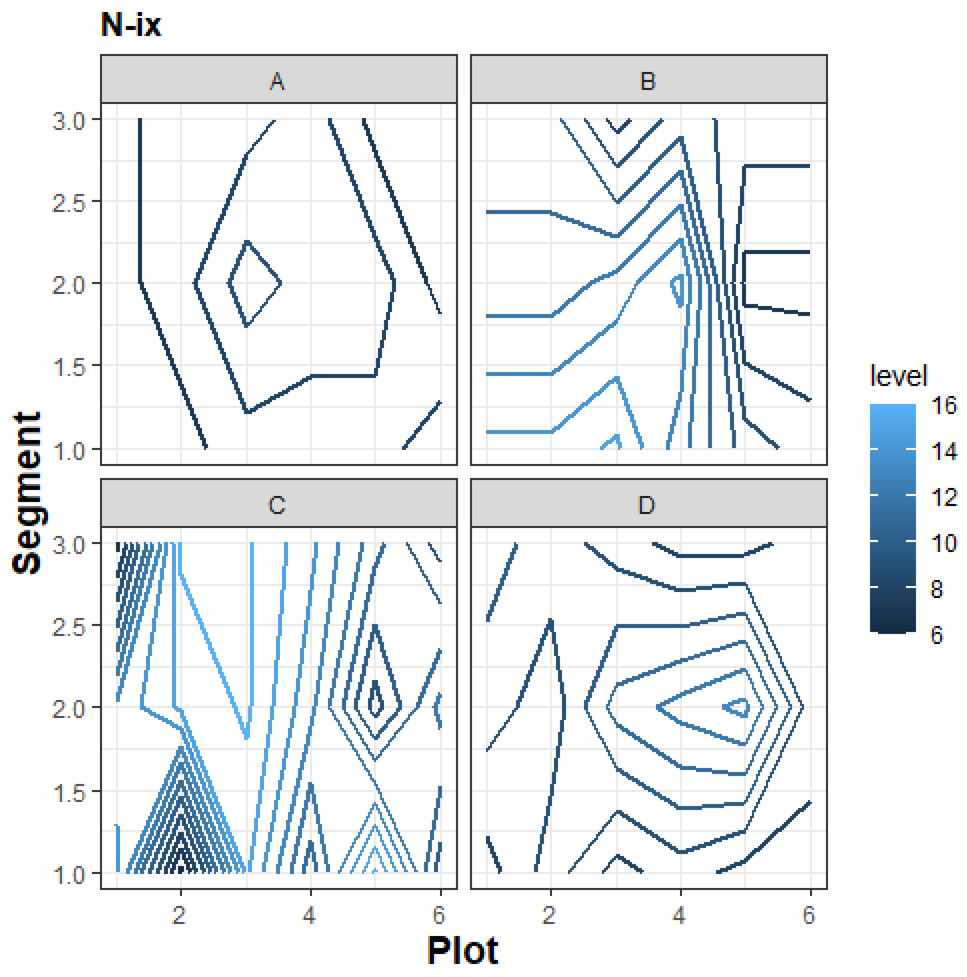
N-ix
A
B
C
D
Segment
Plot
level
16
14
12
10
8
6
3.0
2.5
2.0
1.5
1.0
2
4
6

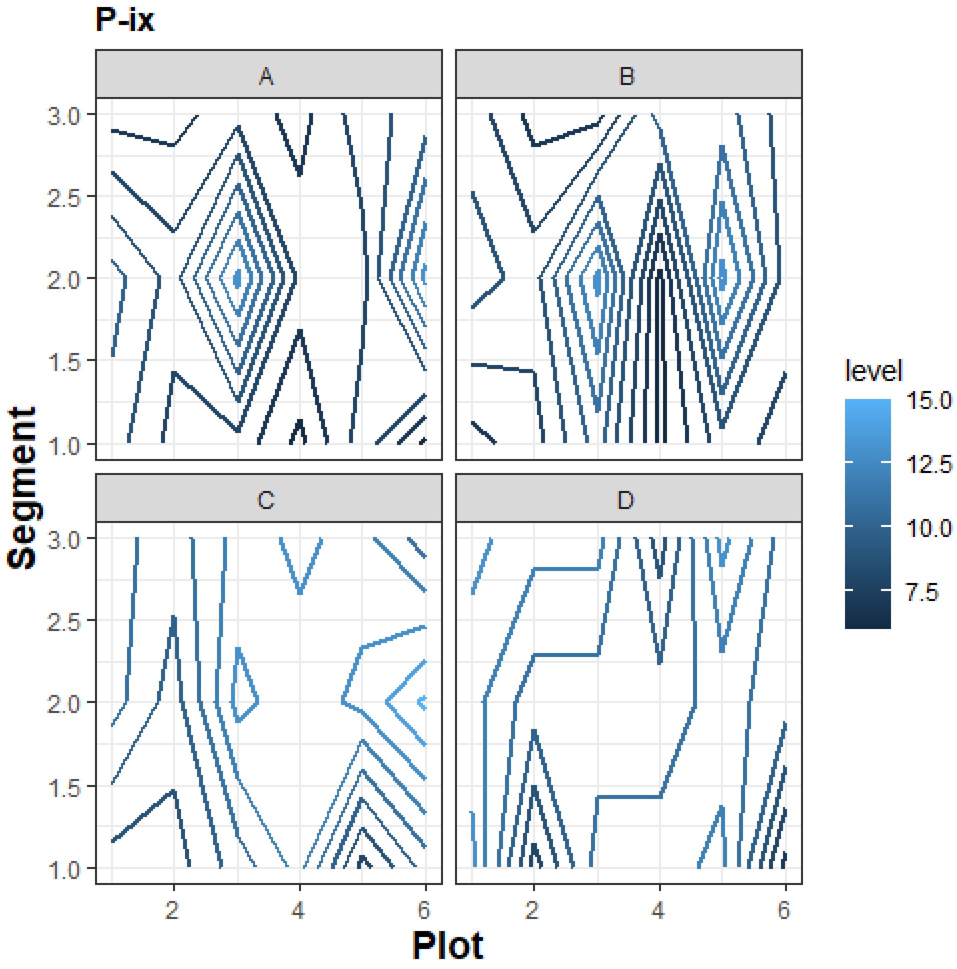
P-ix
A
B
C
D
3.0
2.5
2.0
1.5
1.0
Segment
2
4
6
Plot
level
15.0
12.5
10.0
7.5

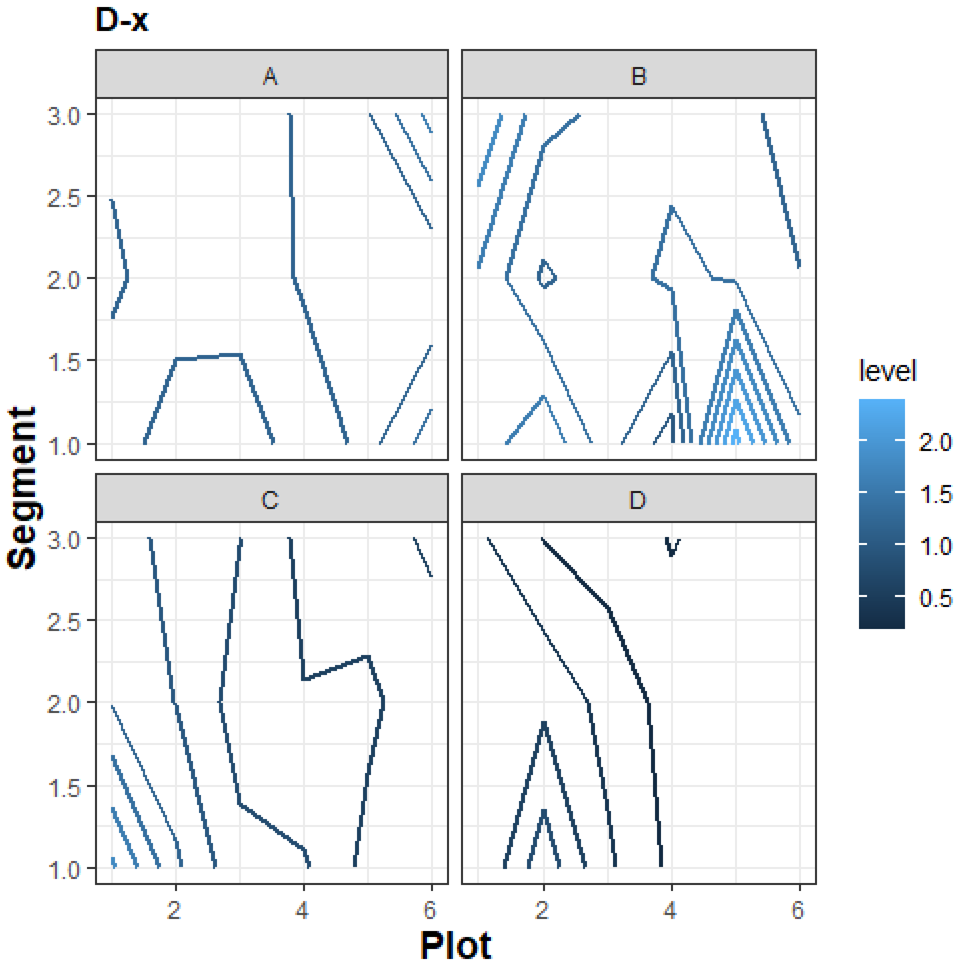

D-x
A
B
C
D
3.0
2.5
2.0
1.5
1.0
Segment
2
4
6
Plot
level
2.0
1.5
1.0
0.5

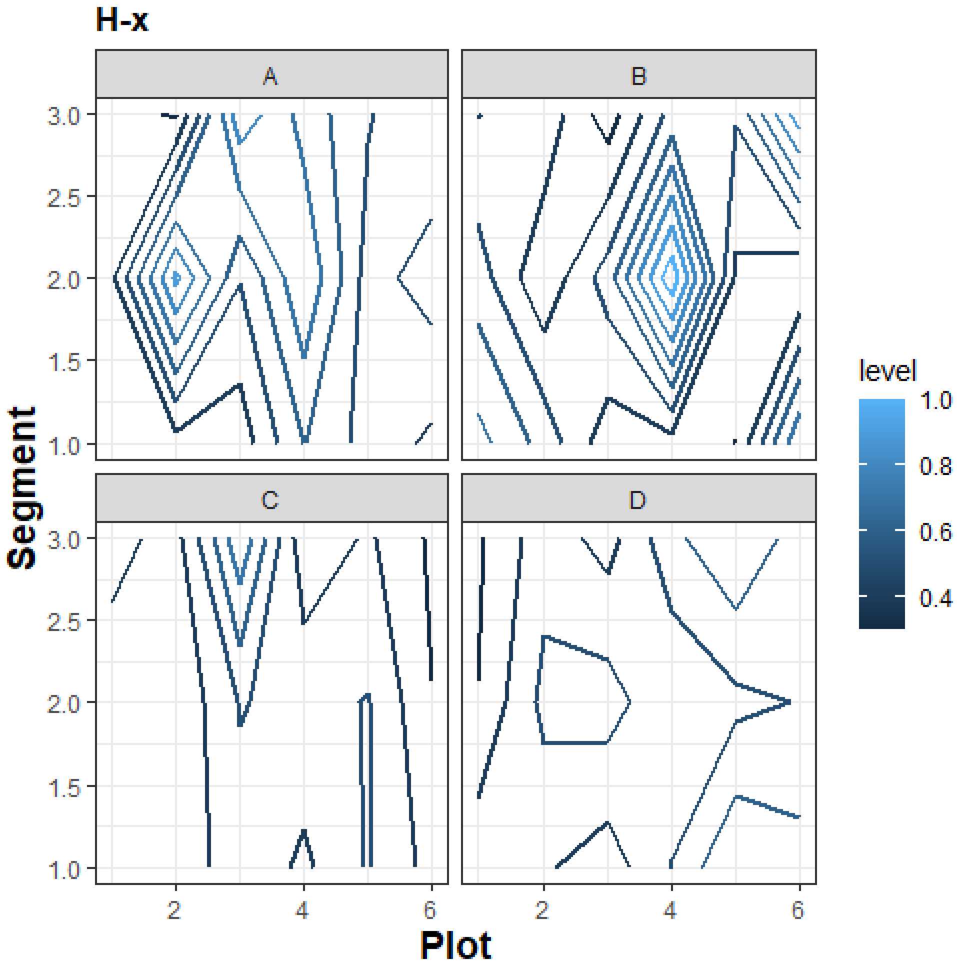

H-x
A
B
C
D
Segment
Plot
level
1.0
0.8
0.6
0.4
3.0
2.5
2.0
1.5
1.0
2
4
6

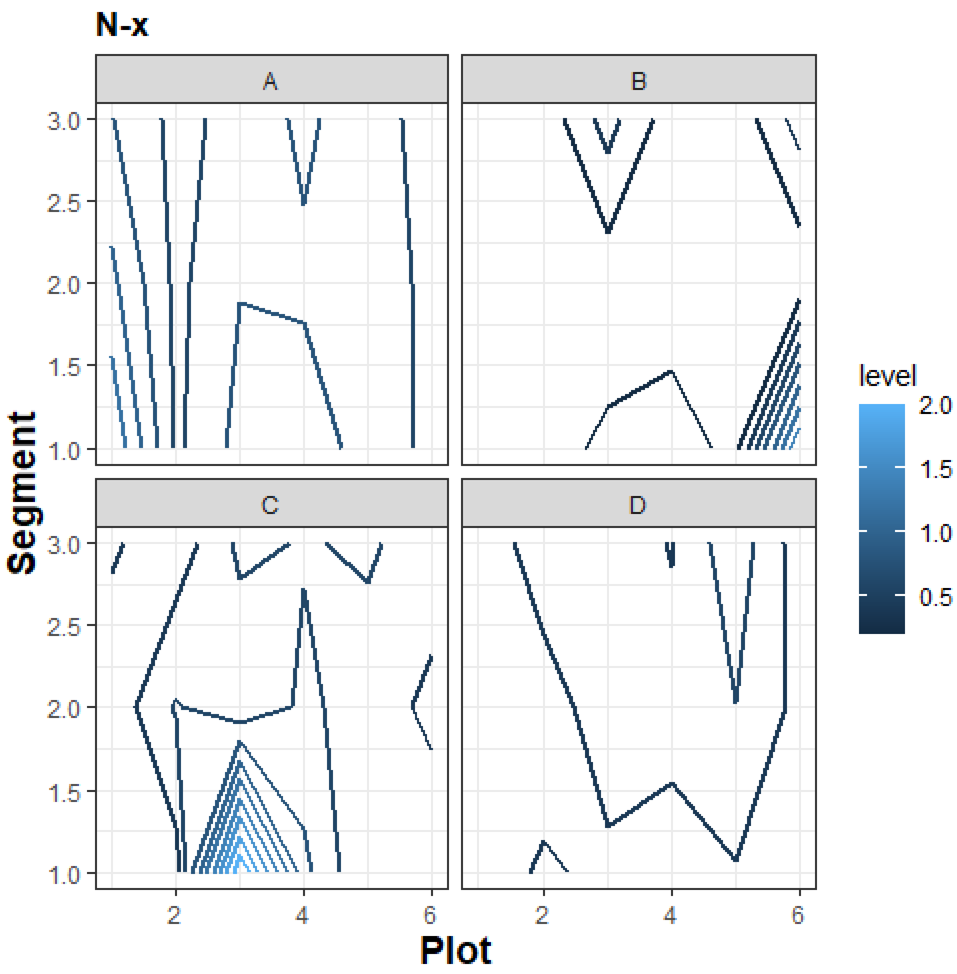

N-x
A
B
C
D
Segment
Plot
level
2.0
1.5
1.0
0.5
3.0
2.5
2
4
6

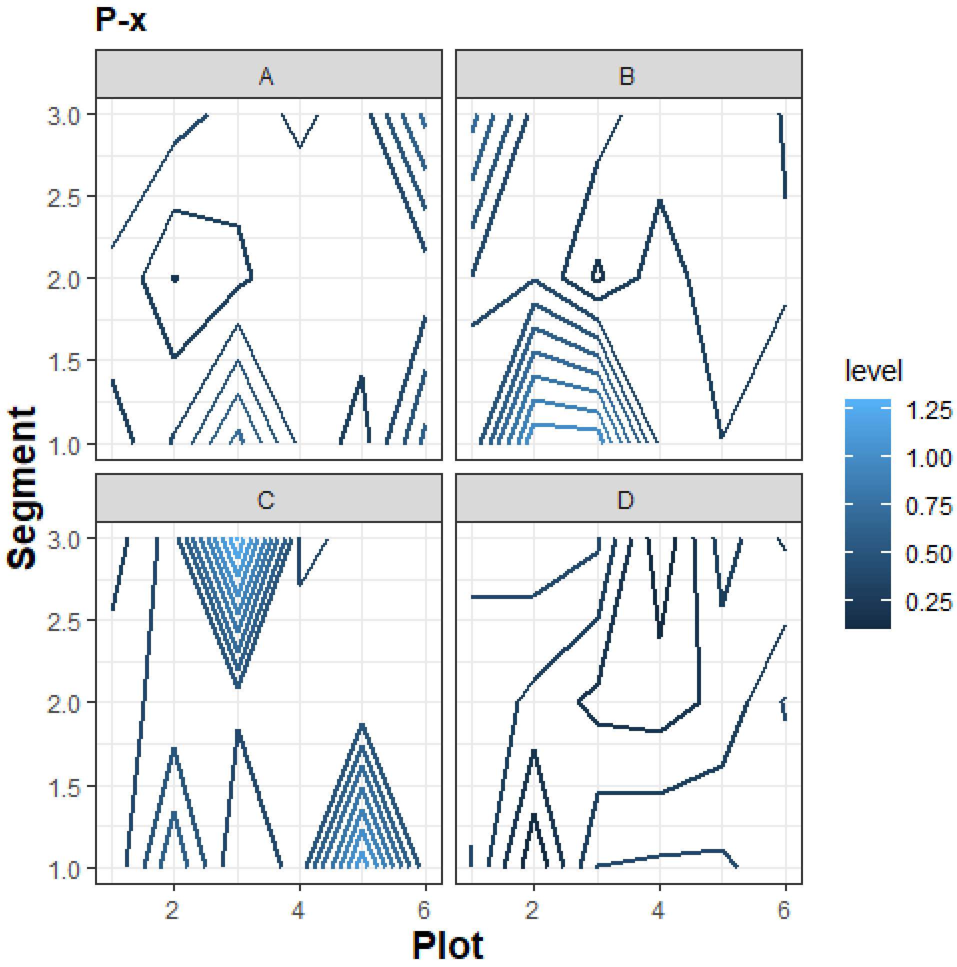

P-x
A
B
C
D
3.0
2.5
2.0
1.5
1.0
Segment
2
4
6
Plot
level
1.25
1.00
0.75
0.50
0.25

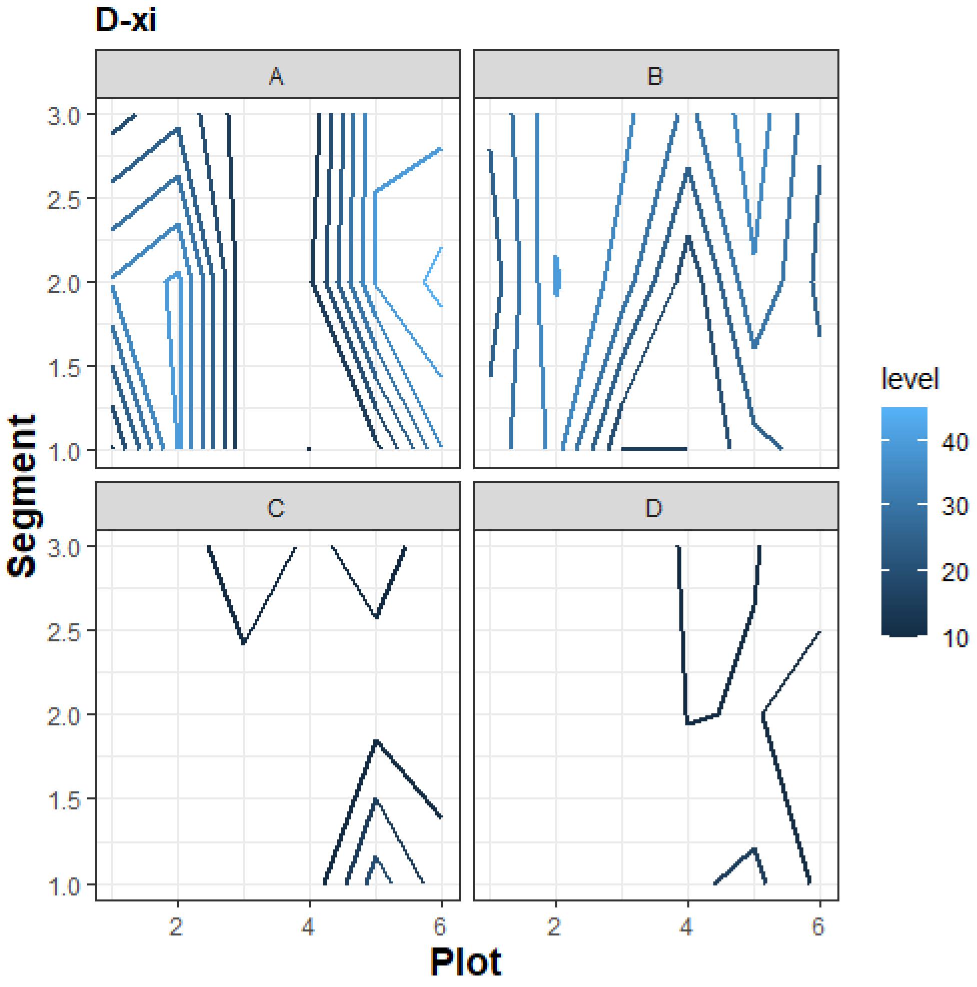
D-xi
A
B
C
D
3.0
2.5
2.0
1.5
1.0
Segment
2
4
6
Plot
level
40
30
20
10

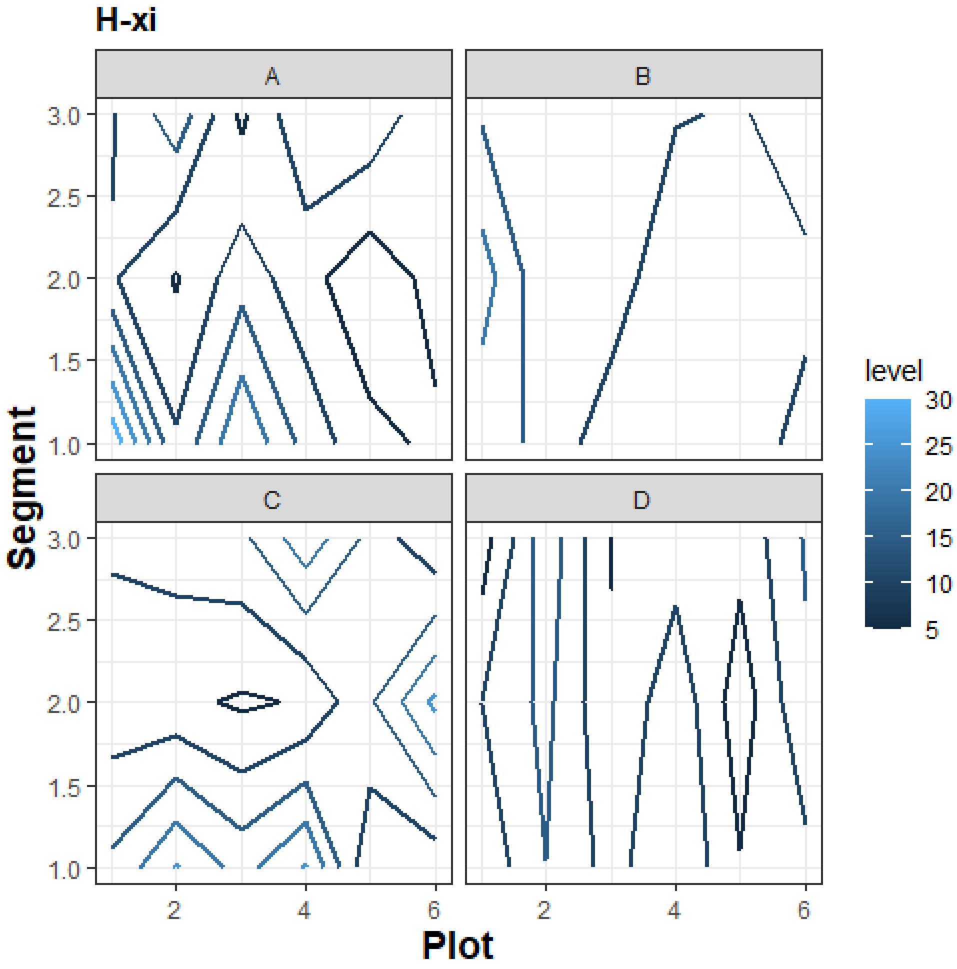

H-xi
A
B
C
D
Segment
3.0
2.5
2.0
1.5
1.0
Plot
2
4
6
level
30
25
20
15
10
5

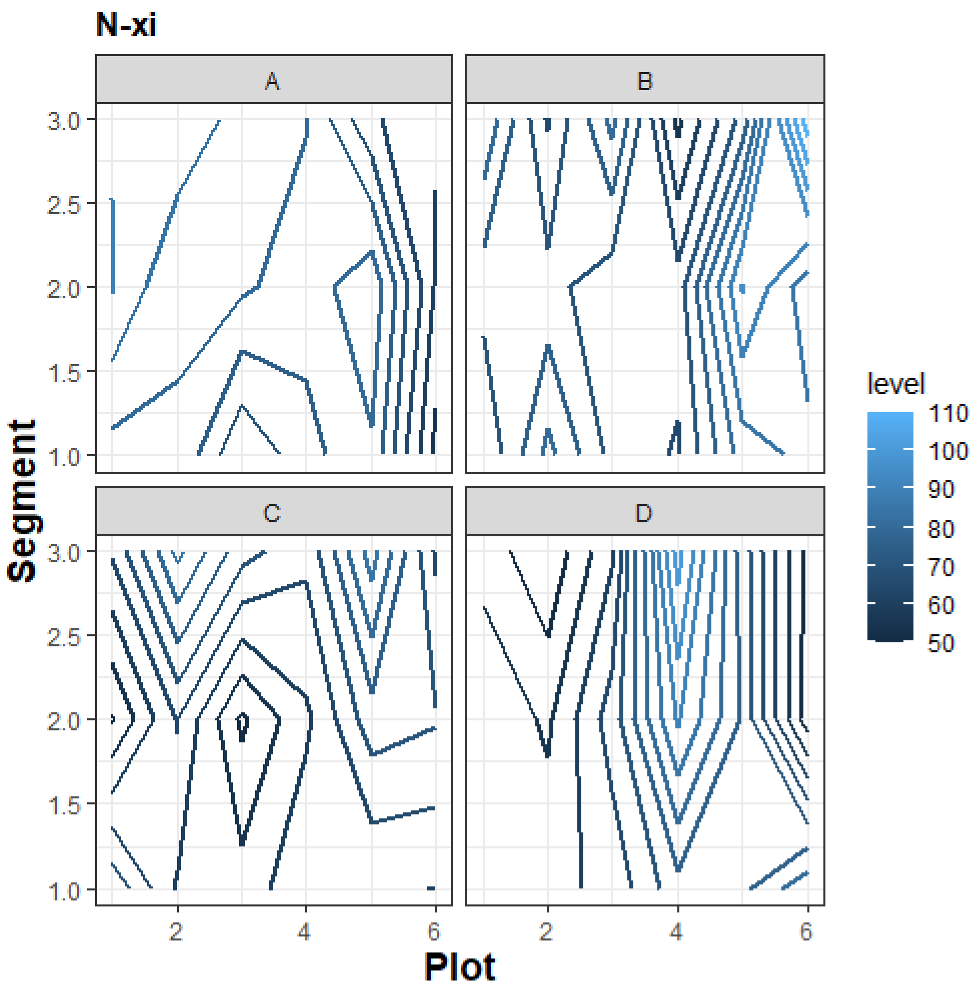
N-xi
A
B
C
D
Segment
3.0
2.5
2.0
1.5
1.0
2
4
6
Plot
level
110
100
90
80
70
60
50

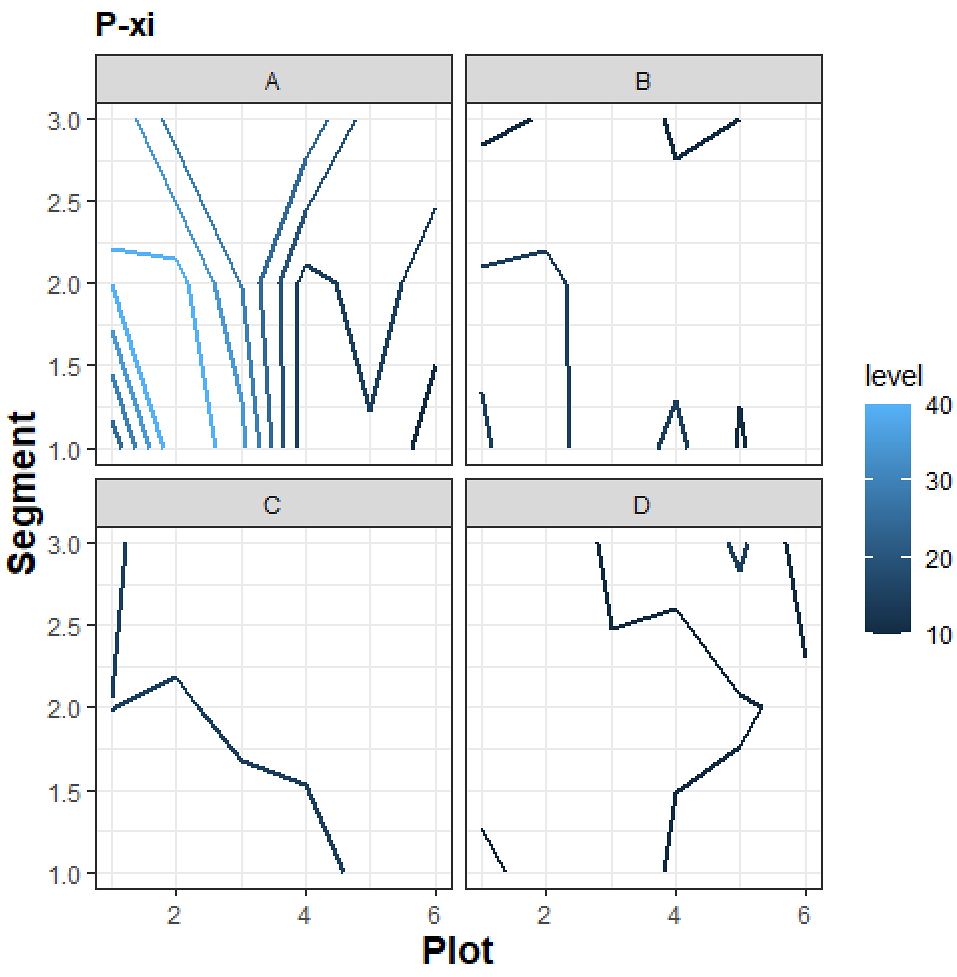
P-xi
A
B
C
D
level
40
30
20
10
3.0
2.5
2.0
1.5
1.0
2
4
6
Segment
Plot

D-xii

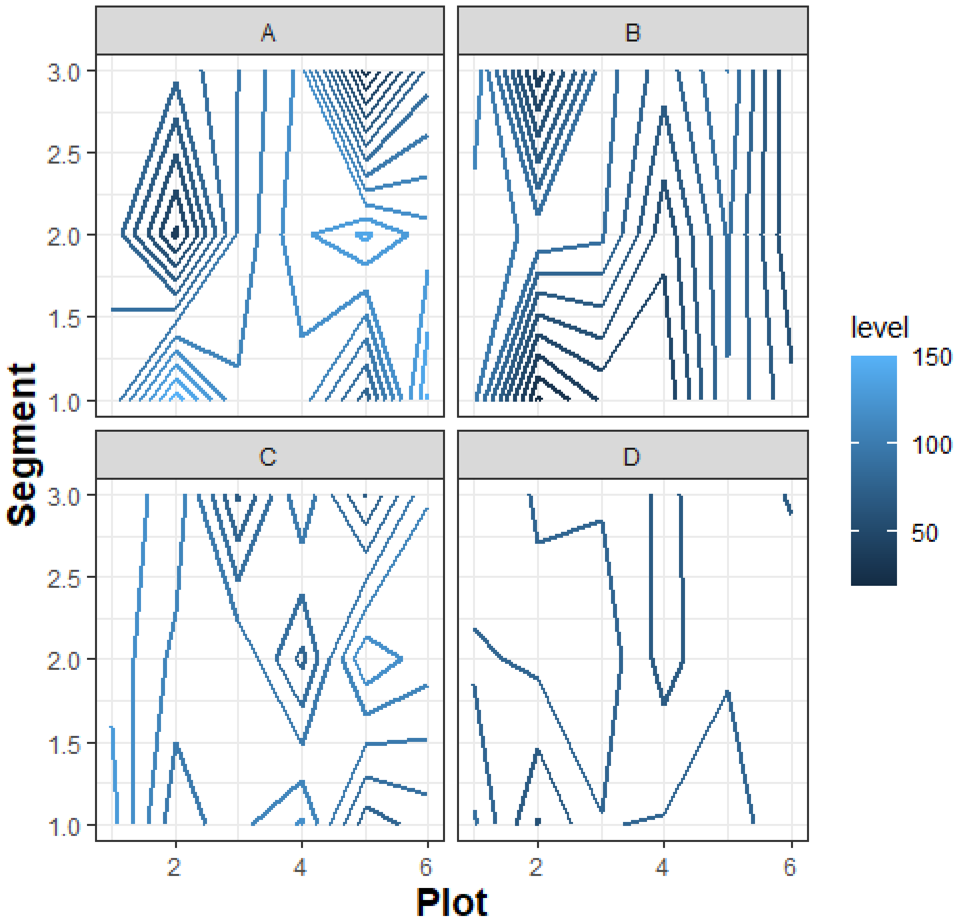

H-xii

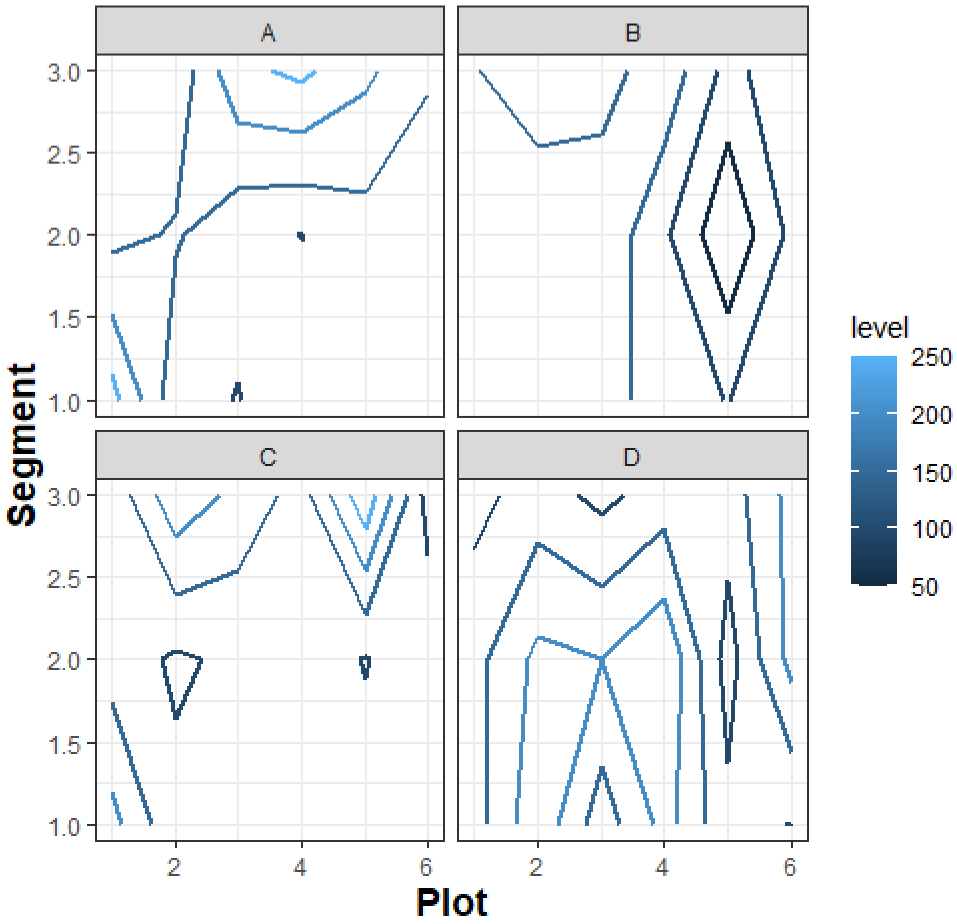

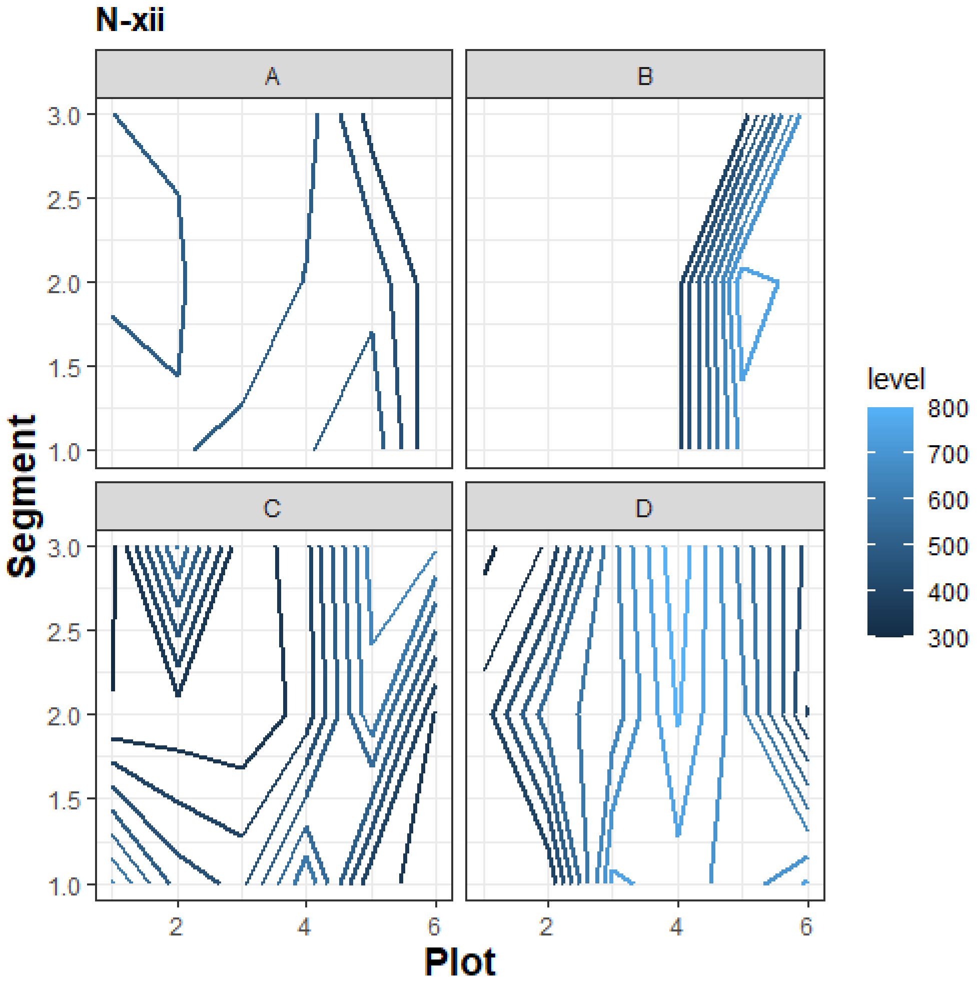
N-xii
A
B
C
D
Segment
Plot
level
800
700
600
500
400
300
3.0
2.5
2.0
1.5
1.0
2
4
6

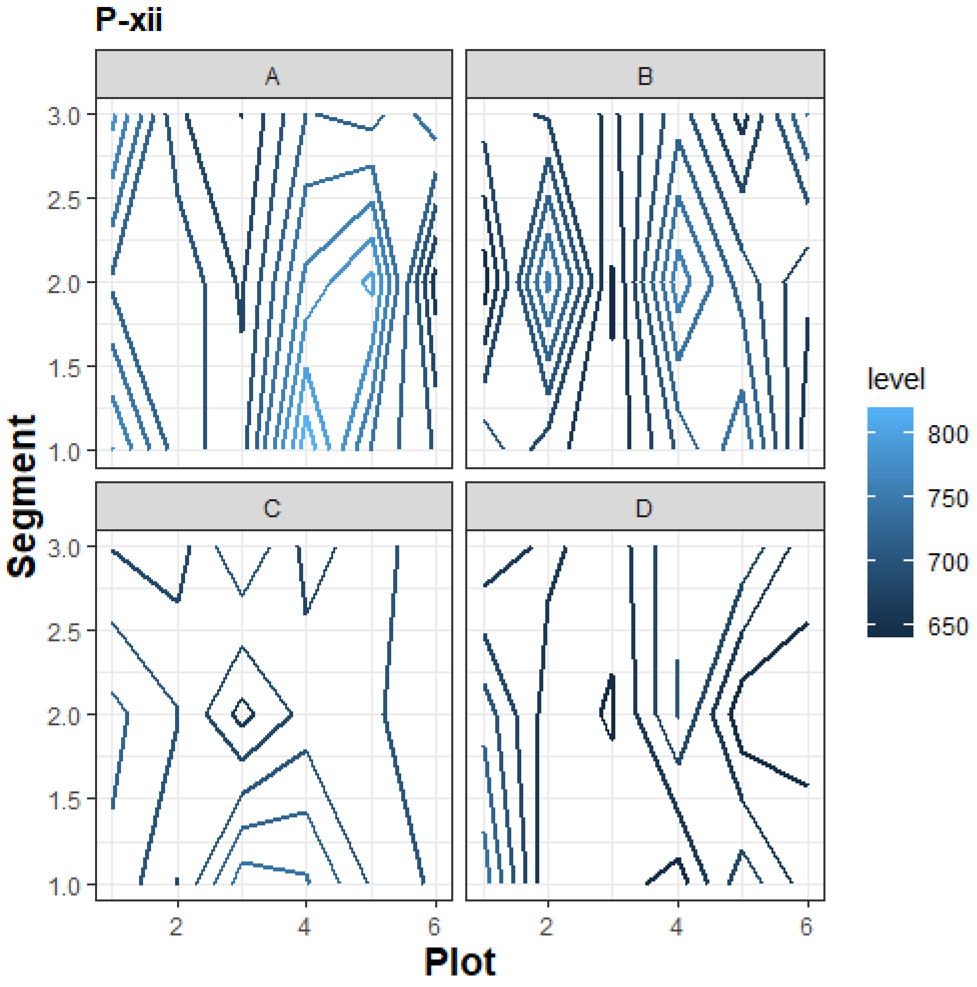
P-xii
A
B
C
D
Segment
3.0
2.5
2.0
1.5
1.0
2
4
6
Plot
level
800
750
700
650

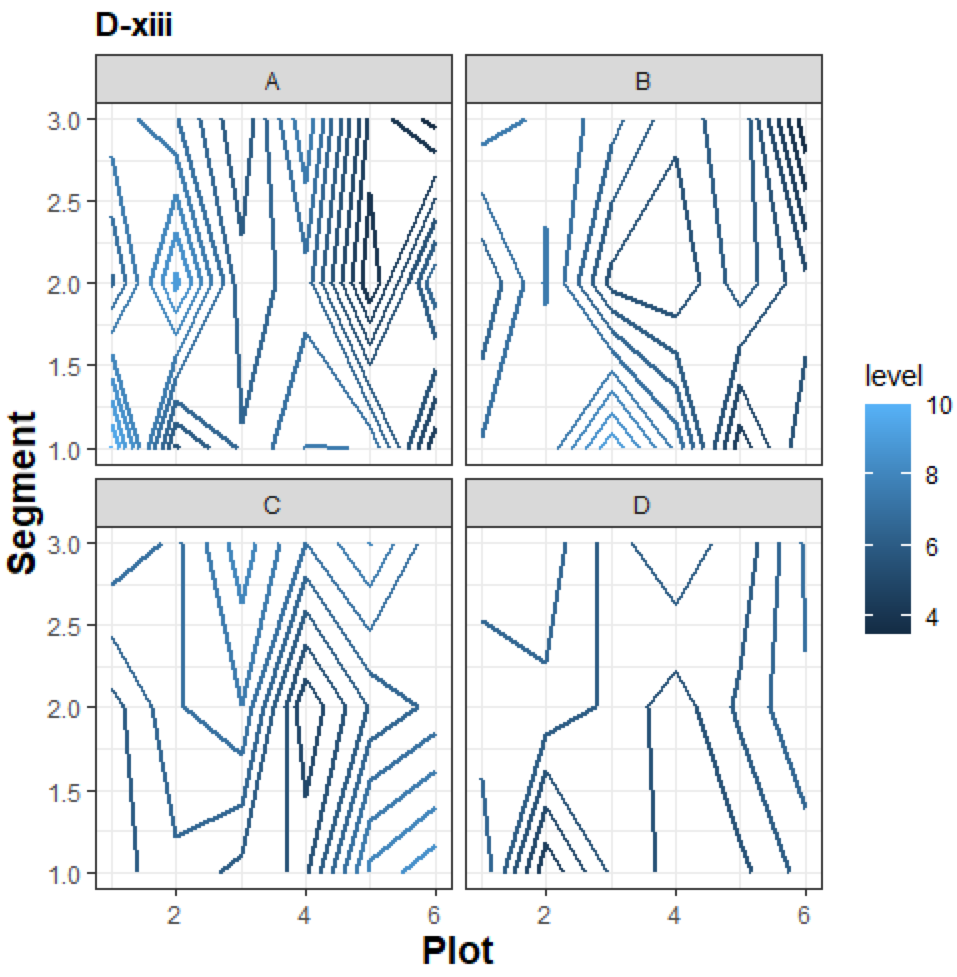
D-xiii
A
B
C
D
Segment
3.0
2.5
2.0
1.5
1.0
Plot
2
4
6
level
10
8
6
4

H-xiii

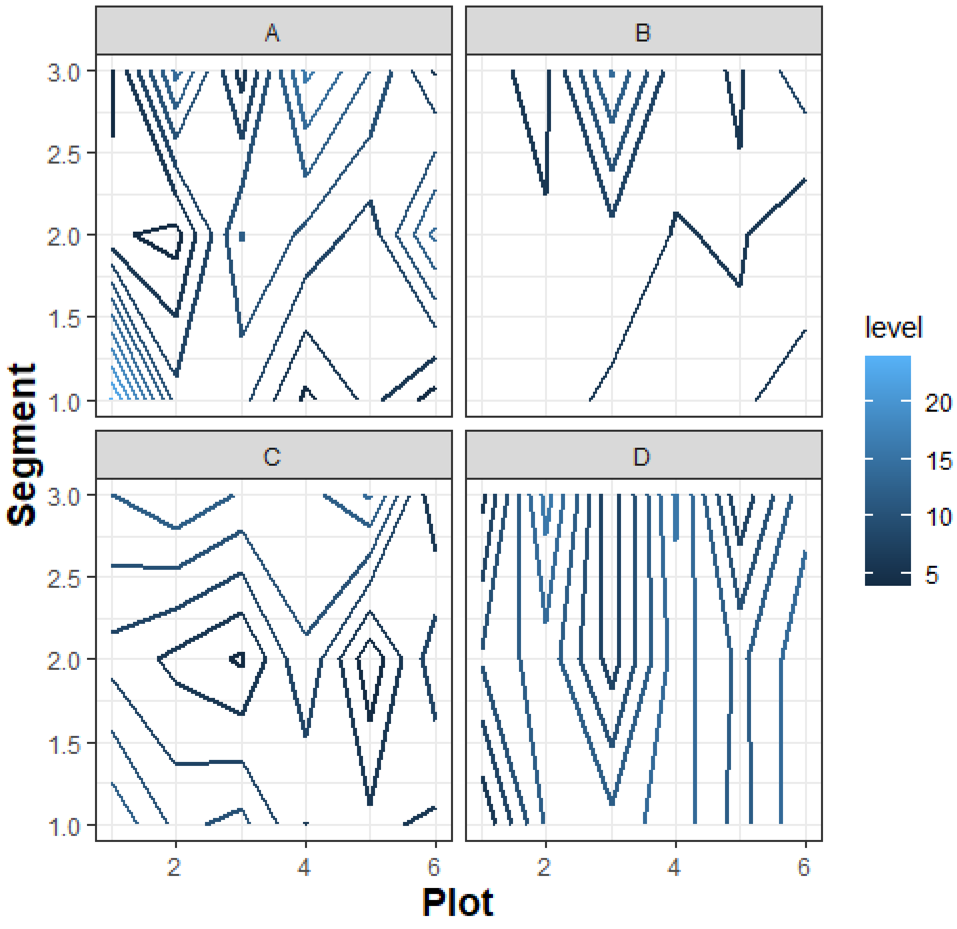

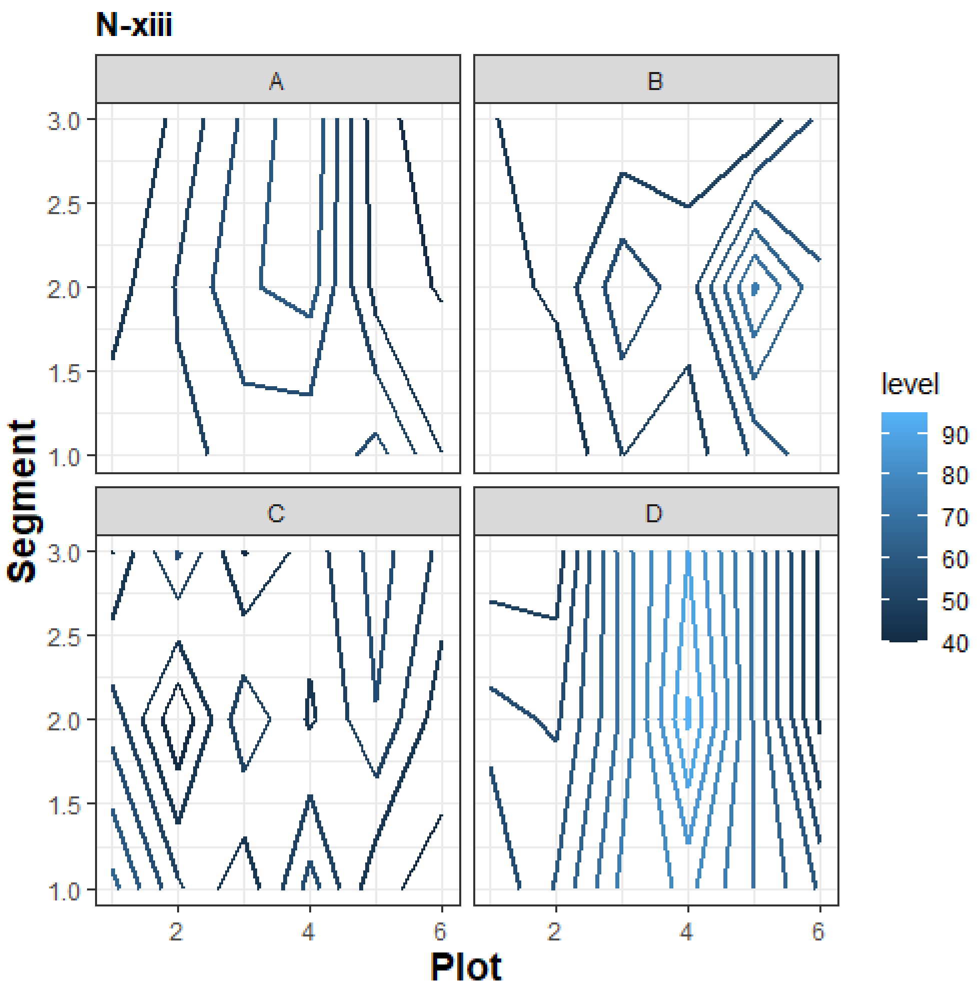
N-xiii
A
B
C
D
Segment
Plot
3.0
2.5
2.0
1.5
1.0
2
4
6
level
90
80
70
60
50
40

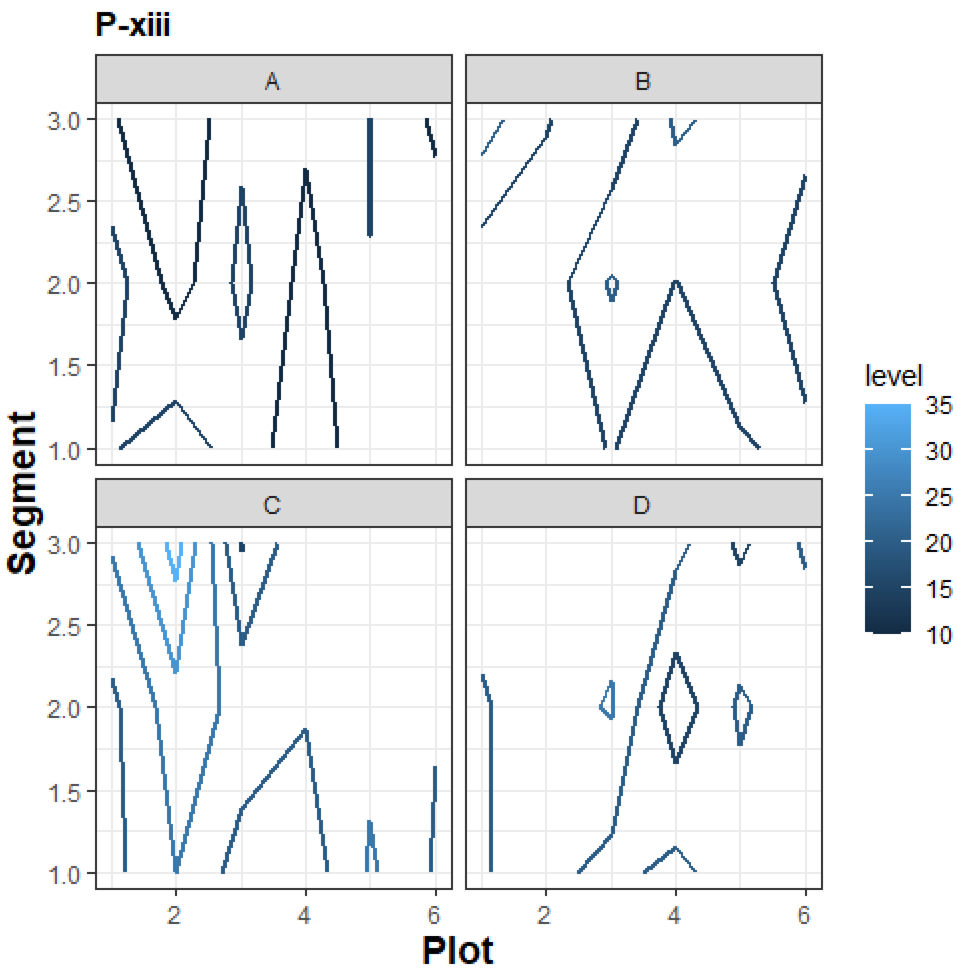


**Figure S2:** Variation in soil features within blocks A, B, C, and D in a location. The letters D, H, N, and P stand for locations Delhi, Haveri, Nagpur, and Pune respectively. There are 13 soil features measured: i) gravimetric moisture content (%), ii) pH, iii) salinity (ppm), iv) sand (%), v) clay (%), vi) silt (%), vii) carbon (mg/100g), viii) phosphorous (mg/100g), ix) nitrogen (mg/100g), x) sulphur (mg/100g), xi) sodium (mg/100g), xii) calcium (mg/100g), and xiii) potassium (mg/100g).

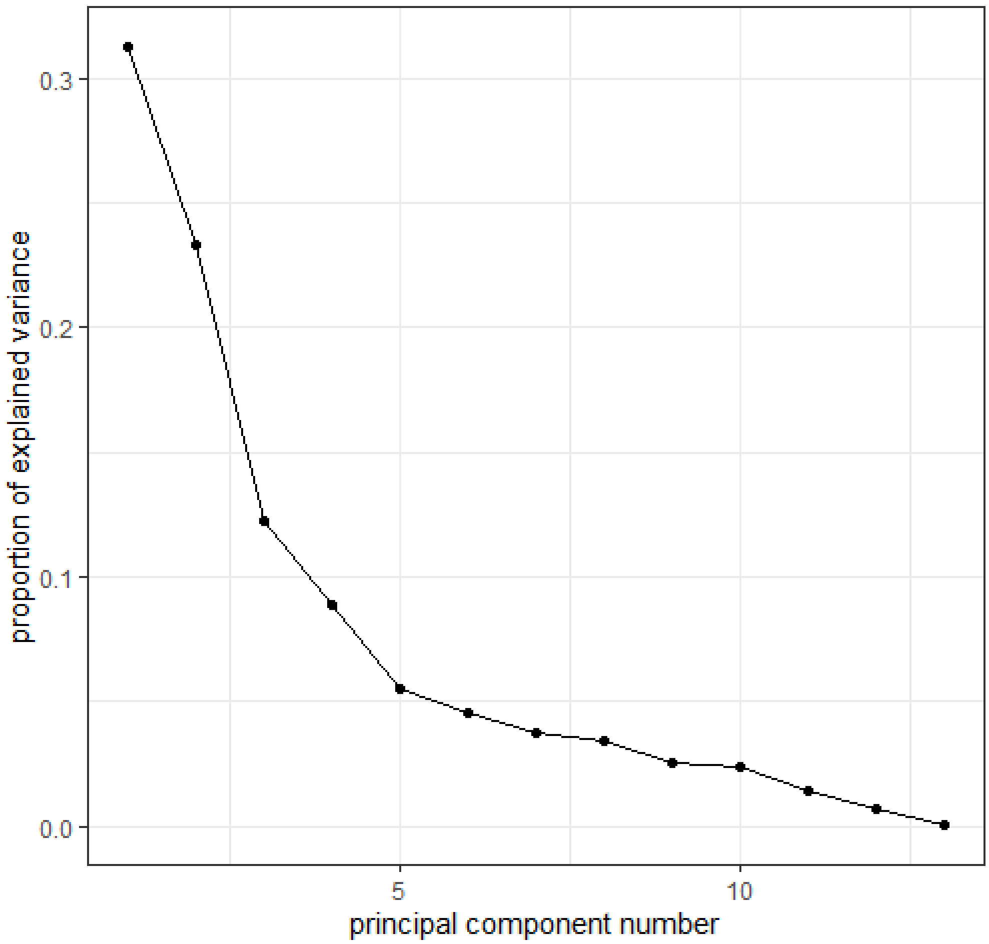


**Figure S3:** Scree plot from principal component analysis (PCA) indicating the proportion of variation explained by each principal component. Together PC 1 and 2 explained 54% of the variation in the data.

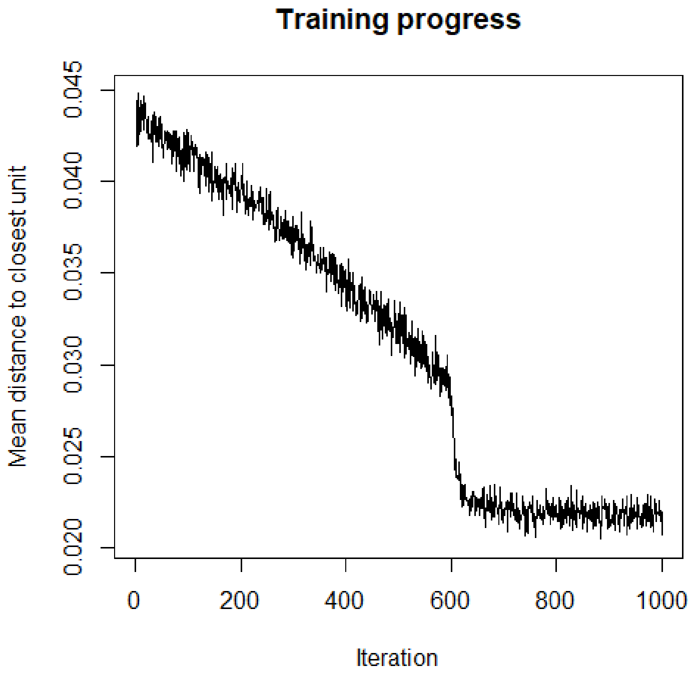


**Figure S4:** Convergence of SOM occurred at around 600 iterations.
